# A cytoplasmic Argonaute protein promotes the inheritance of RNAi

**DOI:** 10.1101/235713

**Authors:** Fei Xu, Xuezhu Feng, Xiangyang Chen, Chenchun Weng, Qi Yan, Ting Xu, Minjie Hong, Shouhong Guang

## Abstract

RNAi-elicited gene silencing is heritable and can persist for multiple generations after its initial induction in *C. elegans*. However, the mechanism by which parental-acquired trait-specific information from RNAi is inherited by the progenies is not fully understood. Here, we identified a cytoplasmic Argonaute protein, WAGO-4, necessary for the inheritance of RNAi. WAGO-4 exhibits asymmetrical translocation to the germline during early embryogenesis, accumulates at the perinuclear foci in the germline, and is required for the inheritance of exogenous RNAi targeting both germline- and soma-expressed genes. WAGO-4 binds to 22G-RNAs and their mRNA targets. Interestingly, WAGO-4-associated endogenous 22G-RNAs target the same cohort of germline genes as CSR-1 and contain untemplated addition of uracil at the 3’ ends. The poly(U) polymerase CDE-1 is required for the untemplated uridylation of 22G-RNAs and inheritance of RNAi. Therefore, we conclude that, in addition to the nuclear RNAi pathway, the cytoplasmic RNAi machinery also promotes RNAi inheritance.

## Introduction

RNAi-elicited gene silencing is heritable and can perpetuate for a number of generations in *C. elegans* (reviewed in (Heard and Martienssen, 2014; Miska and Ferguson-Smith, 2016; Rechavi and Lev, 2017). Both exogenously derived siRNAs (exo-siRNAs) and endogenous small RNAs, such as endo-siRNAs and PIWI-interacting small RNAs (piRNAs), can trigger heritable RNAi. Transgenerational inheritance of RNAi allows organisms to remember their exposure to genome parasites, such as viruses and transposons, transmit the experience to descendants, and promote evolutional advantages to enable selection for physiologically important traits. Although many lines of evidence have demonstrated that the RNAi machinery and small RNAs are involved in the initial establishment and ultimate maintenance of silencing, the precise nature of the trait-specific information that is transmitted from the parents to their progeny remains largely unclear (Ashe et al., 2012; Buckley et al., 2012; Burton et al., 2017; Kalinava et al., 2017; Klosin et al., 2017; Lev et al., 2017; Luteijn et al., 2012; Marre et al., 2016; Shirayama et al., 2012; Spracklin et al., 2017; Weiser et al., 2017).

The mechanisms of transgenerational inheritance of RNAi are widely being investigated in *C. elegans*. The RNAi-mediated silencing effect can be transmitted *via* parental gametes (Alcazar et al., 2008; Grishok et al., 2000) and its maintenance depends on the expression of the targeted genes in germline and post-transcriptional mechanisms (Minkina and Hunter, 2017). Chromatin-modifying enzymes and their associated factors, including HPL-2, SET-25, SET-32, MES-2, and HDA-4 are engaged in the re-establishment and maintenance of transgenerational gene silencing (Ashe et al., 2012; Luteijn et al., 2012; Mao et al., 2015; Shirayama et al., 2012; Vastenhouw et al., 2006). The RNAi spreading defective factors RSD-2 and RSD-6 also promote genome silencing by maintaining siRNA populations (Sakaguchi et al., 2014). Especially, the nuclear RNAi defective (Nrde) pathway plays essential roles in the inheritance of RNAi silencing (Ashe et al., 2012; Buckley et al., 2012; Burton et al., 2011; Gu et al., 2012; Mao et al., 2015; Marre et al., 2016; Shirayama et al., 2012; Weiser et al., 2017). The germline-expressed nuclear Argonaute protein HRDE-1 may carry heritable small RNAs and engage in RNAi inheritance (Buckley et al., 2012). It was shown that the nuclear RNAi pathway maintains siRNA expression in the progeny of dsRNA-treated animals (Burton et al., 2011).

Although the precise nature of the trait-specific information that is directly inherited remains unknown, both small RNAs and dsRNAs have been reported to transmit from the parents to progeny in *C. elegans* (Rechavi and Lev, 2017). In addition, it is unclear whether and to what extent the cytoplasmic RNAi machinery contributes to the inheritance of RNAi (Spracklin et al., 2017). The cytoplasmic Argonaute protein WAGO-1 has been implicated in the maintenance of silencing (Shirayama et al., 2012). Yet another study showed that WAGO-1 is not required for RNAi inheritance of a *pie-1_p_::gfp::h2b* transgene (Buckley et al., 2012).

To further understand the mechanisms of RNAi inheritance, we searched for factors required for silencing a germline-expressed *h2b::gfp* transgene in the progenies of animals exposed to exogenous dsRNA. We identified the cytoplasmic Argonaute protein WAGO-4. WAGO-4 is specifically required for the exogenous dsRNA-induced silencing of germline-expressed genes. WAGO-4 binds to germline-enriched 22G-RNAs containing untemplated addition of uracil at 3’ ends, which depends on the poly(U) polymerase CDE-1. After fertilization, in the zygotes and early embryos, WAGO-4 exhibits asymmetrical translocation to the germline. Therefore, we conclude that cytoplasmic RNAi machineries also contribute to the inheritance of RNAi, likely through the Argonaute protein WAGO-4.

## Results

### WAGO-4 is required for inheritance of RNAi

DsRNA targeting *lin-15b* elicits a multivulva (Muv) phenotype in enhanced RNAi (*eri*) animals through the nuclear RNAi defective (Nrde) pathway (Guang et al., 2008). However, the animals exhibit the Muv phenotype only in the F1 progeny but not in the parent animals, suggesting that the progenies have inherited a silencing signal. We have previously tested a number of chromatin modification factors and Argonaute proteins by examining *lin-15*-RNAi induced Muv phenotype and *lir-1*-RNAi-induced larval arrest. However, we failed to identify any except for *nrde* genes and *mes-2* that were required for both RNAi-induced phenotypes (Mao et al., 2015). Interestingly, we found that an Argonaute gene, *wago-4*, was required for *lin-15*-RNAi-induced Muv phenotype in the F1 progeny but not for *lir-1*-RNAi-induced larval arrest in the parental generation, suggesting that WAGO-4 may act through the germline to mediate RNAi inheritance.

WAGO-4 belongs to the worm-specific clade Argonaute proteins (Yigit et al., 2006). We acquired two deletion alleles *tm1019* and *tm2401*, from the National BioResource Project (NBRP) (Figure S1A). In addition, we used dual sgRNA-mediated CRISPR/Cas9 technology to generate an allele, *ust42* (Figures S1A) (Chen et al., 2014). *tm1019* lacks the MID domain of Argonaute proteins; however, the remainder of the protein is translated in frame. The alleles, *tm2401* and *ust42*, result in stop codons and are likely to be null alleles. *wago-4(ust42)* has been outcrossed twice and is used as the reference allele.

We confirmed that WAGO-4 is involved in dsRNA-induced gene silencing by feeding RNAi targeting a number of soma- and germline-expressed genes. RDE-4 is a dsRNA-binding protein which is indispensable for feeding RNAi. *rde-4* mutants were resistant to the RNAi targeting *pos-1, lir-1*, and *lin-15b* (Figure S1B). The nuclear Argonaute protein NRDE-3 was required for dsRNA targeting both *lir-1* and *lin-15b*, the two examples used in nuclear RNAi analysis (Guang et al., 2008). *wago-4* was required for exogenous RNAi targeting *pos-1* or *lin-15b*, but not for RNAi targeting *lir-1* gene. These data suggest that WAGO-4 is involved in dsRNA-induced gene silencing. Since *pos-1* is mainly silenced by RNAi in cytoplasm (Guang et al., 2008), these data also suggest that WAGO-4 could conduct gene silencing in the cytoplasm.

We further examined whether *wago-4* is required for the inheritance of RNAi (Figure 1A). We first used a germline-expressed *pie-1p::gfp::h2b* (abbreviated as *h2b::gfp*) transgene as a reporter, which can inherit RNAi-induced gene silencing for multiple generations (Buckley et al., 2012). The nuclear Argonaute *hrde-1* was not required for exogenous *gfp* dsRNA to silence the *h2b::gfp* transgene in the parental generation but essential for the silencing in F1 generation (Figure 1B). Similarly, *wago-4* was not required for exogenous *gfp* dsRNA to silence the *h2b::gfp* transgene in the P0 generation, but was necessary for silencing in F1 progeny. Next we used a soma-expressed *sur-5::gfp* transgene as another reporter of RNAi inheritance. *sur-5::gfp* was silenced by exogenous *gfp* dsRNA in both P0 and F1 generations in wild-type animals (Figure 1C). In *wago-4* mutant animals, *sur-5::gfp* was silenced in P0 generation but desilenced in F1 generation, suggesting that the progeny failed to inherit the silencing effect. Interestingly, the fluorescence intensity of both *h2b::gfp* and *sur-5::gfp* transgenes did not reach the levels of untreated animals in *wago-4* mutants, suggesting the presence of a weak RNAi inheritance even in the absence of *wago-4* (Figures 1B and 1C). The *C. elegans′* genome encodes 27 Argonaute proteins, twelve of which belong to the WAGO clade (Yigit et al., 2006). We speculate that other WAGO proteins may function redundantly with WAGO-4 in mediating the inheritance of RNAi.

**Figure 1:**
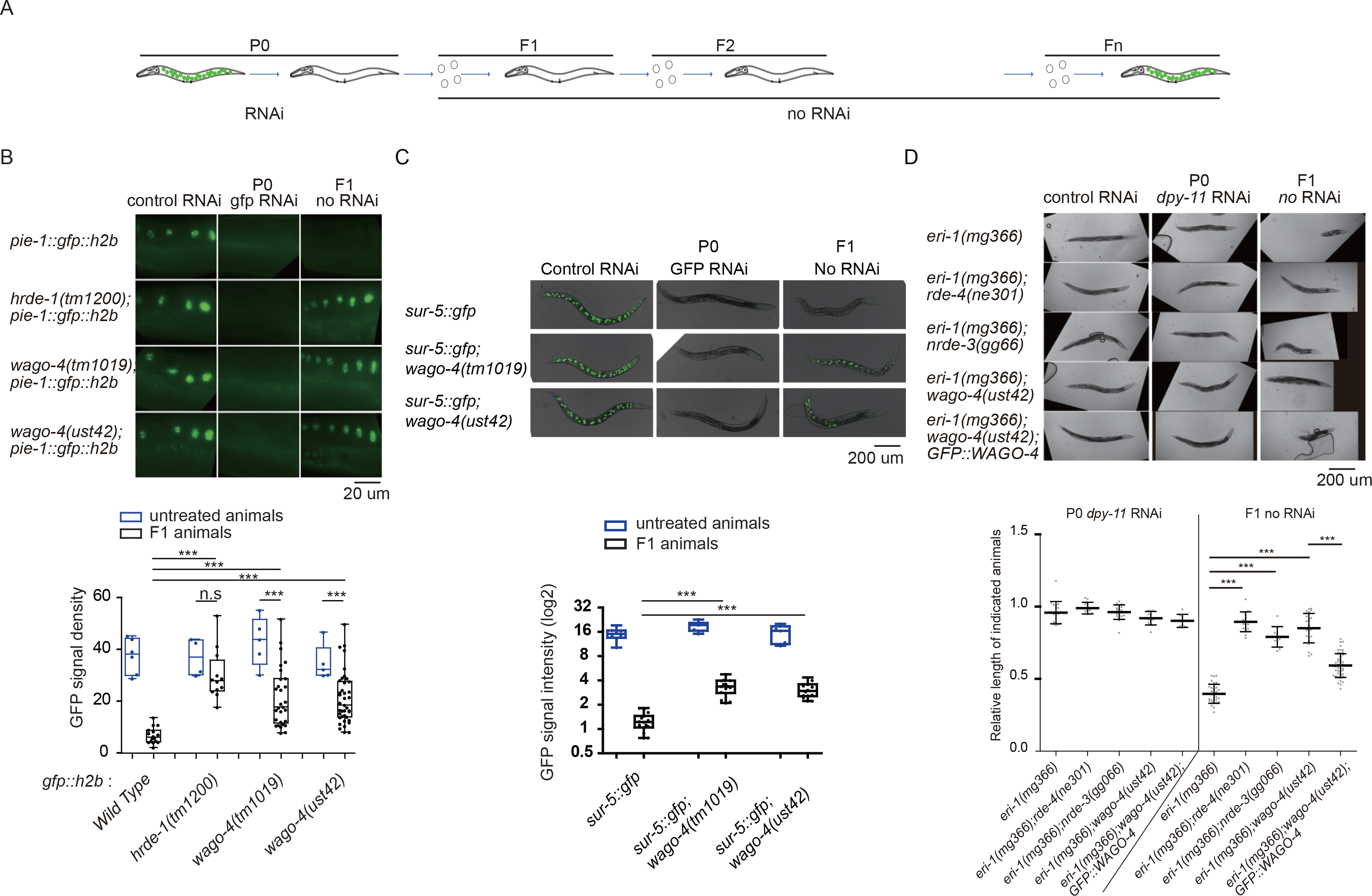
WAGO-4 is required for the inheritance of RNAi. (a) A scheme of RNAi procedures. (b) *pie-1_p_::gfp::h2b* and (c) *sur-5::gfp* transgenic animals were exposed to bacteria expressing *gfp* dsRNA, respectively. F1 embryos were isolated and grown on control bacteria in the absence of further *gfp* dsRNA treatment. GFP expression was imaged (upper panel) and the levels were scored (lower panel). (d) Rescue assay with the GFP::WAGO-4 transgene. The indicated animals were treated with *dpy-11* dsRNA and F1 animals were transferred to control bacteria (upper panel). The levels of dumpyness were scored (lower panel). ***p<0.001.

To further demonstrate that WAGO-4 promotes RNAi inheritance, we generated a single-copy WAGO-4 transgene *wago-4p::3xFLAG::GFP::WAGO-4* (abbreviated as *GFP::WAGO-4*) using MosSCI technology (Frokjaer-Jensen et al., 2014). This transgene rescued *pos-1* and *mex-5* dsRNA-elicited embryonic lethality and partially rescued *lin-15* dsRNA-induced Muv phenotype in *wago-4(ust42)* animals (Figures S1C-E), suggesting that the GFP::WAGO-4 represents the function and activity of endogenous WAGO-4 protein. *dpy-11* is an ortholog of human TMX1 and TMX4 genes and is required for body morphogenesis (Minkina and Hunter, 2017). dsRNA targeting *dpy-11* in *eri-1* parental animals induced a dumpy phenotype in the F1 progeny in the absence of further RNAi, suggesting that the RNAi signal has been inherited (Figure 1D). While *rde-4* animals did not respond to feeding RNAi at all, the depletion of NRDE-3 or WAGO-4 both caused the loss of dumpy phenotype in F1 animals, suggesting that NRDE-3 and WAGO-4 were required for the inheritance of RNAi-targeting somatic genes. The introduction of GFP::WAGO-4 transgene rescued the inheritance defects in *wago-4* mutants (Figure 1D), which confirmed the role of WAGO-4 in RNAi inheritance.

### WAGO-4 is a germline-expressed Argonaute protein and required for the silencing of germline-expressed genes

We downloaded the expression data of *wago-4* from Wormbase (version WS260). *wago-4* was exclusively detected in the hermaphrodite germline but not in the soma (Figure 2A). Consistently, GFP::WAGO-4 was exclusively expressed in the germline and all oocyte cells in gravid adults in hermaphrodites but not significantly expressed in the male germline (Figures 2B and S2A). In early embryos, WAGO-4 is expressed in the P1 and EMS cells (Figure S2B). In late embryos, WAGO-4 was exclusively expressed in Z2/Z3 cells. Interestingly, we observed that WAGO-4 accumulated at some distinct perinuclear foci (Figure 2B). Many RNA processing and regulatory factors, including the RNAi machinery, are enriched in the perinuclear region and exhibit distinct foci, which are termed P-granules (Chen et al., 2016; Claycomb et al., 2009; Gu et al., 2009; Tu et al., 2015; Wedeles et al., 2013). We crossed *GFP::WAGO-4* with a chromatin marker strain *mCherry::H2B* and the P-granule marker strain *mRuby::PGL-1* respectively and found that WAGO-4 partially co-localized with the P-granule marker PGL-1 but not with the chromatin marker (Figure 2C).

**Figure 2:**
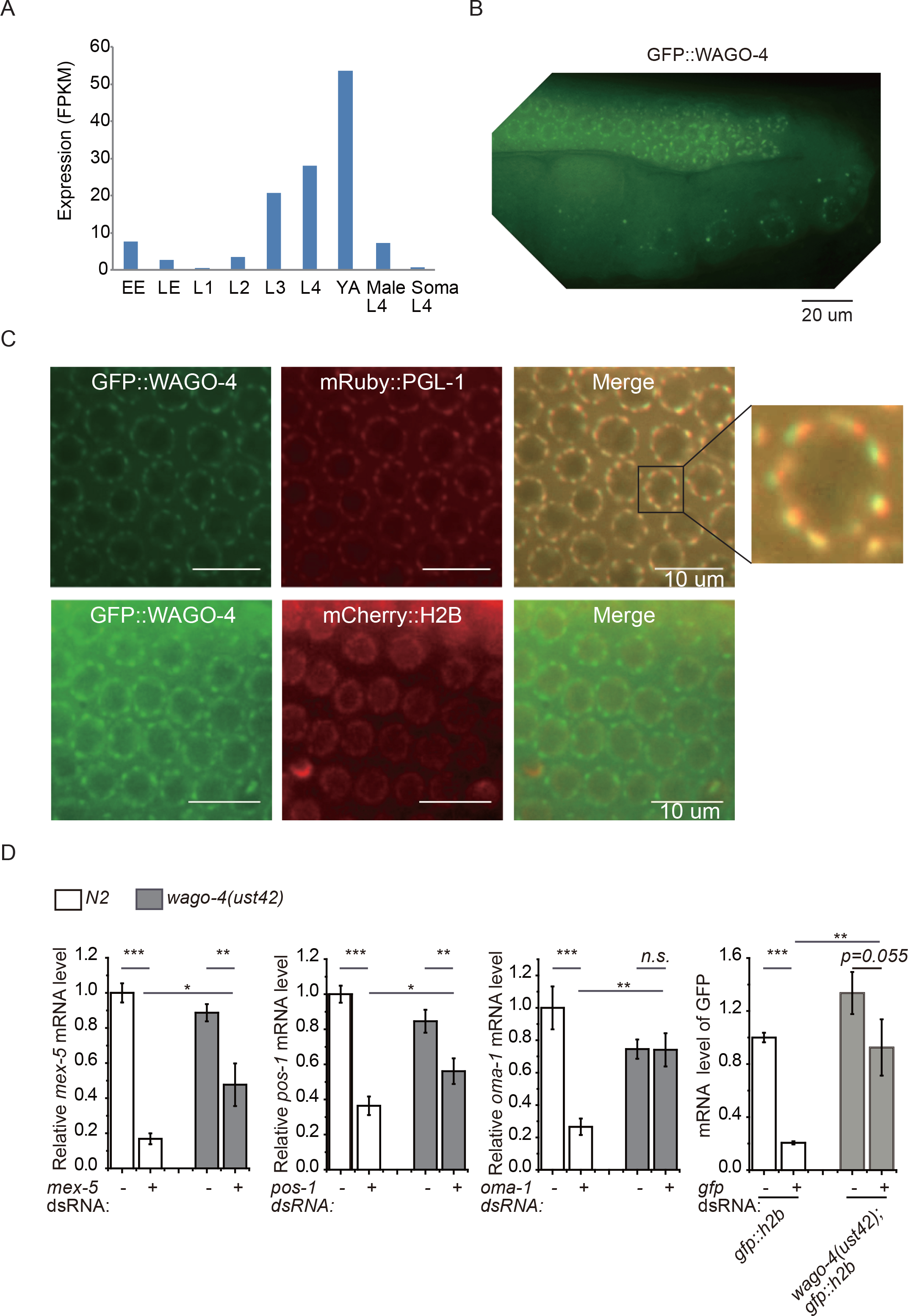
WAGO-4 is a germline-expressed Argonaute protein. (a) The expression levels of *wago-4* mRNA at different developmental stages. Data were downloaded from Wormbase (version WS260). EE, early embryos; LE, late embryos; YA, young adults. (b) Images of the germline of the GFP::WAGO-4 strain. (c) Pachytene germ cells of GFP::WAGO-4 and the chromatin marker H2B::mCherry and P-granule marker mRuby::PGL-1 were imaged. (d) Wild-type and *wago-4(-)* animals were treated with indicated dsRNAs. Total RNAs were isolated at L4 stage of P0 generation and the indicated mRNAs were quantified by real-time PCR. mean ± s.d. n=3. *p<0.05, **p<0.01, ***p<0.001, ns, not significant.

WAGO-4 lacks the DDH catalytic triad of amino acids considered necessary for Argonaute-based slicer activity (Yigit et al., 2006). We tested whether WAGO-4 was required for RNAi targeting germline-expressed mRNAs. We fed L1 animals on dsRNA, harvested L4 animals of the same generation 48 hours later, and quantified mRNA levels by real-time PCR. We found that WAGO-4 was required for the decrease of the targeted mRNAs (Figure 2D), suggesting that either WAGO-4 contains some slicer activity, or siRNA/WAGO-4 complex silences targeted mRNAs through other mechanisms. Interestingly, *mex-5* and *pos-1* mRNAs were not fully desilenced in *wago-4* mutants, suggesting that other germline Argonaute proteins may function redundantly with WAGO-4 to silence germline genes.

### WAGO-4 and HRDE-1 act differently in promoting RNAi inheritance

Both HRDE-1 and WAGO-4 are required for the inheritance of RNAi, yet they exhibit distinct subcellular locations. We hypothesized that they function via different mechanisms or at distinct steps in the process of transmitting siRNAs and/or re-establishing the silencing effects in the progeny.

To test this hypothesis, we fed *sur-5::gfp* parental animals with *gfp* dsRNA and examined *gfp* expression in F1 embryos and L4 larva without further feeding RNAi. *sur-5::gfp* was silenced in wild-type animals but not in *rde-1* mutants in the parental and F1 animals (Figures 3A and S3). In *hrde-1* and *nrde-1/2/4* mutants, *sur-5::gfp* was silenced in both parent animals and F1 embryos, indicating that the animals defective for nuclear RNAi are able to inherit *gfp* silencing signals. During larval development, the heritable *gfp* silencing was relieved in *nrde-1/2/4* mutants (Figure S3), which was also shown in *nrde-3* mutants previously (Burton et al., 2011). However, *hrde-1(-)* mutants still exhibited gene silencing in L4 larva in the soma, further supporting that *hrde-1(-)* animals inherited silencing signals (Figures 3A and S3). Strikingly, when *wago-4(-);sur-5::gfp* animals were fed with *gfp* dsRNA, the *sur-5::gfp* was expressed in both F1 embryos and L4 larva (Figure 3A), although not to the levels of the *rde-1* mutant. These results suggest that WAGO-4 is likely engaged in transmitting small RNAs from the parents to zygotes or at least at an earlier step than HRDE-1 to re-establish the silencing state in the progeny.

**Figure 3:**
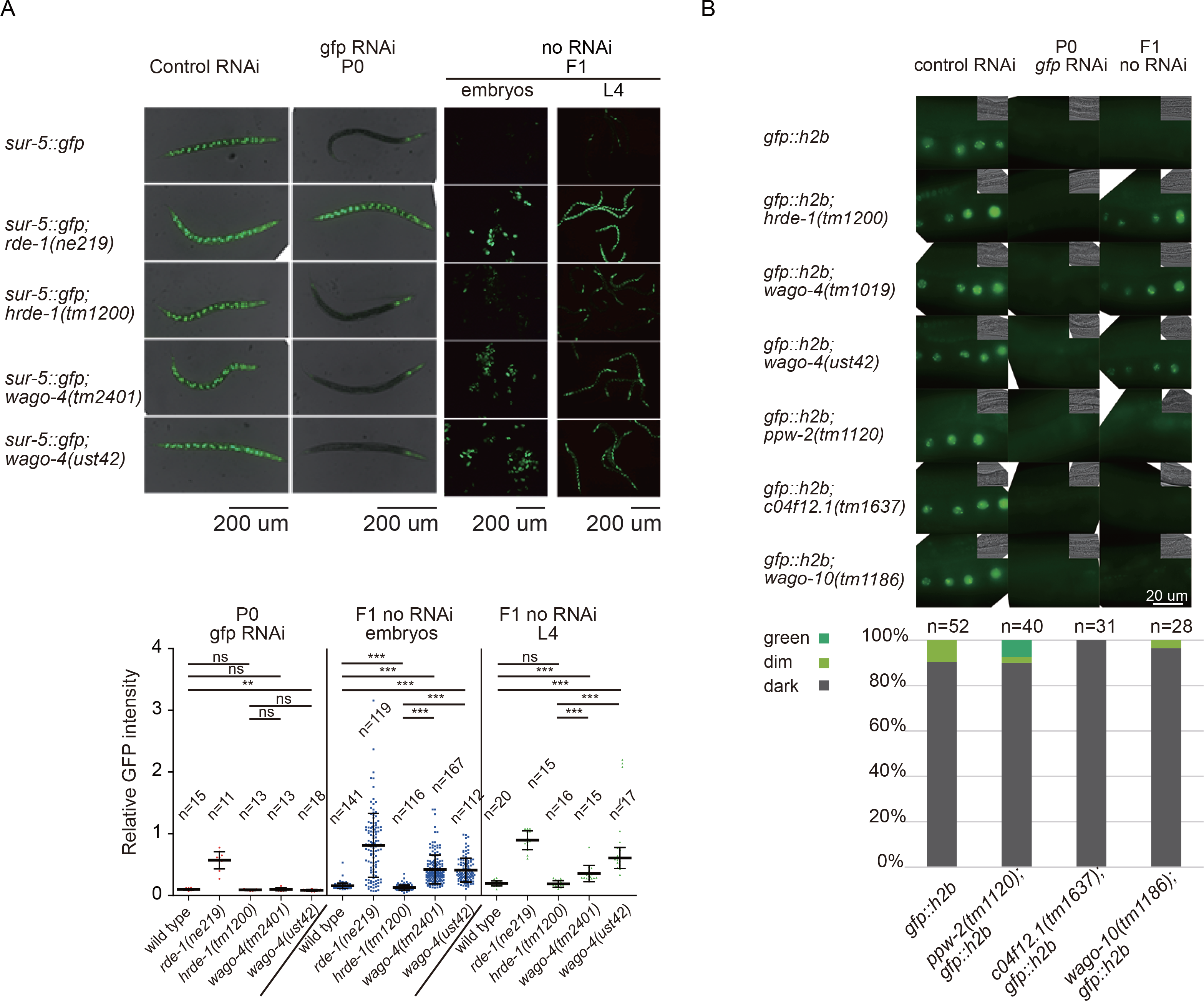
WAGO-4 and HRDE-1 act differently to promote RNAi inheritance. (a) Images of indicated animals after *gfp* RNAi. *sur-5::gfp* animals were fed on *gfp* dsRNA and the F1 embryos were incubated on control bacteria. Upper panel: fluorescent images of indicated animals after *gfp* RNAi. Bottom panel: GFP intensity levels of the indicated animals were measured by ImageJ. The number of animals assayed is indicated. *p<0.05, **p<0.01, ***p<0.001, ns, not significant. (b) Images of germline cells of indicated animals. *h2b::gfp* animals were fed on *gfp* dsRNA and the F1 embryos were incubated on HT115 control bacteria. The *gfp* levels were scored and shown in the lower panel.

Previous genetic screens failed to identify WAGO proteins other than HRDE-1 that was required to mediate the inheritance of RNAi targeting *h2b::gfp* transgene (Buckley et al., 2012). The identification of WAGO-4 in RNAi inheritance suggested that these WAGO proteins should be re-investigated individually. We selected three other Argonaute proteins, in which C04F12.1 is closest to CSR-1, WAGO-10 is closest to HRDE-1, and PPW-2 is closest to WAGO-4 based on sequence comparison (Gu et al., 2009). However, none of these three proteins was required for the inheritance of RNAi targeting the *h2b::gfp* transgene (Figure 3B). This data suggests that either WAGO-4 and the nuclear Argonaute HRDE-1 and NRDE-3 played special roles in RNAi inheritance, or C04F12.1, WAGO-10, and PPW-2 exhibit functional redundancy (Shirayama et al., 2012).

### WAGO-4 acts in multigenerational inheritance of RNAi

To test whether *wago-4* is required for multigenerational inheritance of RNAi, we fed germline-expressed *h2b::gfp* animals with *gfp* dsRNA and scored *gfp* levels in later generations. The *h2b::gfp* transgene was silenced in the parental generation and the silencing was maintained to F1 and F2 generations, although at reduced levels (Figure 4A). Both *hrde-1* and *wago-4* were required for the inheritance of *gfp* silencing to F1 and F2 generations.

**Figure 4.**
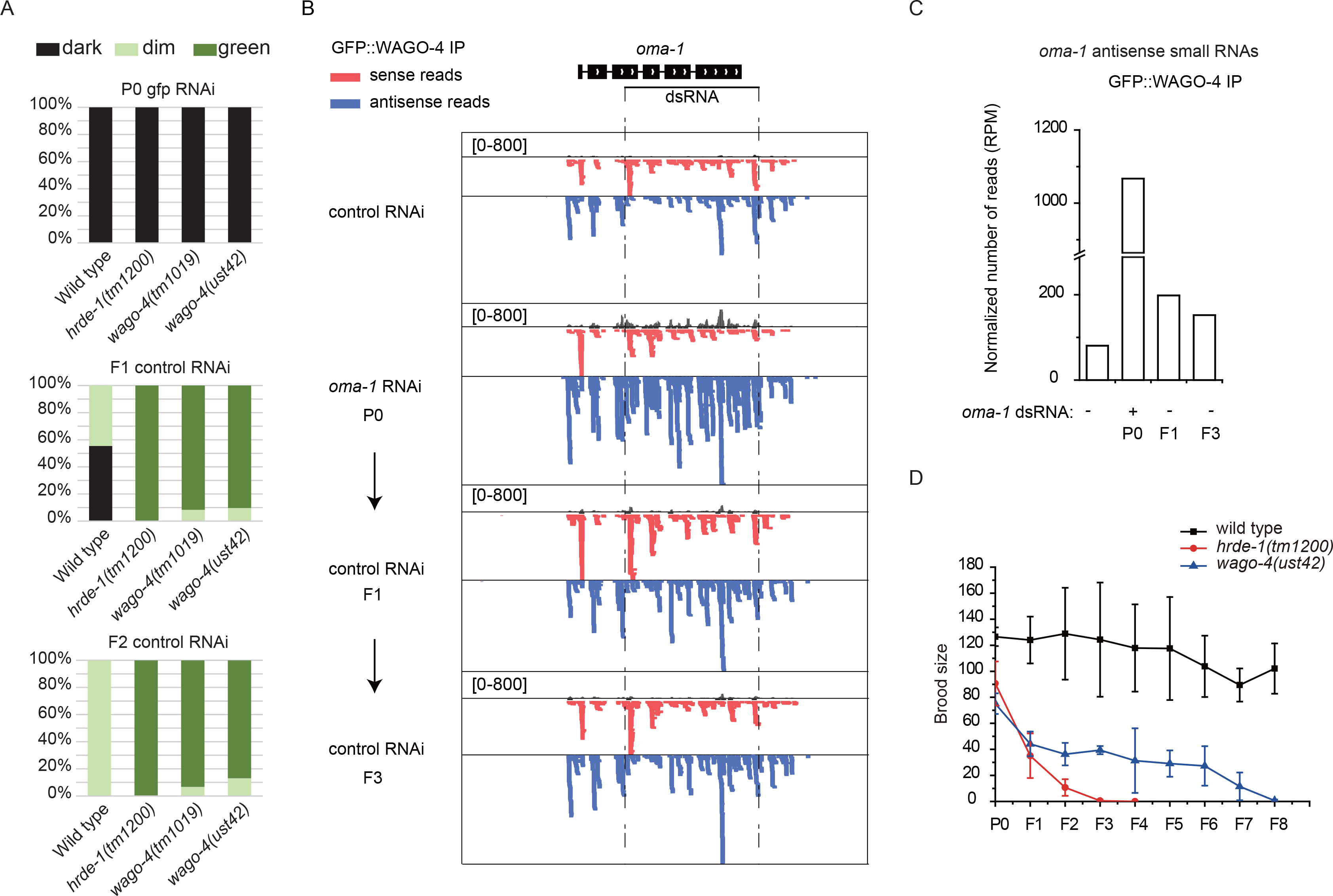
WAGO-4 acts in multigenerational inheritance of RNAi. (a) *pie-1_p_::gfp::h2b* transgenic animals were exposed to bacteria expressing *gfp* dsRNA. F1 and F2 embryos were isolated and grown on control bacteria in the absence of further *gfp* dsRNA treatment. Percentage of F1 animals expressing GFP was counted (N>50). (b) WAGO-4-associated small RNAs targeting *oma-1* locus is shown. Red ticks are sense reads to *oma-1* sequence, blue ticks are antisense reads. GFP::WAGO-4 animals were treated with *oma-1* dsRNA at P0 generation and then fed on control bacteria in subsequent generations. GFP::WAGO-4 was immunoprecipitated from young adult animals and the associated small RNAs were deep sequenced. Animals fed on control RNAi bacteria were set as control. (c) Normalized number of antisense reads of indicated animals. (d) Brood size of indicated animals by Mrt assay at 25℃. N=20, mean±s.d.

Then we fed GFP::WAGO-4 animals with dsRNA targeting *oma-1* only in P0 generation and isolated WAGO-4-bound siRNAs in later generations, followed by small RNA deep sequencing. WAGO-4 bound to *oma-1* siRNAs not only in the P0 and F1 generations, but also in the F3 generation (Figures 4B and 4C), suggesting that WAGO-4 act in multigenerational inheritance of RNAi.

Many RNAi inheritance mutants, including *hrde-1*, exhibit a Mrt phenotype that gradually loses the fertility along generations (Buckley et al., 2012; Spracklin et al., 2017). We compared the Mrt phenotype of *hrde-1* and *wago-4* animals by examining their brood size at 25℃ (Figure 4D). While *hrde-1* animals gradually lost the fecundity and achieved sterility at F3/F4 generations, *wago-4* animals exhibited a weaker Mrt phenotype and got sterile at the F8/F9 generations.

### WAGO-4 binds to 22G-RNAs that target germline-expressed protein coding genes

To better understand the functions of WAGO-4, we immunoprecipitated WAGO-4 from gravid P0 animals before and after exogenous *lin-15b* RNAi, purified WAGO-4-associated mRNAs, and quantified mRNAs by real-time PCR. We found that WAGO-4 bound to *lin-15b* mRNA after *lin-15b* RNAi (Figures S4A and 5A), suggesting that WAGO-4 can be directed to targeted mRNAs by exogenous dsRNA. We then deep-sequenced WAGO-4-associated small RNAs in a 5’-phosphate-independent manner (Zhou et al., 2014). Small RNA reads were aligned to the *C. elegans* transcriptome (WS243 assembly), and the number of reads targeting each gene was counted. WAGO-4 preferentially binds to small RNAs antisense to *lin-15b* mRNA in the presence of exogenous *lin-15b* dsRNA (Figure 5B). Although most of the reads locate inside the dsRNA region, a portion of siRNAs are derived from outside of the dsRNA region (Figure 5C). *lin-15b* and *lin-15a* localize in an operon and are transcribed as a single pre-mRNA, spliced in the nucleus, and exported individually into the cytoplasm. The siRNAs targeting *lin-15a* were not increased, which is consistent with the idea that RNA dependent RNA polymerases (RdRPs) use mRNAs in the cytoplasm, but not pre-mRNAs in the nucleus, as templates to amplify secondary siRNAs, and suggests that WAGO-4 bound to secondary siRNAs (Pak and Fire, 2007).

**Figure 5:**
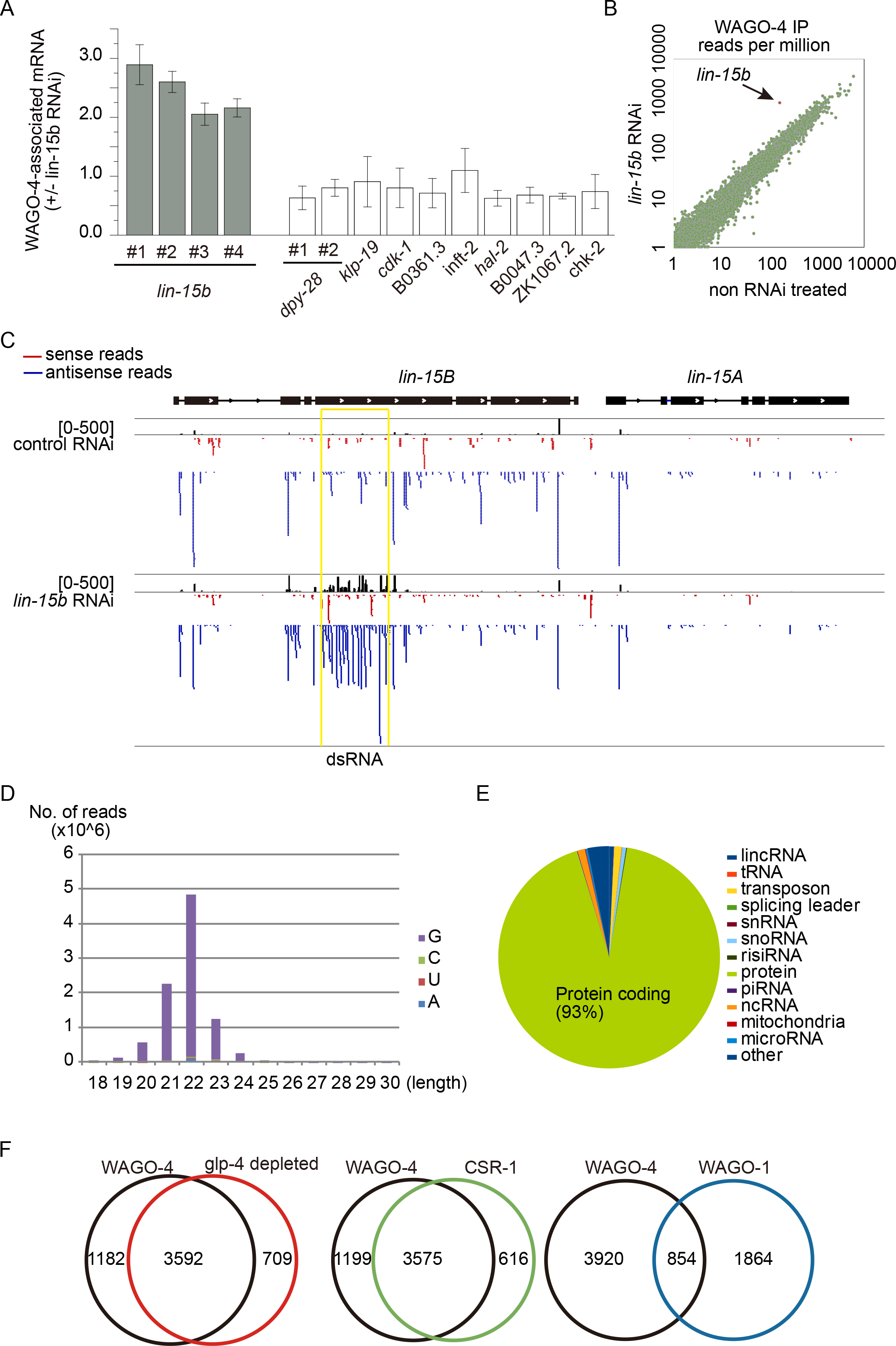
WAGO-4 binds to 22G-RNAs that target germline-expressed protein coding genes. (a) WAGO-4 was immunoprecipitated at young gravid stage after *lin-15b* RNAi, and the associated RNAs were quantified by real-time PCR. mean ± s.d. n=3. (b) WAGO-4 was immunoprecipitated at young gravid stage, and the associated siRNAs were deep sequenced. The number of reads were normalized and then compared. (c) Reads targeting *lin-15b/a* genomic loci were plotted. Yellow box indicates the dsRNA-targeted region. (d) The length distribution and the first nucleotides of WAGO-4-associated siRNAs were analyzed. (e) WAGO-4-associated 22G-RNAs were grouped into different categories. (f) WAGO-4-associated 22G-RNAs were compared with other known siRNA categories.

We further characterized WAGO-4-associated endogenous siRNAs. WAGO-4-associated endogenous siRNAs were 22 nt in length (Figure 5D), and the vast majority of the siRNAs started with G or GA at their 5’ ends (Figures 5D and S4B), which are consistent with the properties of 22G-RNA. Approximately 93% of WAGO-4-bound 22G-RNAs target protein-coding genes (Figure 5E). We selected potential WAGO-4 targets that had greater than 25 reads per million and identified 4,774 genes (Table S1).

We then compared WAGO-4-bound 22G-RNAs to other siRNA categories (Gu et al., 2009; Tu et al., 2015; van Wolfswinkel et al., 2009). Consistent with its germline expression, WAGO-4 targets exhibited a pronounced overlap with *glp-4*-dependent genes (Figure 5F). Although WAGO-1, WAGO-4, and CSR-1 are all expressed in germline, WAGO-4 targets dramatically overlapped with those of CSR-1 but not WAGO-1, suggesting that WAGO-4 and CSR-1 may regulate the same cohorts of protein-coding genes in the germline. However, although *csr-1* mutant is homozygous lethal and exhibits chromosomal segregation defects (Claycomb et al., 2009; Gerson-Gurwitz et al., 2016), *wago-4* animals are largely normal and have similar brood size to that of wild-type animals at 20℃ (Figure S4C). We quantified the mRNA levels of WAGO-4 targets, but failed to detect pronounced desilencing of the mRNA levels in *wago-4* mutants compared to wild-type animals (Figure S4D). These data suggest that CSR-1 and WAGO-4 may function differently in gene regulation and animal development, although they bind to the same cohorts of germline-expressed 22G-RNAs.

### WAGO-4-associated 22G-RNAs contain untemplated 3’-end uridylation

We analyzed the 3’-ends of WAGO-4-associated 22G-RNAs and found that approximately 7.6% of them contained untemplated addition of a uracil (Figure 6A). For reads longer than 23 nt, approximately 31% of them contained an extra U and 24.5% contained extra UU dinucleotides. We then examined the nucleotide distribution at each position (Figure S5A). Although the first two nucleotides exhibit a strong propensity towards the GA dinucleotide sequence and the nucleotides at position 3 to 5 have a modest enrichment of A, the four nucleotides (A, U, C, and G) are approximately equally distributed at each position in the middle of 22G-RNAs in wild type animals. At the 3’-end, WAGO-4-associated 22G-RNAs exhibited a strong representation towards U.

**Figure 6:**
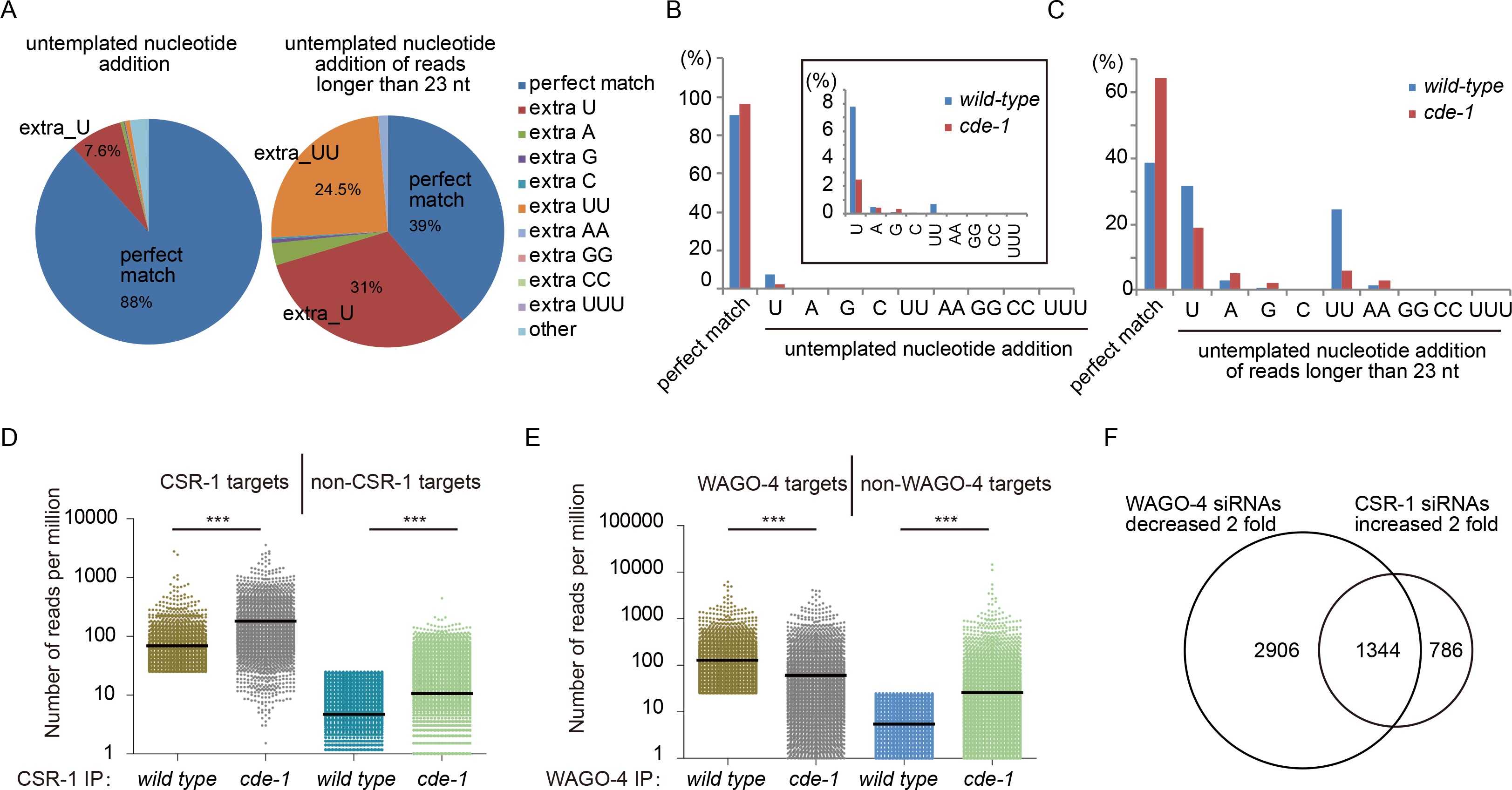
CDE-1 uridylates WAGO-4-associated 22G-RNAs. (a) The untemplated 3’-end addition of nucleotides in WAGO-4-bound 22G-RNAs was analyzed. Left, total reads; right, reads longer than 23 nt. (b) Comparison of the untemplated addition of nucleotides in WAGO-4-associated 22G-RNAs in wild-type animals and *cde-1* mutants. The insert plot is a zoom-in of the main figure. (c) Comparison of the untemplated addition of nucleotides in WAGO-4-bound 22G-RNAs for reads longer than 23 nt. (d) Analysis of CSR-1 22G-RNAs in *cde-1* mutants. (e) Analysis of WAGO-4 22G-RNAs in *cde-1* mutants. (f) Comparison of WAGO-4 and CSR-1 22G-RNAs in *cde-1* mutants. ***p<0.001.

CDE-1 is a polyuracil polymerase that adds untemplated uracils to 22G-RNAs (van Wolfswinkel et al., 2009). We examined whether CDE-1 is involved in uridylating WAGO-4-associated 22G-RNAs by deep sequencing WAGO-4-associated small RNAs from *cde-1(tm1021);3xFLAG::GFP::WAGO-4* animals. The vast majority of WAGO-4-associated 22G-RNAs still targeted protein-coding genes (Figure S5B) and exhibited a tendency towards GA at 5’-end (Figure S5A), suggesting that CDE-1 is not essential for the biogenesis of WAGO-4-associated siRNAs. Yet the untemplated addition of uracil was decreased in *cde-1* mutants. The percentage of an extra U was 7.6% in wild type animals and 2.5% in *cde-1* animals (Figure 6B). For reads longer than 23 nt, the percentage of an extra U was 31% in wild type animals and 19% in *cde-1* animals respectively (Figure 6C).

Noticeably, in *cde-1* mutants, WAGO-4 bound approximately 24-fold more antisense ribosomal siRNAs (risiRNA) (31,409 reads per million in *cde-1* vs. 1,322 reads per million in wild-type animals) and 22-fold more small RNAs related to splicing leaders (Figure S5B). We re-analyzed the published deep sequencing data sets and identified similar increase of risiRNAs in *cde-1* mutants (data not shown) (van Wolfswinkel et al., 2009). How CDE-1 is involved in risiRNA biogenesis remains unclear (Zhou et al., 2017).

CDE-1-mediated uridylation is thought to destabilize CSR-1-bound 22G-RNAs (van Wolfswinkel et al., 2009). We re-analyzed published data of small RNAs associated with CSR-1 in *cde-1* and wild-type animals (van Wolfswinkel et al., 2009). CSR-1 targets with greater than 25 reads per million were selected. As reported, CSR-1 bound more 22G-RNAs of CSR-1 targets in the *cde-1* mutant (Figure 6D). In contrast, WAGO-4 bound fewer 22G-RNAs of WAGO-4 targets in the *cde-1* mutant (Figure 6E). Previous studies suggested that there exists a loading balance between WAGO pathway and CSR-1 pathway, which can be regulated by CDE-1 mediated uridylation (de Albuquerque et al., 2015; Phillips et al., 2015; van Wolfswinkel et al., 2009). We selected WAGO-4-associated 22G-RNAs that were decreased 2-fold and CSR-1-associated 22G-RNAs that were increased 2-fold in the *cde-1* mutant, and found that 1,344 genes were decreased in WAGO-4 binding but simultaneously increased in CSR-1 binding (Figure 6F). These data suggest that CDE-1-engaged uridylation likely stabilizes the binding of 22G-RNAs to WAGO-4 but destabilizes the binding to CSR-1.

We then asked whether *cde-1* is required for the inheritance of RNAi. Consistent with a recent report, *cde-1* was not required for exogenous dsRNA to silence the germline-expressed *h2b::gfp* transgene in the parental generation but was imperative for silencing in F1 progeny (Figure S5C) (Spracklin et al., 2017). Taken together, we hypothesize that CDE-1 mediates RNAi inheritance by uridylating WAGO-4-bound 22G-RNAs.

### WAGO-4 promotes the transmission of siRNAs for RNAi inheritance

We hypothesized that WAGO-4 is able to directly transmit siRNAs from parents to progeny. Three lines of evidence cumulatively supported this idea. First, WAGO-4 was expressed in both germline cells and oocytes (Figures 2B and 7A). In −3 to −1 oocytes, WAGO-4 still exhibited distinct foci, but changed localization from peri-nuclear region to cytoplasm. Second, WAGO-4 is expressed in one-cell embryos (Figure 7B). Early in the one-cell stage before chromosomal alignment, WAGO-4 is evenly distributed in the zygote without pronounced aggregation. After chromosomal alignment, WAGO-4 began to accumulate in P-granule foci at the posterior ends. The expression of WAGO-4 in −1 oocyte and the one-cell embryos suggest that WAGO-4 is able to carry siRNAs from parents to progeny.

**Figure 7:**
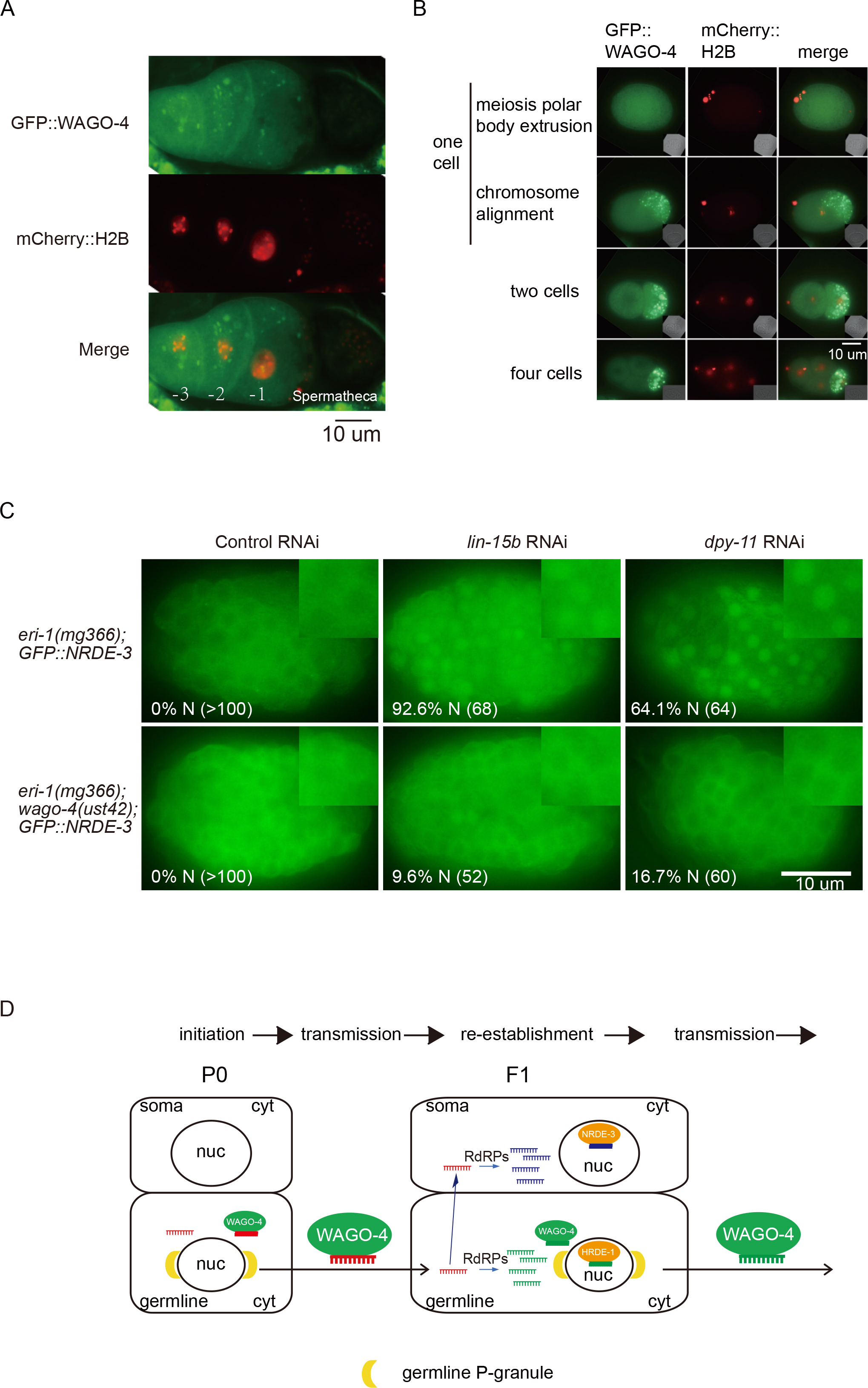
WAGO-4 promotes transmission of siRNAs from parents to progeny. (a, b) Images of oocytes and early embryos of the *GFP::WAGO-4;mCherry::H2B* strain. (c) Images of F1 embryos of the indicated animals after feeding RNAi targeting L4 animals for 24 hours. The percentage of nuclear localized GFP::NRDE-3 in F1 embryos was indicated. The number of scored animals were indicated in the parentheses. N, nucleus. (d) A working model of WAGO-4-mediated inheritance of RNAi. WAGO-4 is involved in transmitting siRNAs along generations as well as conducting gene silencing in the germline. Inherited small RNAs targeting soma-expressed genes are translocated to the soma, amplified, and subsequently silence somatic gene expression. Inherited siRNAs targeting germline-expressed genes are amplified and silence gene expression in the germline of the progeny. WAGO-4 transmits germline siRNAs to the next generation and maintains the silencing status for multiple generations.

Third, we used the subcellular localization of GFP::NRDE-3 as a reporter of siRNA abundance to test whether the germline-expressed WAGO-4 is required for the siRNA accumulation in the soma in the progeny. NRDE-3 is a somatic Argonaute protein that localizes to the nucleus when it binds to siRNAs. In *eri-1* animals, NRDE-3 accumulates in the cytoplasm in the absence of siRNA binding (Guang et al., 2008). In *eri-1;GFP::NRDE-3*, but not *eri-1;wago-4;GFP::NRDE-3*, animals, feeding RNAi targeting soma-expressed *dpy-11* and *lin-15b* genes elicited a cytoplasm-to-nucleus translocation of GFP::NRDE-3 in the F1 embryos (Figure 7C). Since the presence of mRNA templates were required for the amplification of secondary siRNAs, the siRNAs targeting soma-expressed genes (*dpy-11* and *lin-15b*) were unable to be amplified in the germline. The fact that the inheritance of RNAi targeting somatic genes depended on the germline-expressed WAGO-4 suggested that siRNAs were likely transmitted from the parental germline to progeny embryos by WAGO-4. These siRNAs were then transported from the germline to soma to be amplified and induced cytoplasm-to-nucleus translocation of NRDE-3.

Taken together, these data suggest that WAGO-4 may directly promote the transmission of siRNAs from the parents to the progeny.

## Discussion

The inheritance of RNAi in *C. elegans* results from multiple steps, including initiation, transmission and re-establishment of silencing. Here, we demonstrated that the cytoplasmic Argonaute protein WAGO-4 and the poly(U) polymerase CDE-1 play important roles in promoting inheritance of RNAi.

### WAGO-4 promotes RNAi inheritance

The nuclear RNAi pathway has been shown to play essential roles in transgenerational inheritance of RNAi (reviewed by (Heard and Martienssen, 2014; Rechavi and Lev, 2017). Here, we demonstrated that WAGO-4 promotes the inheritance of RNAi. WAGO-4 partially localized to the perinuclear P-granule foci, suggesting that cytoplasmic RNAi machinery is also involved in the inheritance of RNAi. P-granules contain numerous RNA processing and regulatory factors, including a few WAGOs (Chen et al., 2016). In a recent genetic screen, Spracklin et al. identified two cytoplasmic proteins GLH-1/VASA and HRDE-2 in transgenerational inheritance of RNAi (Spracklin et al., 2017), further supporting that the cytoplasmic factors are indispensable for the inheritance of RNAi. Previously, the cytoplasmic Argonaute protein WAGO-1 has been implicated in the maintenance of silencing (Shirayama et al., 2012). Yet another study showed that WAGO-1 is not required for RNAi inheritance of a *pie-1_p_::gfp::h2b* transgene (Buckley et al., 2012). Therefore, a systematic investigation of the function of WAGO-1 in RNAi inheritance is needed.

Our data suggest that WAGO-4 is involved in the transmission of siRNAs from the parents to progeny. First, WAGO-4 is expressed in both −1 oocyte and the one-cell embryos, suggesting that WAGO-4 is able to carry 22G-RNAs from the parental oocyte to the progeny. Second, using the subcellular localization of GFP::NRDE-3 as a siRNA reporter, we showed that the germline expressed Argonaute protein WAGO-4 is required for the production of siRNAs in the soma. A working model of RNAi inheritance was shown in Figure 7D.

Epigenetic changes can arise in the F1 embryos that are present inside the parents when the parental animals are exposed to environmental stimuli. These phenomena can be called as parental effects or intergenerational epigenetic inheritance (Anava et al., 2014; Heard and Martienssen, 2014; Houri-Zeevi and Rechavi, 2017). The epigenetic changes that are inherited and maintained for more than two or three generations are then called transgenerational inheritance. Stable maintenance of silencing requires an additional class of siRNAs that must be amplified in each generation. These siRNAs can be classified as secondary or tertiary siRNAs (Sapetschnig et al., 2015). WAGO-4 binds to siRNAs in F3 generation after initial RNAi, suggesting that WAGO-4 is engaged in transgenerational inheritance of RNAi and binds to secondary and tertiary siRNAs.

WAGO-4- and CSR-1-associated 22G-RNAs target similar cohorts of germline-expressed protein-coding genes. However, *csr-1* mutant is homozygous lethal, while *wago-4* mutants are largely normal when grew at 20℃, suggesting that these two Argonaute proteins may use different gene regulatory mechanisms (Shirayama et al., 2012). Interestingly, although WAGO-4 lacks the DDH catalytic triad of amino acids considered necessary for Argonaute-based slicer activity (Yigit et al., 2006), it was required for the silencing of the targeted mRNAs.

### Untemplated uridylation by CDE-1 is required for RNAi inheritance

An important question is how Argonaute proteins selectively identify and bind to their siRNA ligands from a pool of short nuclear acids in the cell. In *Arabidopsis*, the 5’ terminal nucleotide directs the sorting of small RNAs into different Argonaute complexes (Mi et al., 2008; Montgomery et al., 2008; Takeda et al., 2008). In *C. elegans*, 22G-RNAs may be sorted to distinct Argonaute proteins upon target recognition. Here, our data suggested that untemplated uridylation at the 3’-ends may function as an additional sorting signal by modulating the binding affinity of 22G-RNAs to different Argonaute proteins (de Albuquerque et al., 2015; Phillips et al., 2015; van Wolfswinkel et al., 2009). Alternatively, uridylation of 22G-RNAs may change their binding affinity to only one Argonaute, while more or less of the rest 22G-RNAs bind to the other Argonaute proteins.

The poly(U) polymerase CDE-1 (also known as PUP-1 and CID-1) adds untemplated uracil to the 3’ termini of RNAs in *C. elegans* (Kwak and Wickens, 2007; van Wolfswinkel et al., 2009). Previous studies suggested that a small RNA loading balance between WAGO pathway and CSR-1 pathway may exist, which is regulated by CDE-1 mediated uridylation (de Albuquerque et al., 2015; Phillips et al., 2015; van Wolfswinkel et al., 2009). Because CDE-1 interacts with a RNA-dependent RNA polymerase EGO-1, but not with the Argonaute protein CSR-1 (van Wolfswinkel et al., 2009), and EGO-1 is required for the production of 22G-RNAs and acts upstream of the loading of 22G-RNAs to Argonaute proteins, we speculated that uridylation happens upstream of small RNA loading to Argonaute proteins in *C. elegans*. Interestingly, in fission yeast, the uridyl transferase Cid16 can add non-templated nucleotides to Argonaute-bound small RNAs *in vitro* (Pisacane and Halic, 2017). Thus we could not exclude the possibility that CDE-1 might be able to uridylate Argonaute-bound siRNAs after loading, since we were only measuring the steady state levels of Argonaute-bound siRNAs by immunoprecipitation from worm extracts. Further *in vitro* biochemical experiments are required to examine the binding affinities of uridylated 22G-RNAs to different Argonaute proteins and how 3’ uridylation affects the stability and target recognition of small RNAs.

Noticeably, Spracklin et al. identified that CDE-1 is required for the inheritance of RNAi (Spracklin et al., 2017), for which the mechanism is not fully understood. Our data suggest that CDE-1 may mediate RNAi inheritance by regulating the binding of 22G-RNAs to WAGO-4. Yet *cde-1* mutants exhibited a weaker inheritance defects than *wago-4* animals. Therefore, the function of CDE-1 in RNAi inheritance needs further investigation.

### Experimental Procedures

#### Strains

Bristol strain N2 was used as the standard wild-type strain. All strains were grown at 20°C. The strains used in this study are listed in Supplemental Experimental Procedures.

#### RNAi

RNAi experiments were conducted as described previously (Timmons et al., 2001). HT115 bacteria expressing the empty vector L4440 (a gift from A. Fire) were used as controls. Bacterial clones expressing dsRNA were obtained from the Ahringer RNAi library (Kamath et al., 2003) and were sequenced to verify their identities. *lin-15b* RNAi clones were described previously (Guang et al., 2008).

#### RNAi inheritance assay

Synchronized L1 animals of the indicated genotypes were exposed to bacteria expressing *gfp* dsRNA. F1 embryos were collected by hypochlorite/NaOH treatment and grown on HT115 control bacteria. The expression levels of GFP in both parental generation and progeny were visualized and scored.

Images were collected on Leica DM2500 microscope. For the quantitation of GFP intensity, the average fluorescence intensities of L4 worms (0.5-second exposure) and embryos (1-second exposure) were analyzed using ImageJ (Borges and Martienssen, 2015). The background of each individual animal was subtracted using "the rolling ball radius" model of 3x3 pixels with the smoothing feature disabled. For subcellular localization analysis, approximately ten germlines were imaged and analyzed by ImageJ.

Images of co-localization analysis were collected on Leica DM4 B microscope system.

### Mrt assay

*wago-4(ust42)(2x)* was successively outcrossed twice with N2 animals to get *wago-4(ust42)(4x)* before conducting Mrt experiments. Twenty larval stage L3 animals were singled onto OP50 plates at each generation and cultured at 25℃. The average brood size at each generation was calculated.

### Construction of plasmids and transgenic strains

For 3xFLAG::GFP::WAGO-4, the 3xFLAG::GFP coding region was PCR amplified from plasmid pSG085 with the primers 5′-AGCTCTTCCTATGGACTACAAAGACCATGAC-3’ and 5′-ATAGCTCCACCTCCACCTCCTTTGTATAGTTCATCCATGCC -3′. The predicted *wago-4* promoter was amplified from N2 genomic DNA with primers 5′-CGTGGATCCAGATATCCTGCAGGTTCGGTTACGCTCTCTCTCCAG -3′ and 5′-TGTAGTCCATAGGAAGAGCTGGCATCCTTCTC -3′. The *wago-4* coding region and predicted 3′ UTR were then amplified by PCR from N2 genomic DNA with primers 5′-GGAGGTGGAGGAGGTGGAGCTATGCCAGCTCTTCCTCCAGTC-3′ and 5′-GACTCACTAGTGGGCAGATCTCATCCCTGATGCCAAGCCGAC-3′. The plasmid pCFJ151 was digested with *Sbf*I and *Bgl*II. The three above PCR fragments were cloned into linearized pCFJ151 with the ClonExpress MultiS One Step Cloning kit (Vazyme C113-02) to generate a 3xFLAG::GFP::WAGO-4 fusion gene. The transgene was then integrated into the *C. elegans* genome using the mos1-mediated single copy insertion (MosSCI) method (Frokjaer-Jensen et al., 2014).

#### Dual sgRNA-directed *wago-4* deletion

Dual sgRNA-guided chromosome deletion was conducted as previously described (Chen et al., 2014). We manually searched for target sequences consisting of G(N)19NGG near the desired mutation sites in N2 genomic region. We replaced the *unc-119* target sequence in the pU6::unc-119 sgRNA vector with the desired target sequence using overlapped extension PCR with primer pairs sgRNA-1F: 5′-GAAGATCCGCTCATGGTCAAGTTTTAGAGCTAGAAATAGCAAGTTA-3′ and sgRNA-1R: 5′-CATGAGCGGATCTTCAAACATTTAGATTTGCAATTCAATTATATAG-3′; sgRNA-2F: 5′-GAATATTGCACCCGGTATTTGTTTTAGAGCTAGAAATAGCAAGTTA-3′ and sgRNA-2R: 5′-CCGGGTGCAATATTCAAACATTTAGATTTGCAATTCAATTATATAG-3′; sgRNA-3F: 5′-GTTCAGAGAAGGCGCAGTAAGTTTTAGAGCTAGAAATAGCAAGTTA-3′ and sgRNA-3R: 5′-GCGCCTTCTCTGAACAAACATTTAGATTTGCAATTCAATTATATAG-3′. Plasmid mixtures containing 50 ng/µl each of pU6::wago4 sgRNA1-3, 50 ng/µl Cas9 expression plasmid, 5 ng/µl pCFJ90 and 5 ng/µl pCFJ104 were co-injected into N2 animals. The deletion mutants were screened by PCR.

#### Quantitative RT-PCR

RNAs were isolated from indicated animals using a dounce homogenizer (pestle B) in TRIzol solution followed by DNase I digestion (Qiagen), as described previously (Guang et al., 2008). cDNAs were generated from RNAs using the iScript cDNA Synthesis Kit (Bio-Rad) according to the vendor’s protocol. qPCR was performed using a MyIQ2 real-time PCr system (Bio-Rad) with iQ SYBR Green Supermix (Bio-Rad). The primers that were used in qRT-PCR are provided in Supplemental Experimental Procedures. *eft-3* mRNA was used as an internal control for sample normalization. Data analysis was performed using a ΔΔCT approach.

#### RNA immunoprecipitation (RIP)

RIP experiments were performed as previously described with adult animals (Guang et al., 2008). Briefly, gravid adults were sonicated in sonication buffer (20 mM Tris-HCl [pH 7.5], 200 mM NaCl, 2.5 mM MgCl2, and 0.5% NP-40). The lysate was pre-cleared with protein G-agarose beads (Roche) and incubated with anti-FLAG M2 agarose beads (Sigma #A2220). The beads were washed extensively and 3xFLAG::GFP::WAGO-4 and its associated RNAs were eluted with 100 μg/ml 3xFLAG peptide (Sigma). The eluates were incubated with TRIzol reagent followed by isopropanol precipitation. WAGO-4-bound RNAs were quantified by real-time PCR.

#### Isolation and sequencing of WAGO-4-associated small RNAs

WAGO-4-associated siRNAs were isolated from gravid adults as described above. The precipitated RNAs were treated with calf intestinal alkaline phosphatase (CIAP, Invitrogen), re-extracted with TRIzol, and treated with T4 polynucleotide kinase (T4 PNK, New England Biolabs) in the presence of 1 mM ATP (Zhou et al., 2014).

WAGO-4-bound siRNAs were subjected to deep sequencing using an Illumina platform, according to the manufacturer’s instructions by the Beijing Genomics Institute (BGI Shenzhen). Small RNAs ranging from 18 to 30 nt were gel-purified and ligated to a 3′ adaptor (5′-pUCGUAUGCCGUCUUCUGCUUGidT-3′; p, phosphate; idT, inverted deoxythymidine) and a 5′ adaptor (5′-GUUCAGAGUUCUACAGUCCGACGAUC-3′). The ligation products were gel-purified, reverse transcribed, and amplified using Illumina’s sRNA primer set (5′-CAAGCAGAAGACGGCATACGA-3′; 5′-AATGATACGGCGACCACCGA-3′). The samples were then sequenced using an Illumina Hiseq platform.

The Illumina-generated raw reads were first filtered to remove adaptors, low-quality tags and contaminants to obtain clean reads at BGI Shenzhen. Clean reads ranging from 18 to 30 nt were mapped to the unmasked *C. elegans* genome and the transcriptome assembly WS243, respectively, using Bowtie2 with default parameters (Langmead and Salzberg, 2012). The number of reads targeting each transcript was counted using custom Perl scripts and displayed by IGV (Thorvaldsdottir et al., 2013). The number of total reads mapped to the genome minus the number of total reads corresponding to sense rRNA transcripts (5S, 5.8S, 18S, and 26S) was used as the normalization number to exclude the possible degradation fragments of sense rRNAs.

#### Statistics

Bar graphs with error bars were presented with mean and s.d. All the experiments were conducted with independent *C. elegans* animals for indicated times. Statistical analysis was performed with two-tailed Student’s t-test.

#### Data availability

Argonaute-associated small RNAs from published data were re-analyzed. CSR-1 IP in wild type animals and in *cde-1* mutants, GSE17787 (van Wolfswinkel et al., 2009); WAGO-1 IP, SRR030711 (Gu et al., 2009); *glp-4(bn2)*, GSM455394 (Gu et al., 2009); WAGO-4 IP, GSE112475.

All other data and materials are available upon request.

## Acknowledgments

We are grateful to Dr. Scott Kennedy and members of the Guang lab for their comments. We are grateful to Dr. Ho Yi Mak, the International *C. elegans* Gene Knockout Consortium, and the National Bioresource Project for providing the strains. Some strains were provided by the CGC, which is funded by NIH Office of Research Infrastructure Programs (P40 OD010440). This work was supported by grants from the Chinese Ministry of Science and Technology (2017YFA0102903), the National Natural Science Foundation of China (Nos. 81501329, 31671346, and 91640110), Anhui Natural Science Foundation (No. 1608085MC50), and the Major/Innovative Program of Development Foundation of Hefei Center for Physical Science and Technology (2017FXZY005).

## Author Contributions

F. X., X. F. and S. G. designed the experiments, F. X., X. F., X. C., C. W., Q. Y., T. X., and M. H. performed experiments. F. X., X. F. and S. G. analyzed the data. F. X., X. F. and S. G. wrote the manuscript. All authors have discussed the manuscript.

## Additional Information

The authors declare no competing financial interests.

## Supplemental Information

Supplemental Information includes Supplemental Experimental Procedures, five figures (Figures S1–S5), and one table.

Table S1. WAGO-4-associated siRNAs targeted genes, Related to Figure 5

---

**Figure S1.**
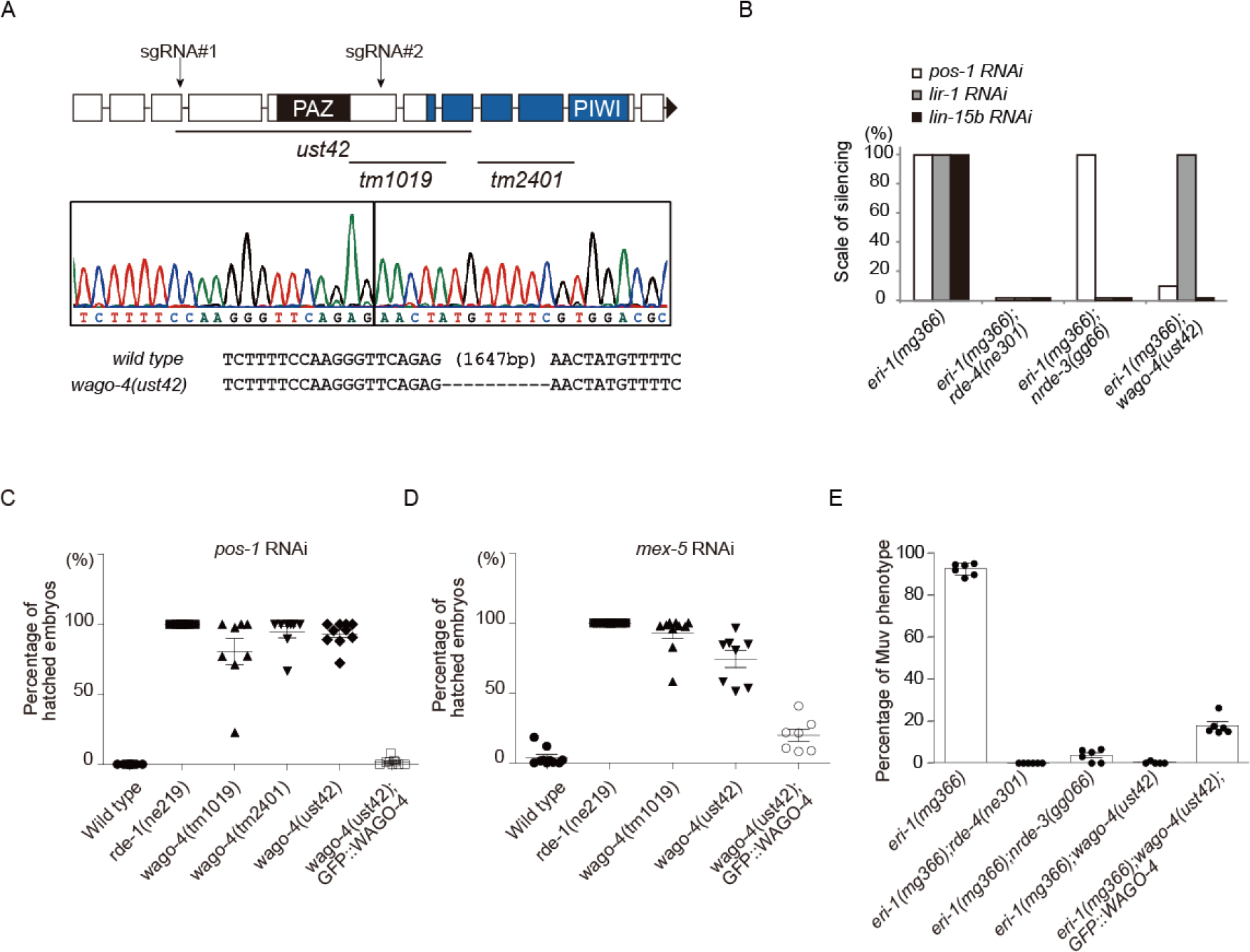
WAGO-4 is involved in RNAi targeting germline-expressed genes, Related to Figure 1. (a) Schematic of WAGO-4 and the *ust42* allele generated by CRISPR/Cas9 technology. (b) Feeding RNAi analysis of indicated animals. *pos-1* RNAi induces embryonic lethality; *lir-1* RNAi elicits growth arrest at the L2 stage; *lin-15b* RNAi results in the muv phenotype in F1 animals. n>100 animals. (c, d, e) The indicated animals were treated with *pos-1, mex-5*, or *lin-15b* dsRNA, and the phenotypes were scored.

**Figure S2.**
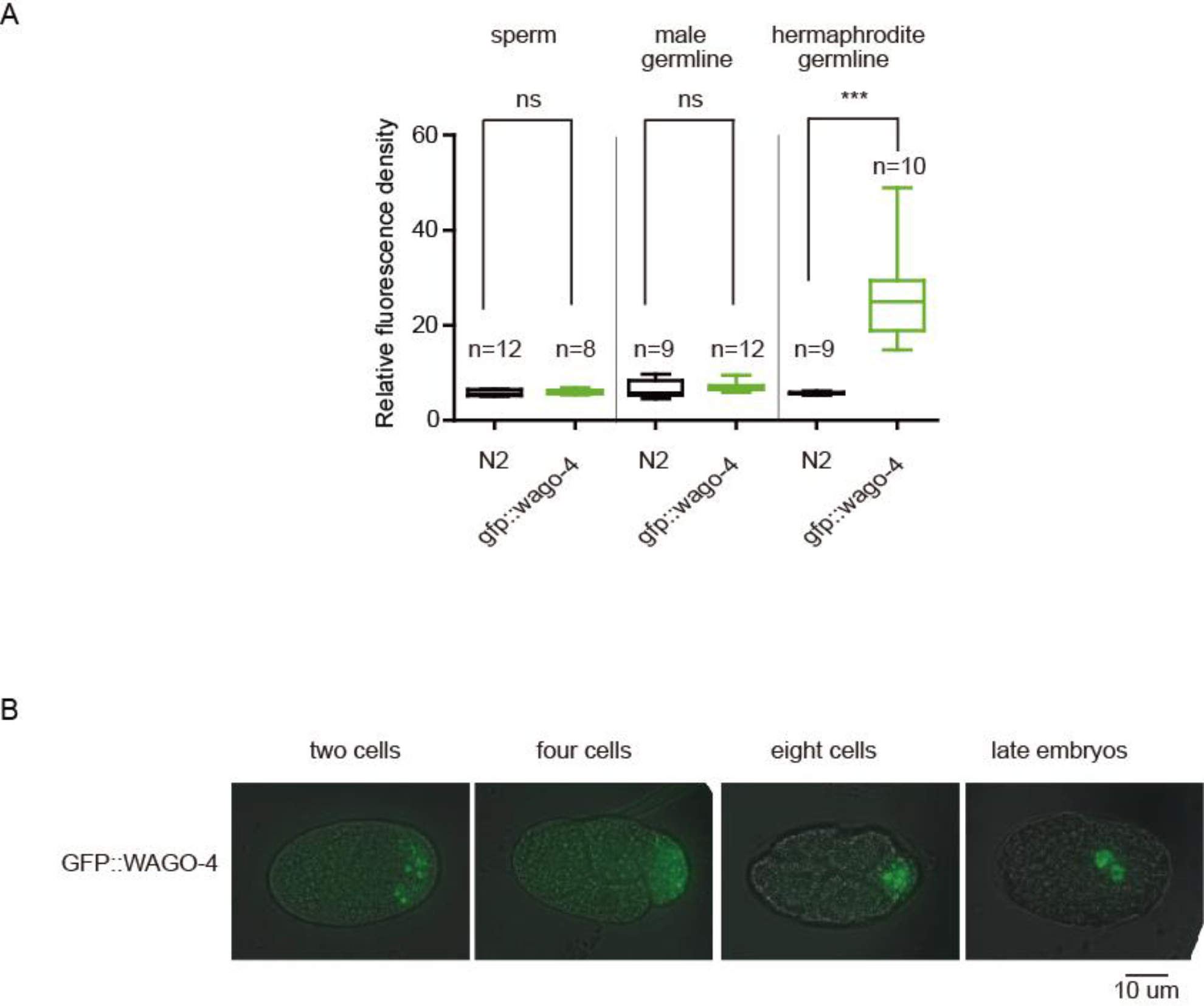
WAGO-4 is a germline-expressed Argonaute protein, Related to Figure 2. (a) Quantification of fluorescence density of *GFP::WAGO-4* animals, measured by ImageJ. n indicates the number of animals quantified. ***p<0.001; ns, not significant. (b) Images of GFP::WAGO-4 at indicated developmental stages.

**Figure S3.**
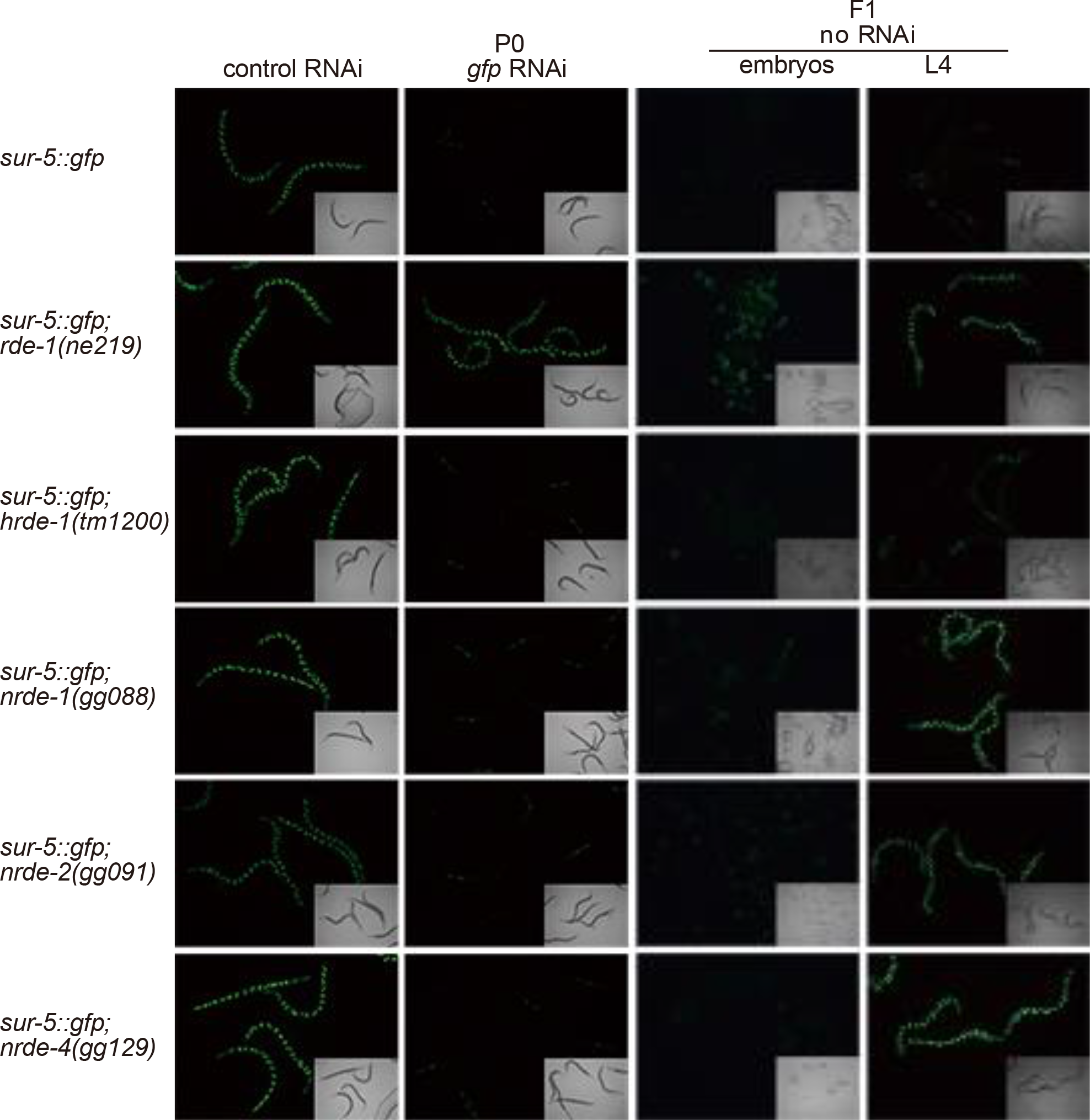
WAGO-4 promotes RNAi inheritance, Related to Figure 3. Images of indicated animals after *gfp* RNAi. F1 embryos were transferred to HT115 control bacteria in the absence of *gfp* dsRNA.

**Figure S4.**
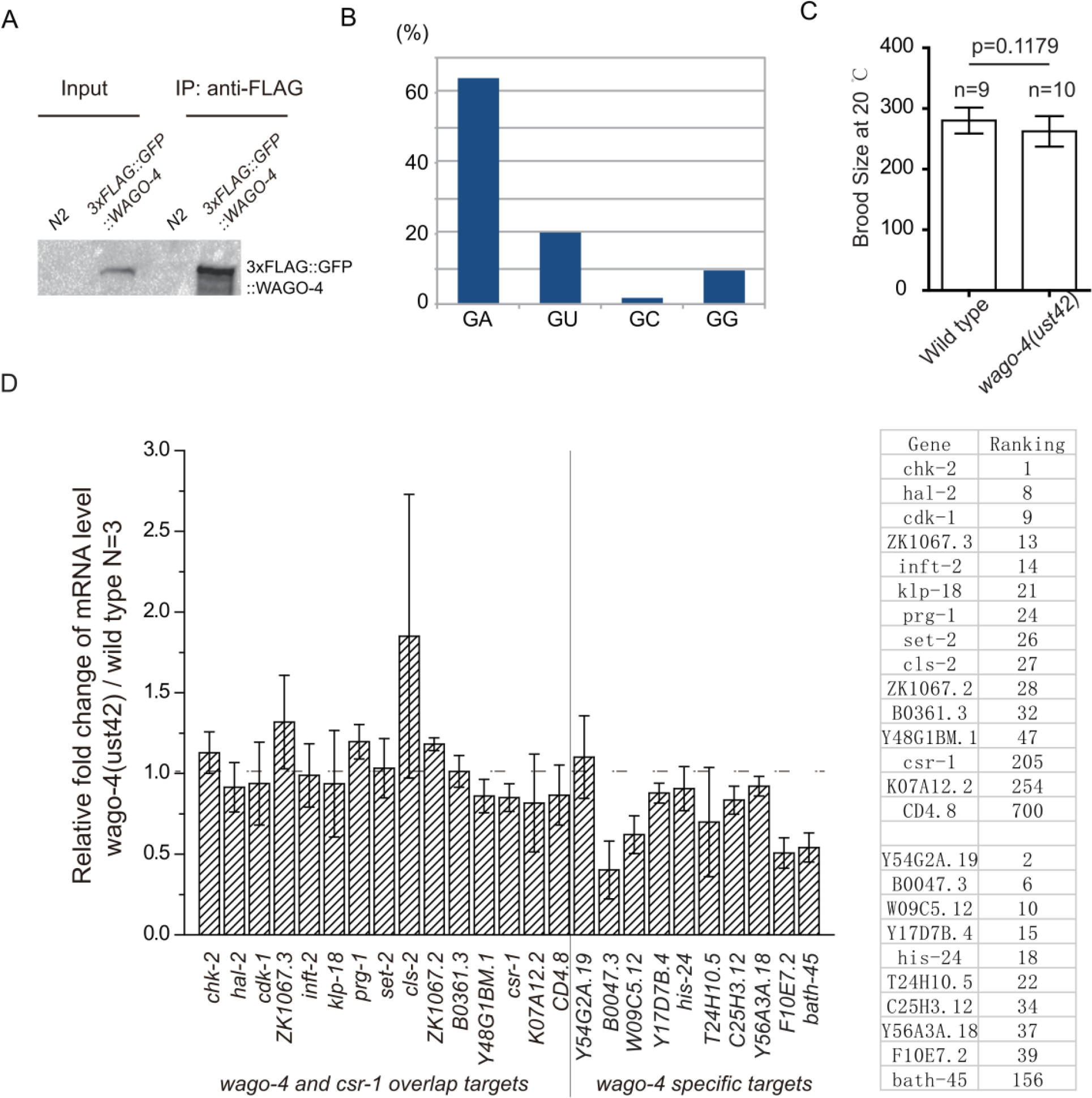
WAGO-4 binds to 22G-RNAs, Related to Figure 5. (a) 3xFLAG::GFP::WAGO-4 was immunoprecipitated from indicated animals and blotted by anti-FLAG antibody. (b) Distribution of the first two nucleotides of WAGO-4 siRNAs. (c) Brood size of indicated animals at 20℃. n indicates the number of animals scored. mean±s.d. (d) Relative mRNA levels of endogenous WAGO-4 22G-RNA targets in *wago-4(ust42)* animals. Right panel indicates the ranking of WAGO-4 22G-RNA targets. n=3, mean±s.d.

**Figure S5.**
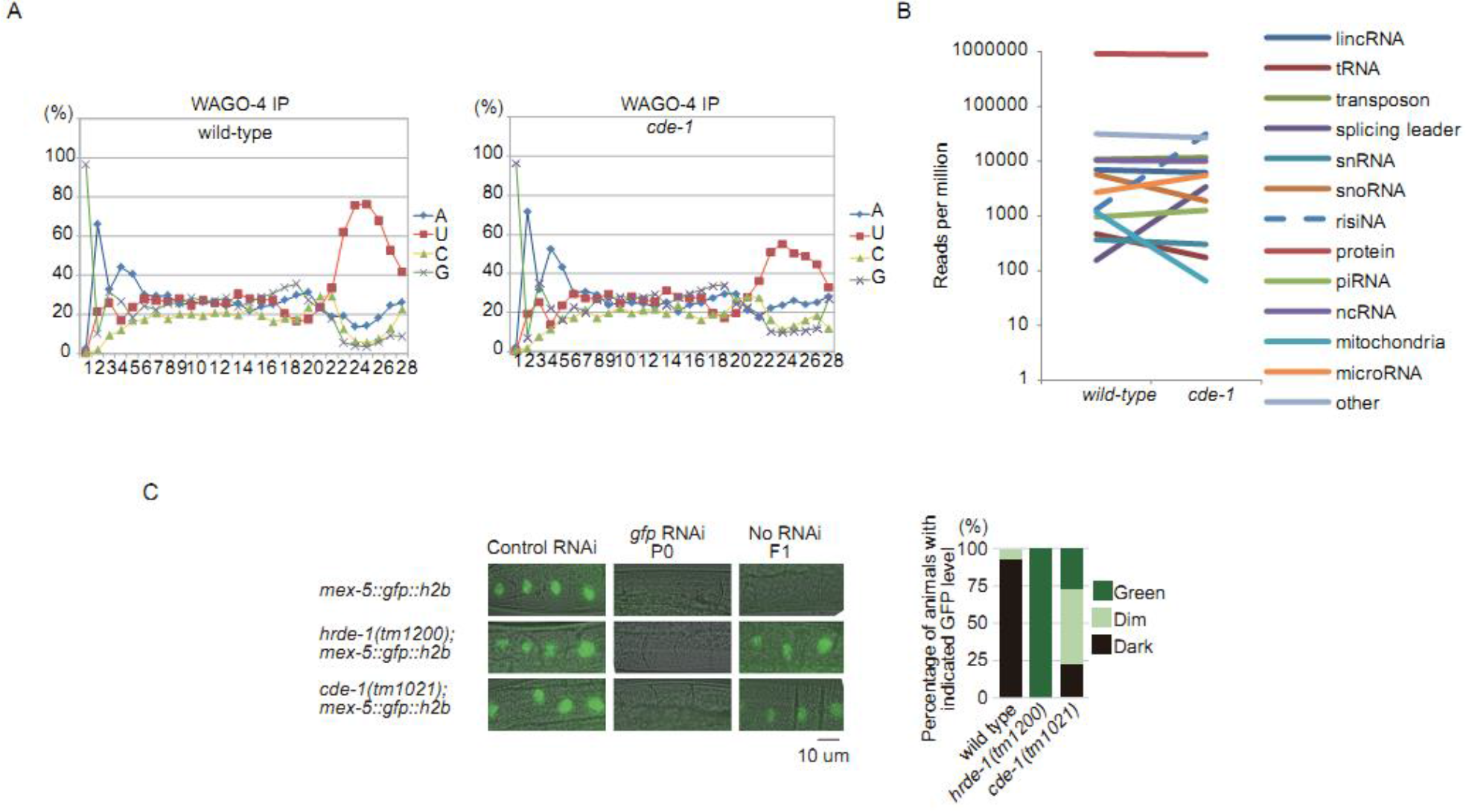
CDE-1 uridylates WAGO-4-associated 22G-RNAs, Related to Figure 6. (a) Nucleotide distribution of WAGO-4-associated 22G-RNAs in wild type animals (left panel) and in *cde-1* mutants (right panel). (b) Small RNA categories of WAGO-4-associated 22G-RNAs in wild type animals and *cde-1* mutants. (c) Images of indicated animals after *gfp* RNAi. F1 animals were grown on HT115 control bacteria. The *gfp* levels were scored in the right panel.

## Supplemental Experimental Procedures

Strains used in this study
| strains | genotype |
| --- | --- |
|  | *hrde-1(tm1200)* |
|  | *wago-4(tm1019)* |
|  | *wago-4(tm2401)* |
|  | *wago-4(ust42)* |
|  | *eri-1(mg366)* |
| (YY380) | *eri-1(mg366);rde-4(ne301)* |
| (YY189) | *eri-1(mg366);nrde-3(gg066)* |
| (SHG503) | *eri-1(mg366);wago-4(ust42)* |
| (YY513) | *pkIs32[pie-1p::h2b::gfp]* |
| (YY528) | *hrde-1(tm1200); pkIs32[pie-1p::h2b::gfp]* |
| (SHG480) | *wago-4(tm1019); pkIs32[pie-1p::h2b::gfp]* |
| (SHG481) | *wago-4(ust42); pkIs32[pie-1p::h2b::gfp]* |
| (SHG482) | *wago-4(tm2401); pkIs32[pie-1p::h2b::gfp]* |
| (MH1870) | *kuIs54[sur-5::gfp]* |
| (YY518) | *nrde-1(gg088);kuIs54[sur-5::gfp]* |
| (SHG039) | *nrde-2(gg091);kuIs54[sur-5::gfp]* |
| (YY521) | *nrde-4(gg129); kuIs54[sur-5::gfp]* |
| (YY548) | *hrde-1(tm1200);kuIs54[sur-5::gfp]* |
| (SHG483) | *rde-1(ne219); kuIs54[sur-5::gfp]* |
| (SHG485) | *wago-4(ust42); kuIs54[sur-5::gfp]* |
| (SHG486) | *wago-4(tm2401);kuIs54[sur-5::gfp]* |
| (SHG487) | *ustIs25[wago-4p::3xFLAG::GFP::wago-4]* |
| (SHG495) | *cde-1(tm1021);ustIs25[wago-4p::3xFLAG::GFP::wago-4]* |
| (SHG366) | *ustIs45[mex-5p::GFP::his-58::tbb-2utr]* |
| (SHG501) | *hrde-1(tm1200);mex-5p::GFP::his-58::tbb-2utr* |
| (SHG502) | *cde-1(tm1021);mex-5p::GFP::his-58::tbb-2utr* |
| (SHG341) | *ustIs25[wago-4p::3xFLAG::GFP::wago-4];hjSi396[dpy-30p::mRuby::pgl-1]* |
| (SHG506) | *ustIs25[wago-4p::3xFLAG::GFP::wago-4];ltIs37[(pAA64) pie-1p::mCherry::his-58 + unc-119(+)].* |

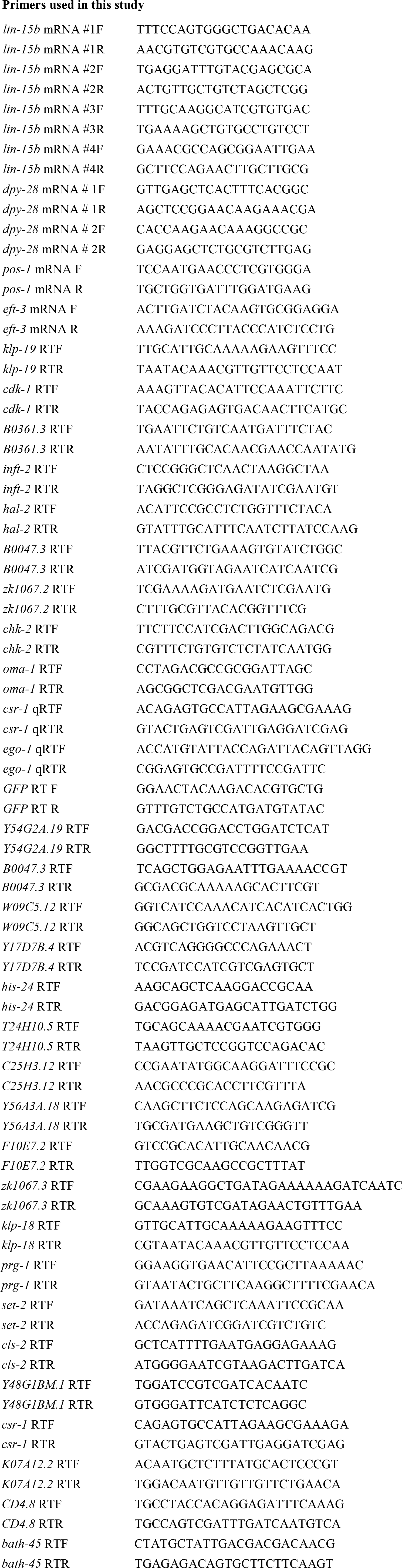

**Table S1.**
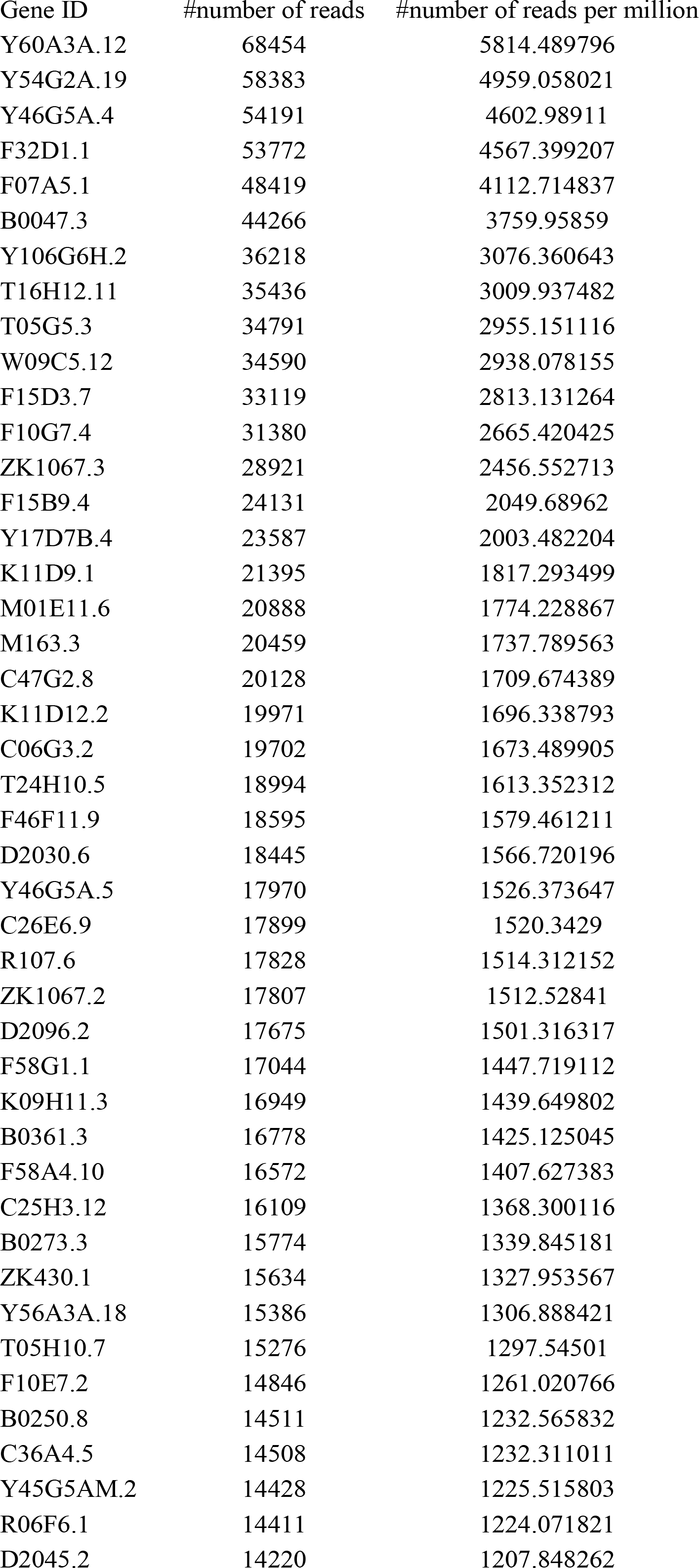
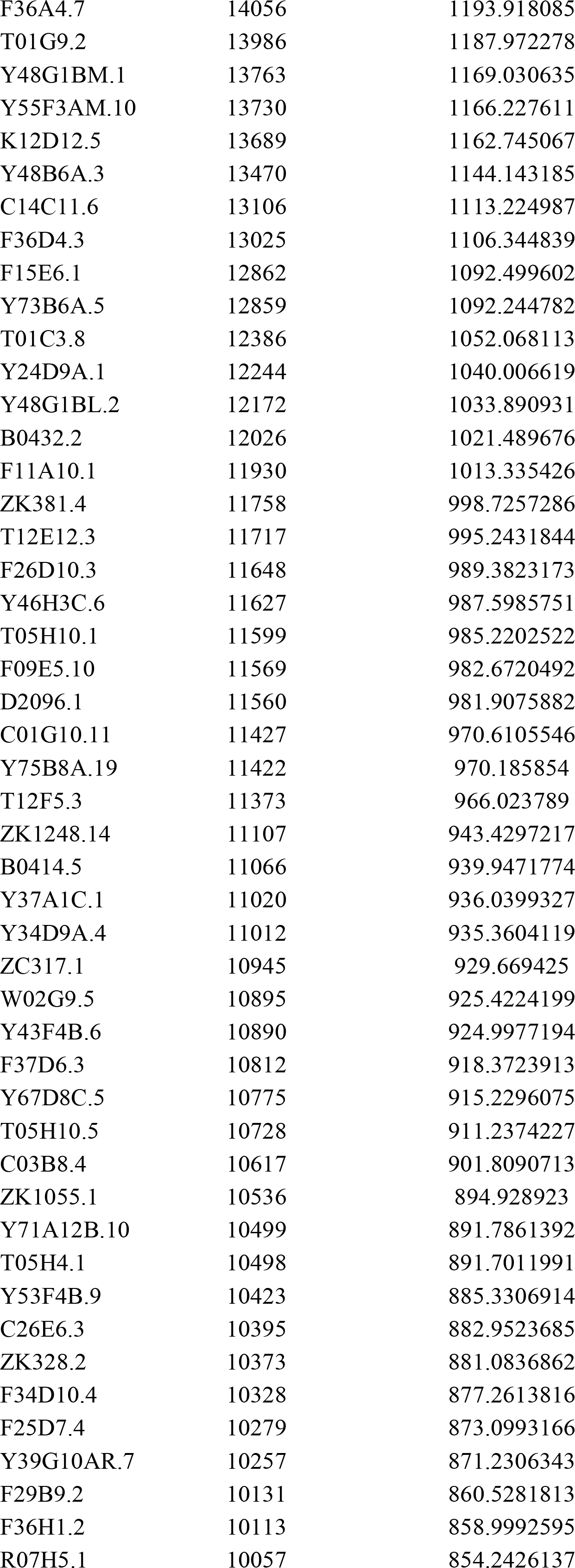
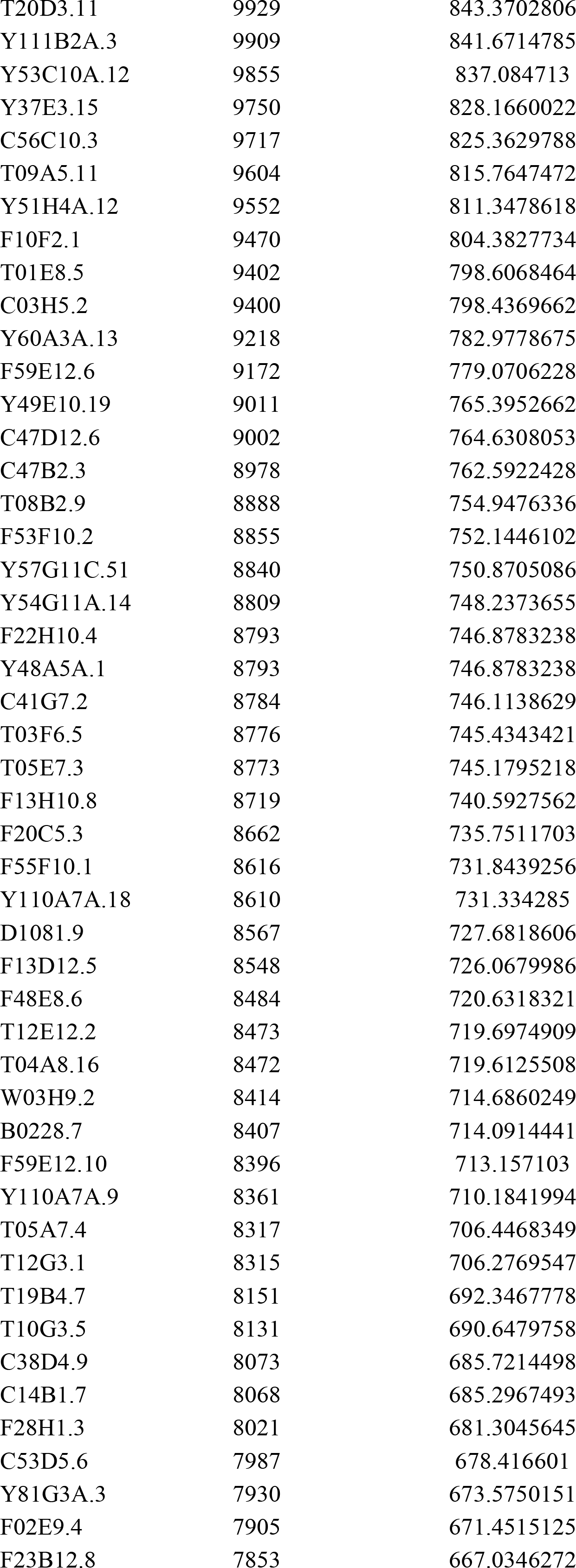
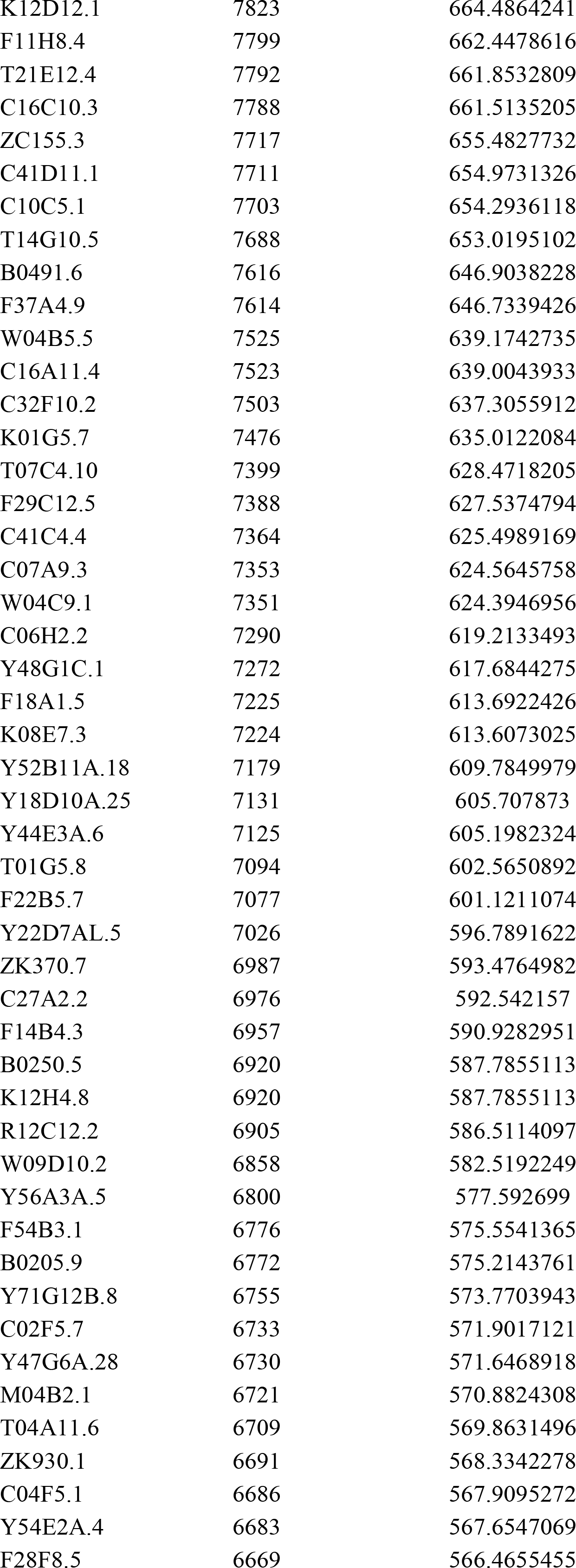
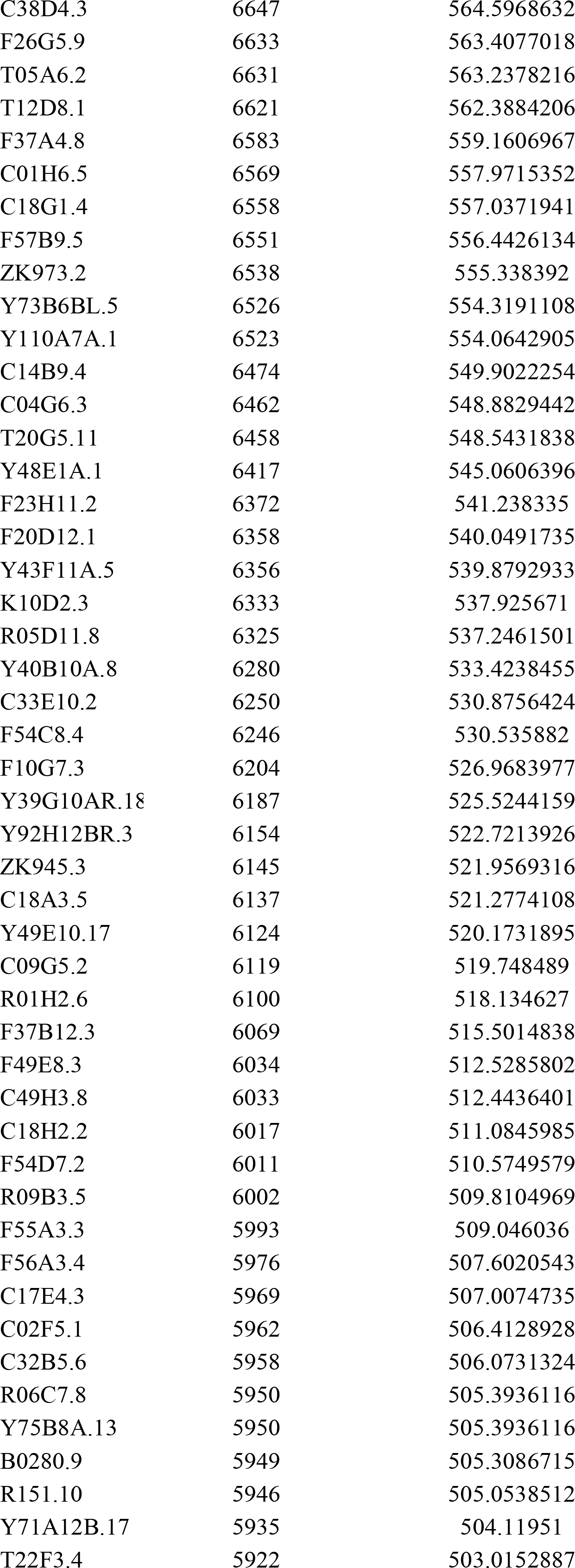
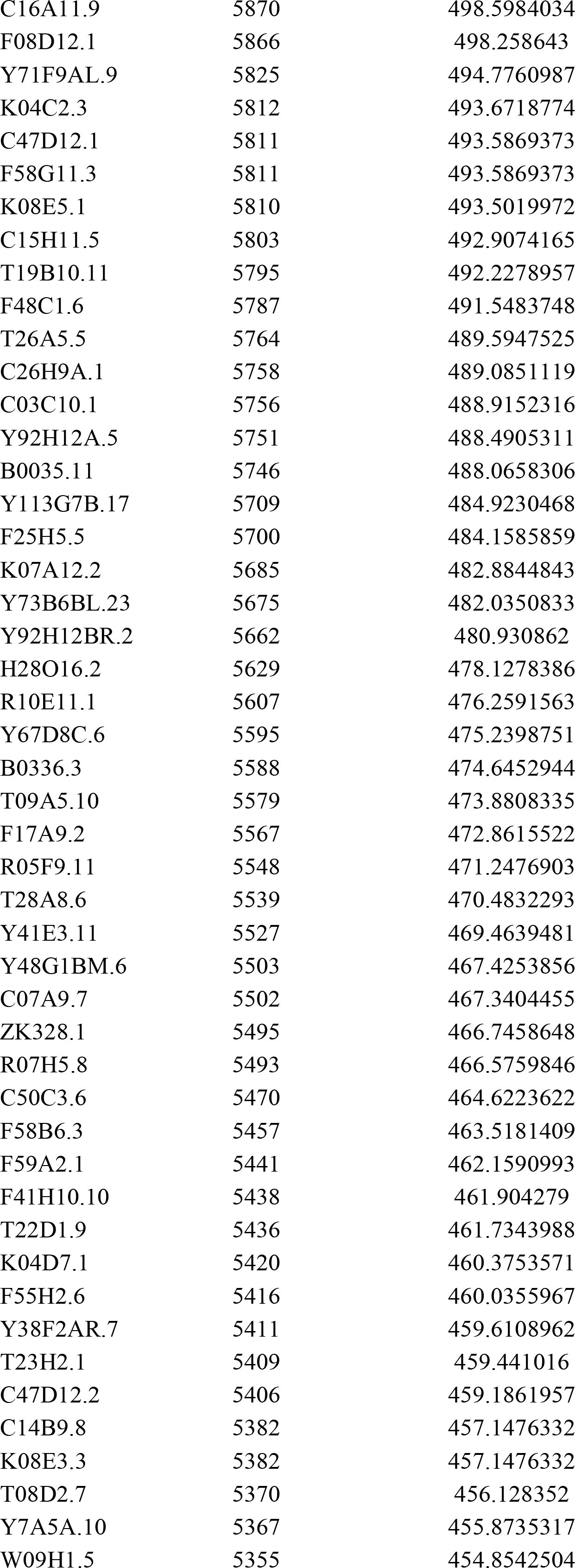
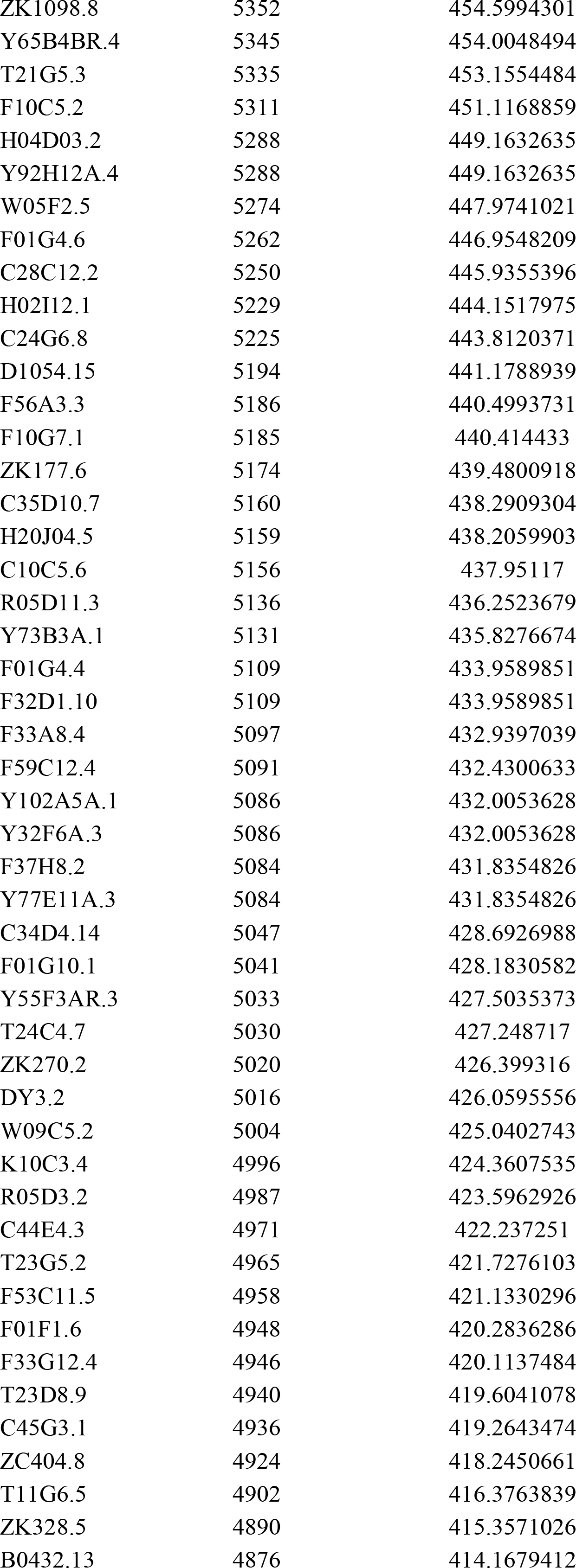
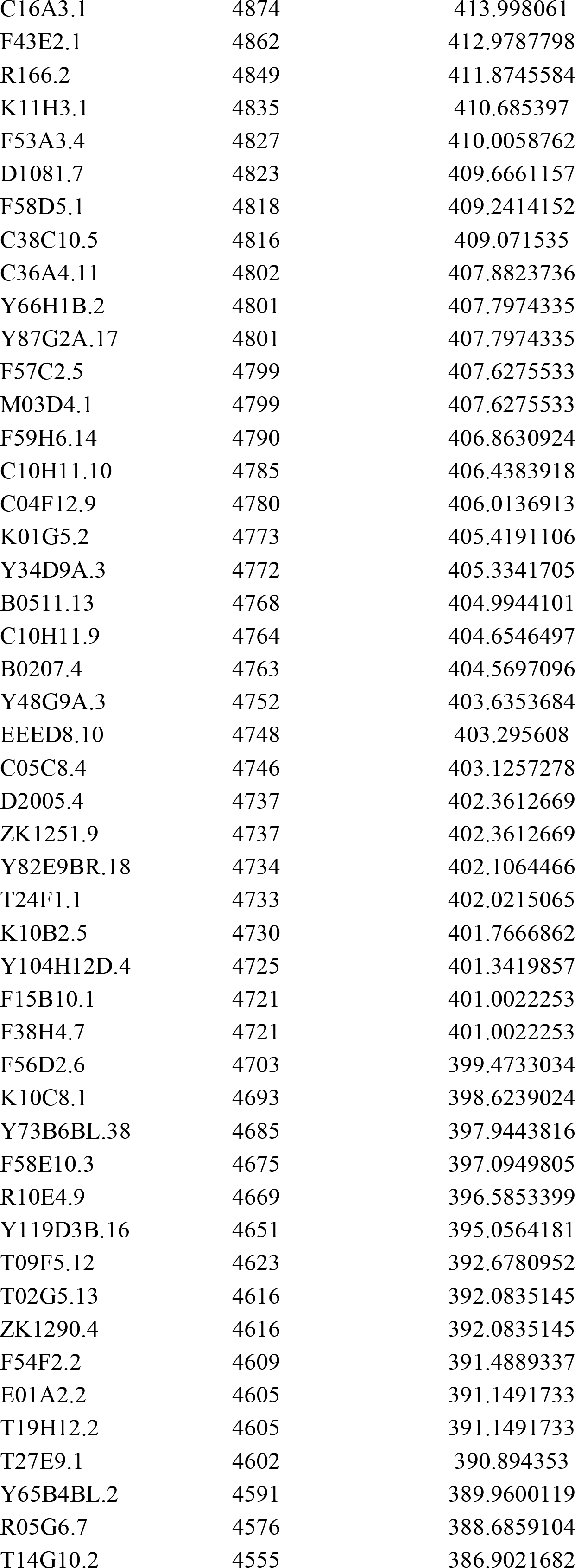
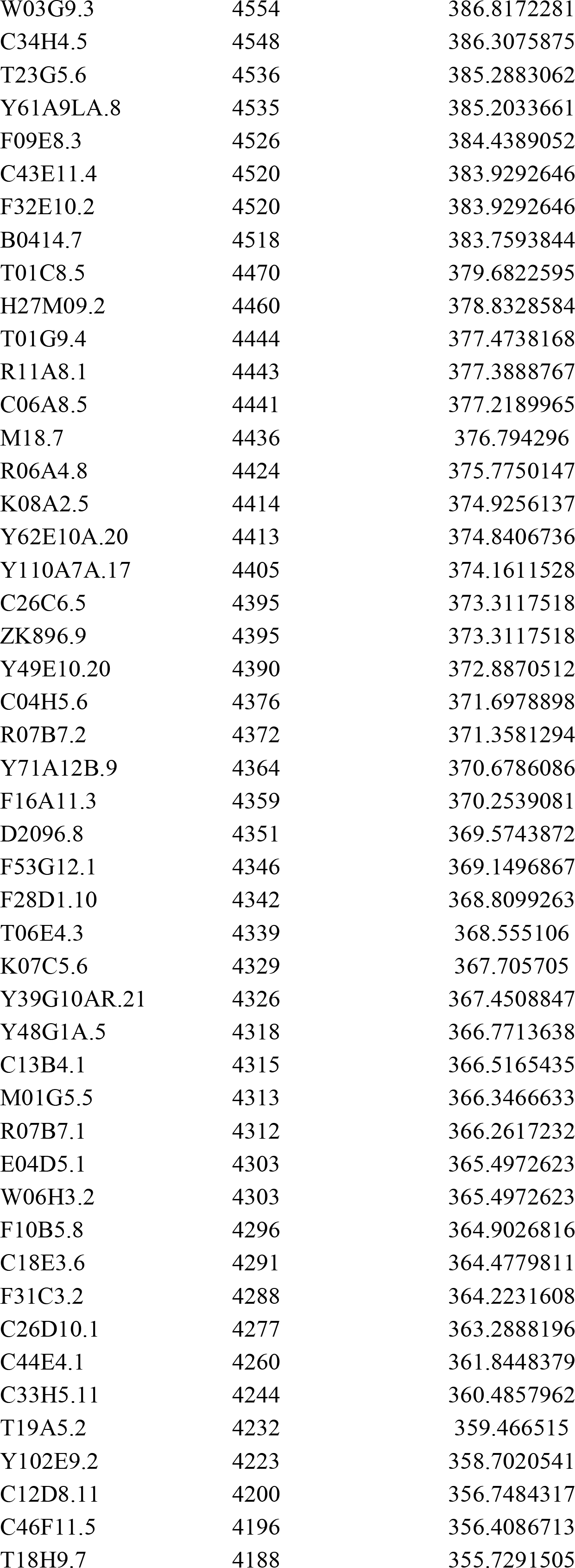
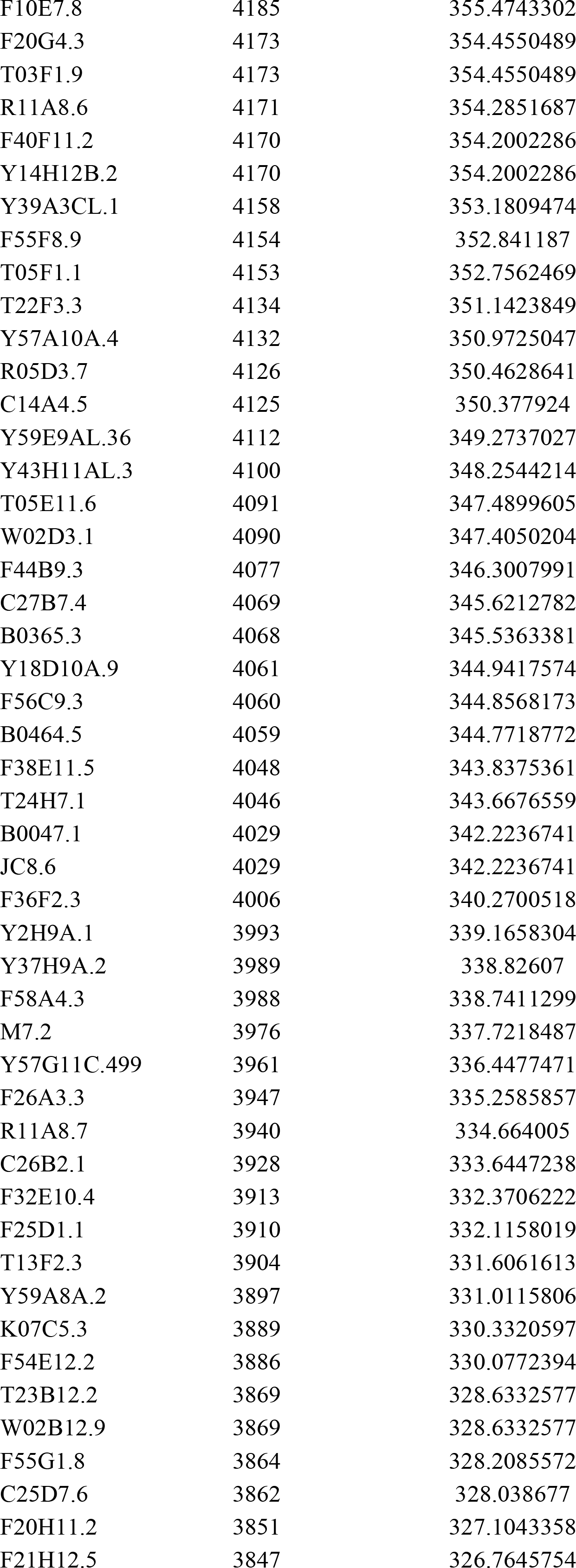
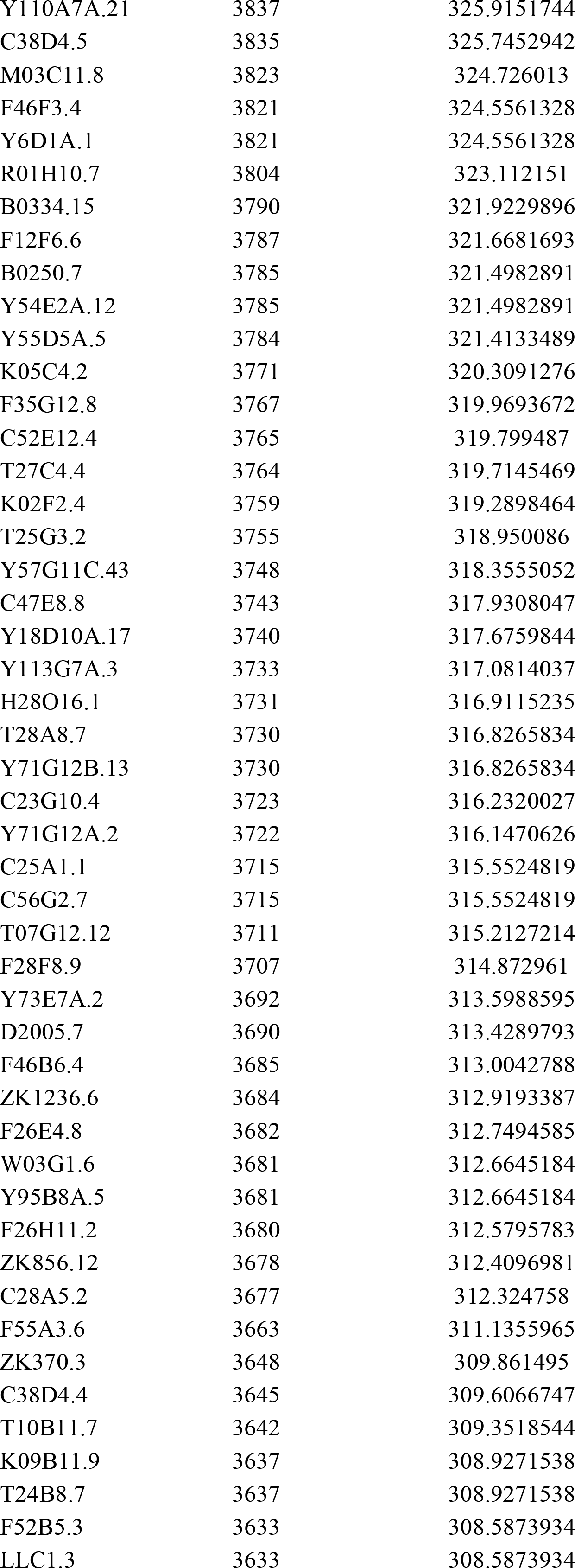
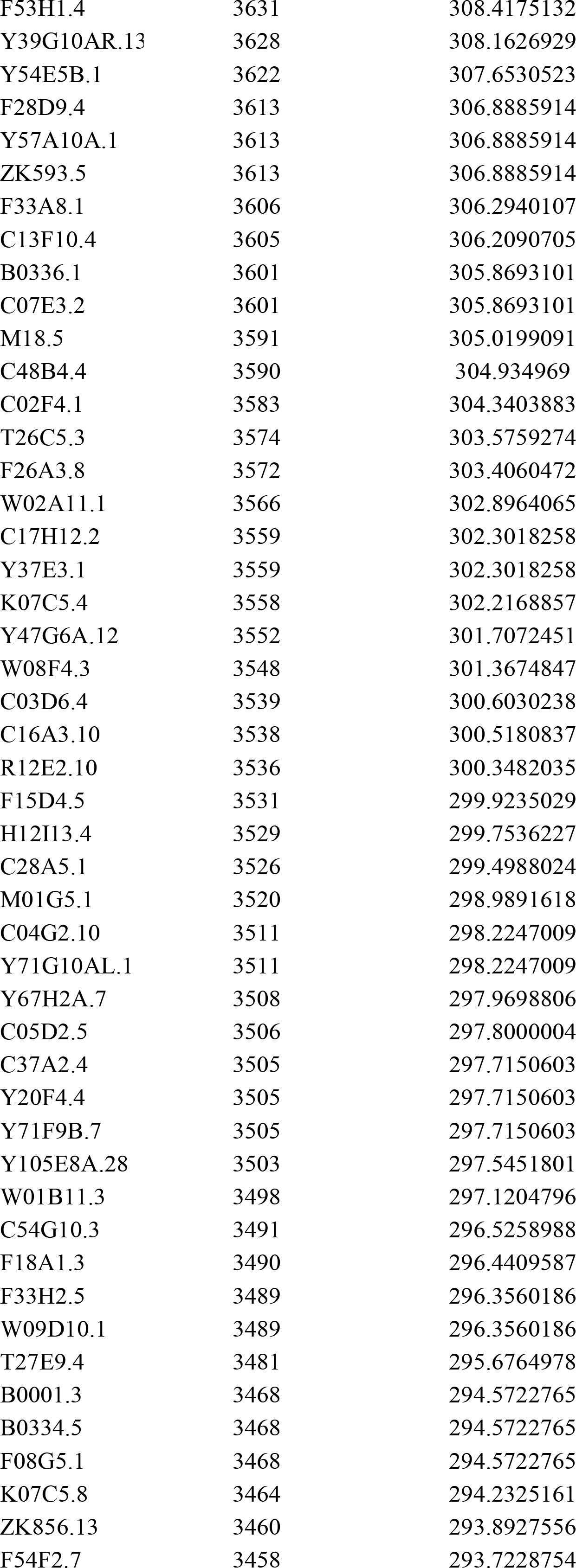
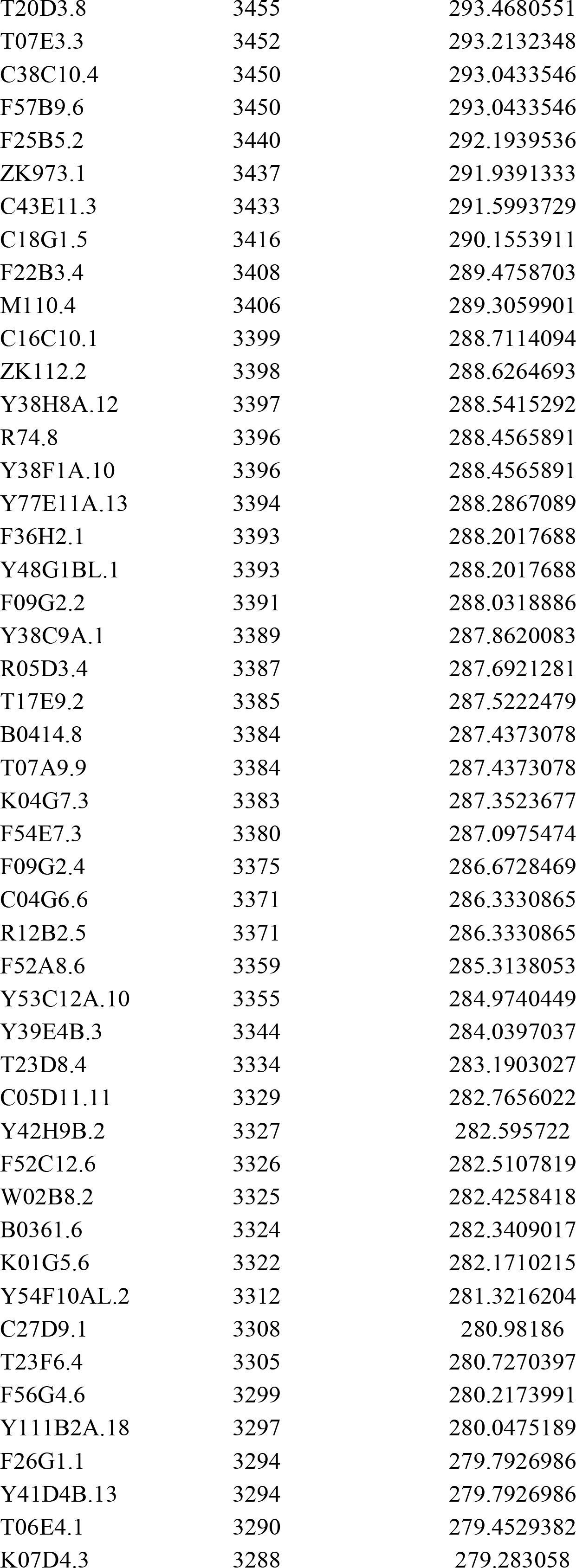
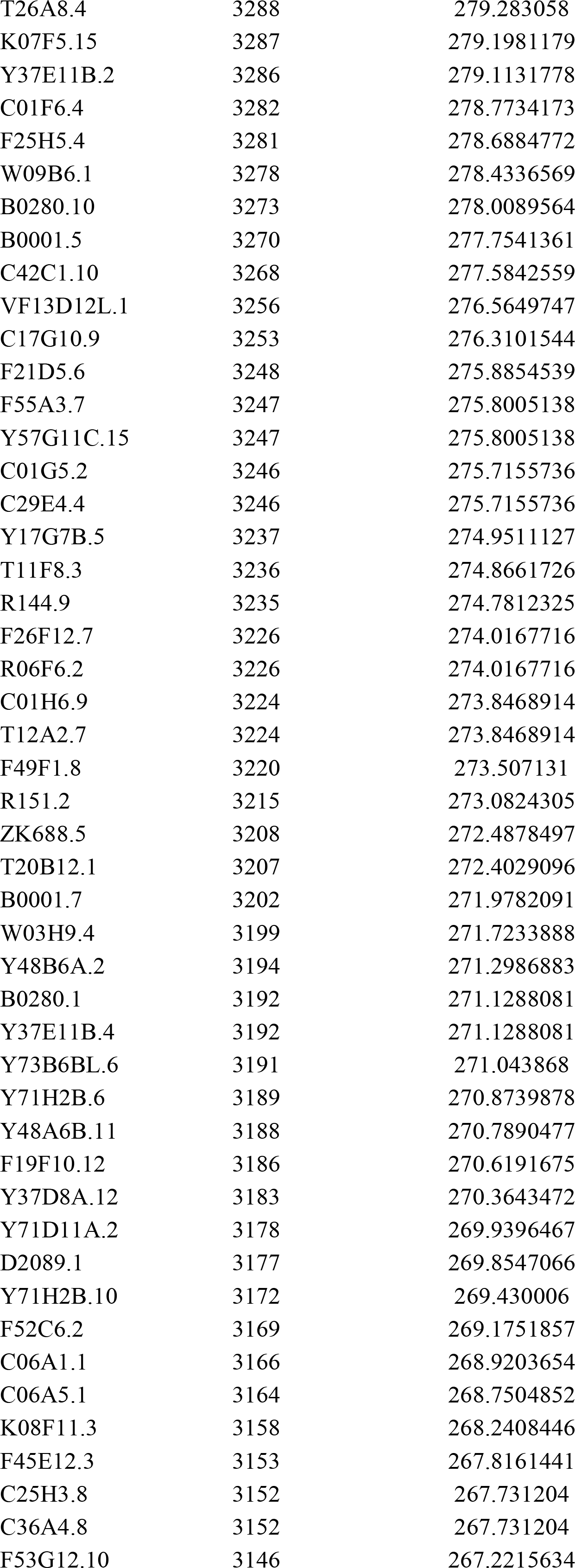
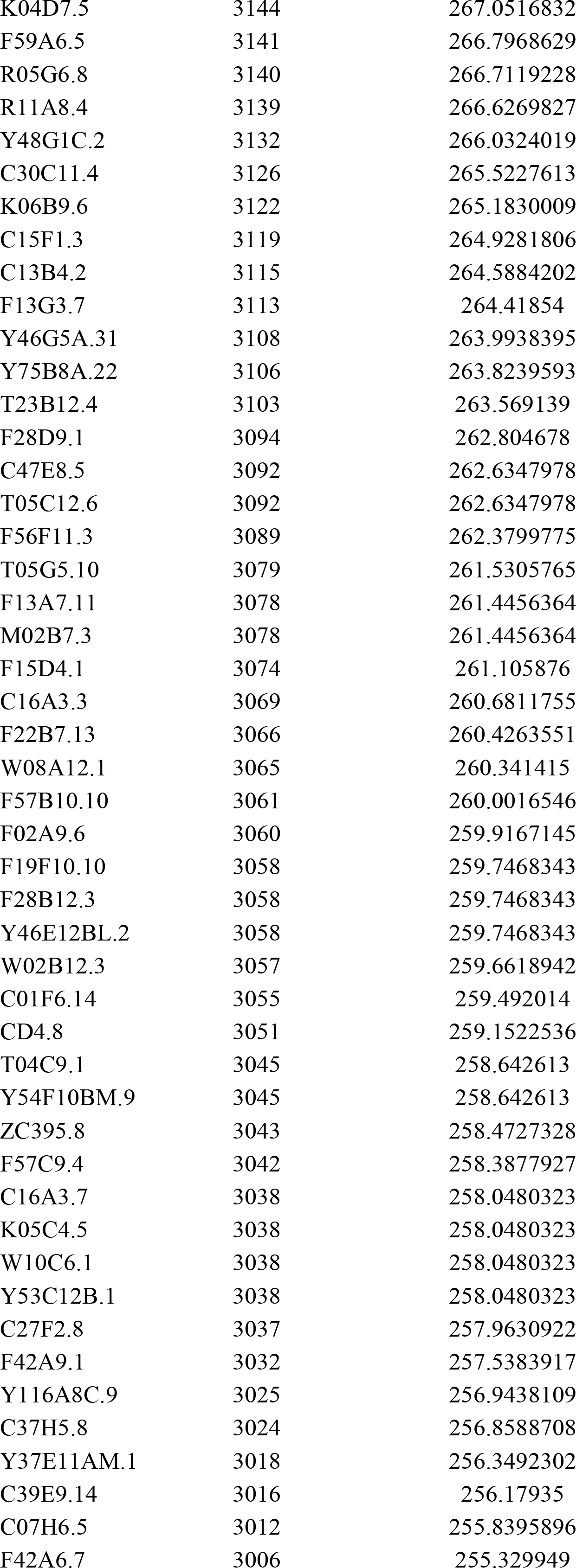
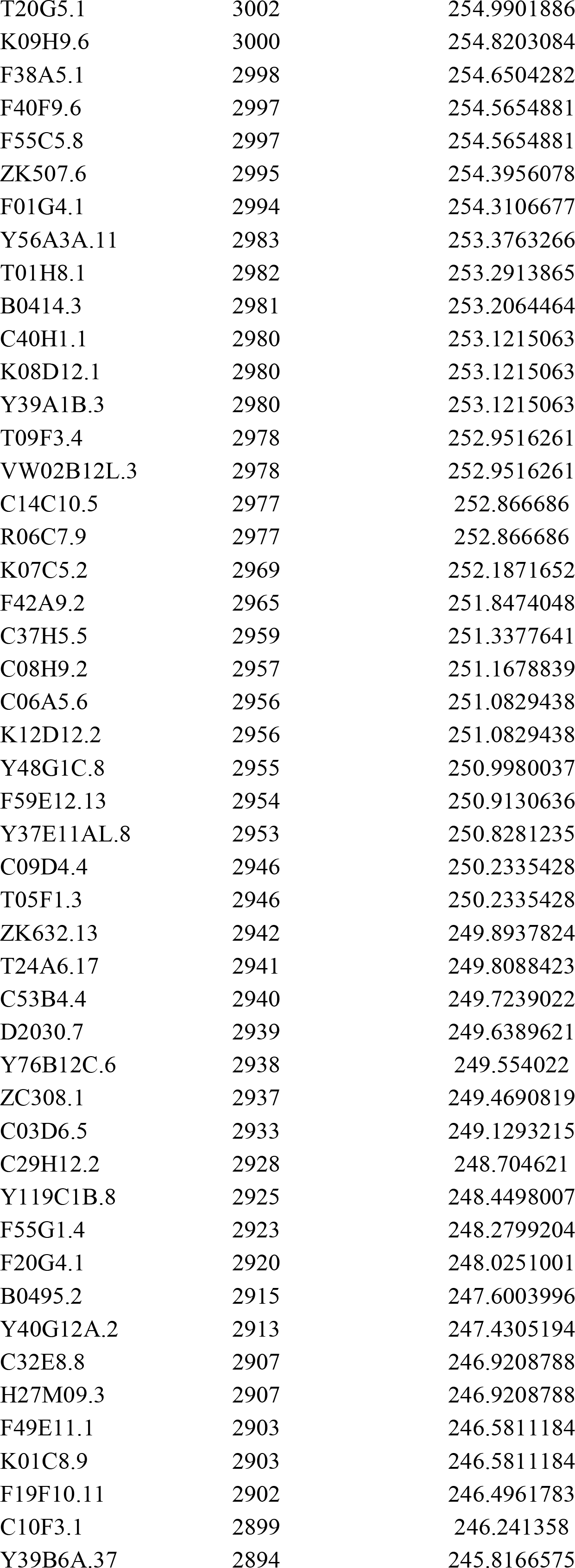
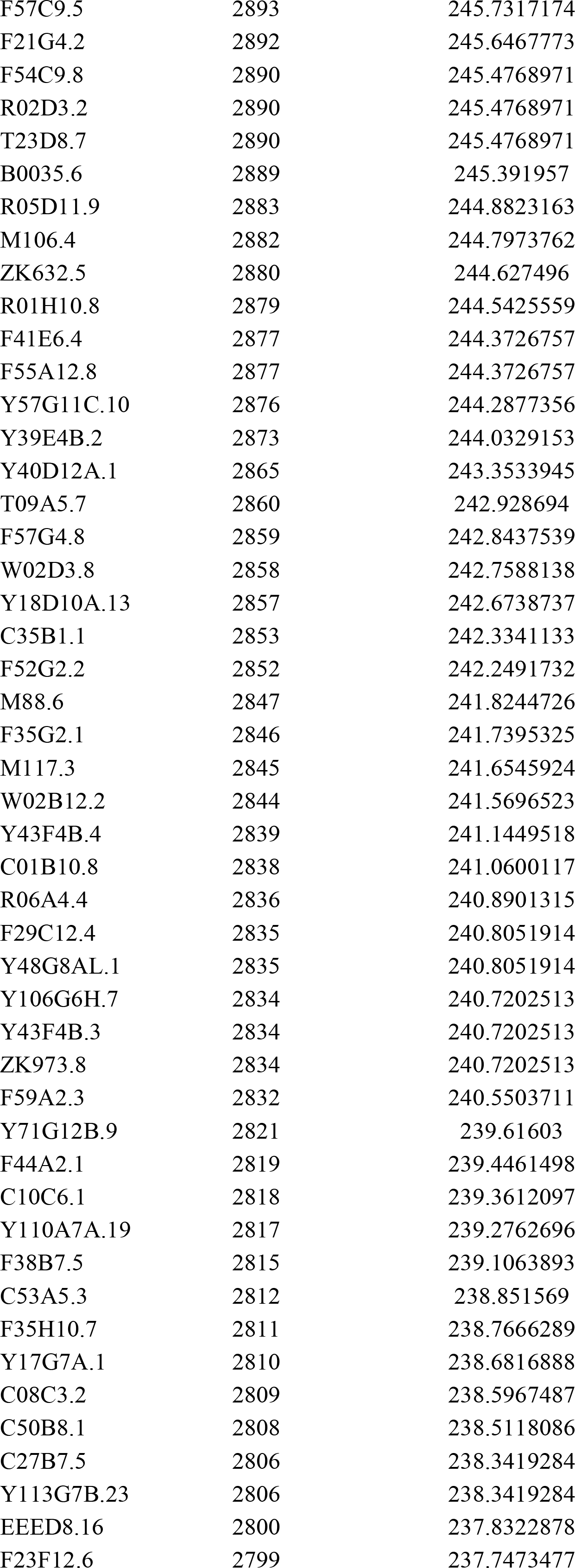
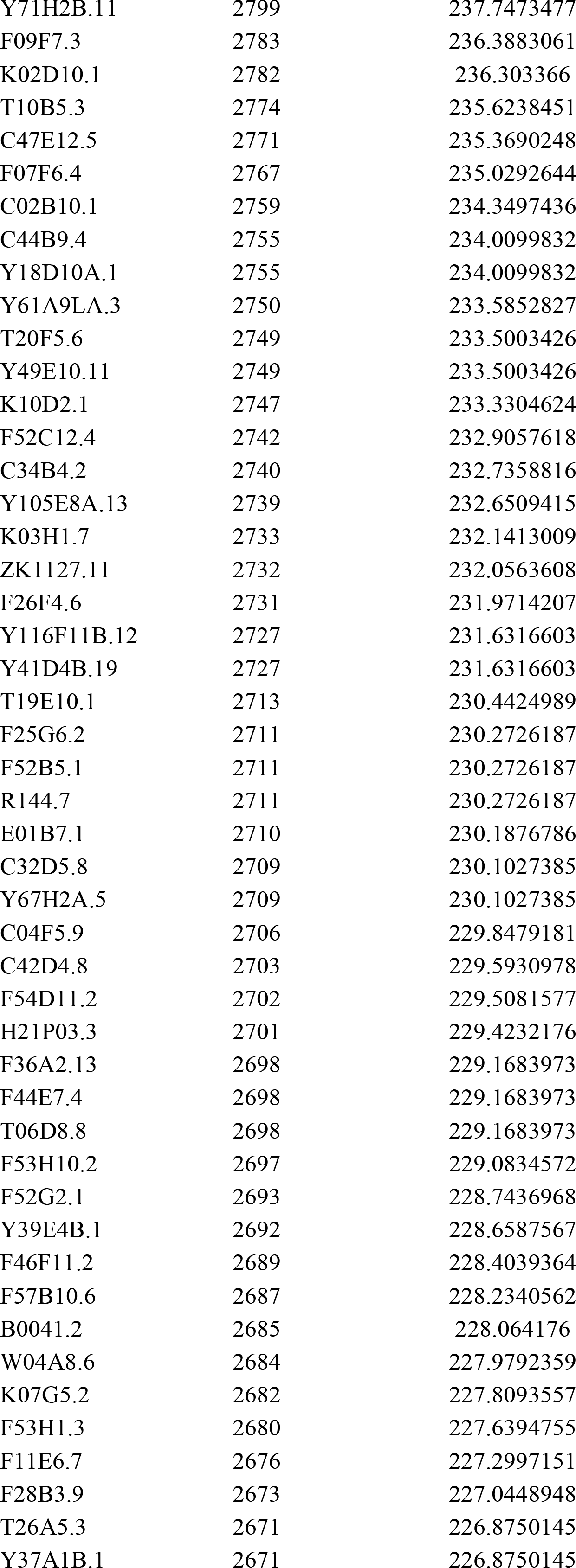
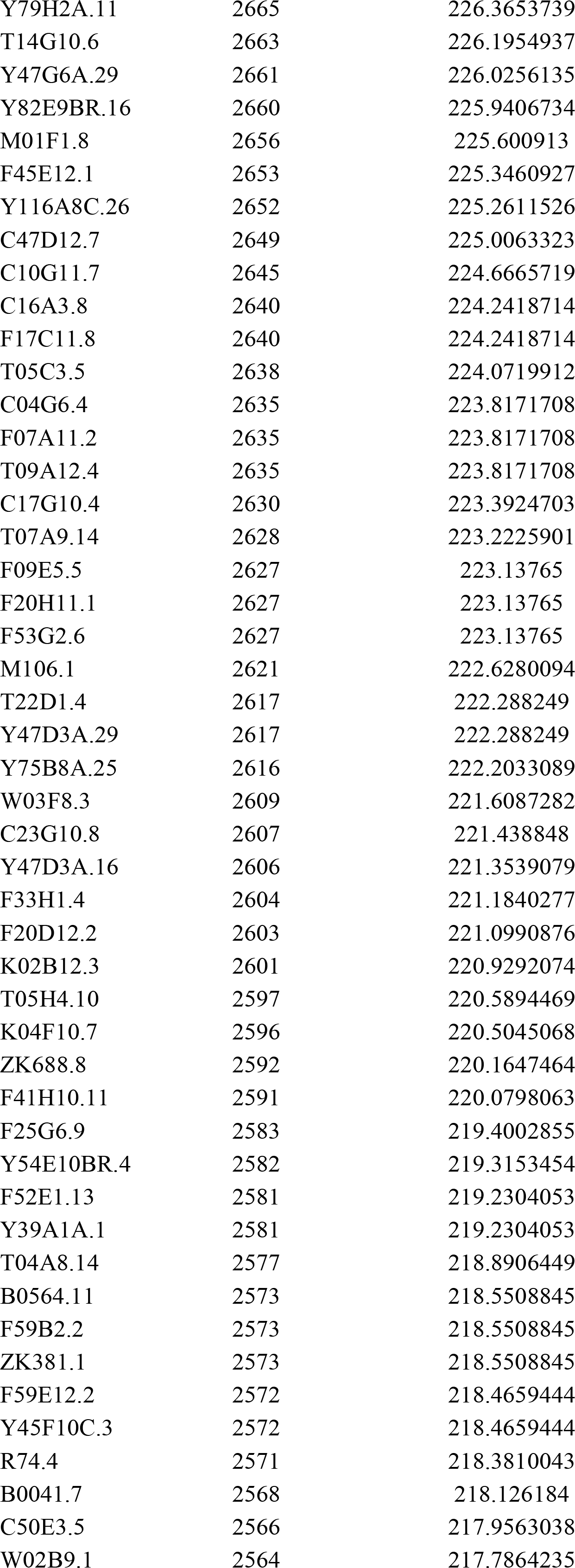
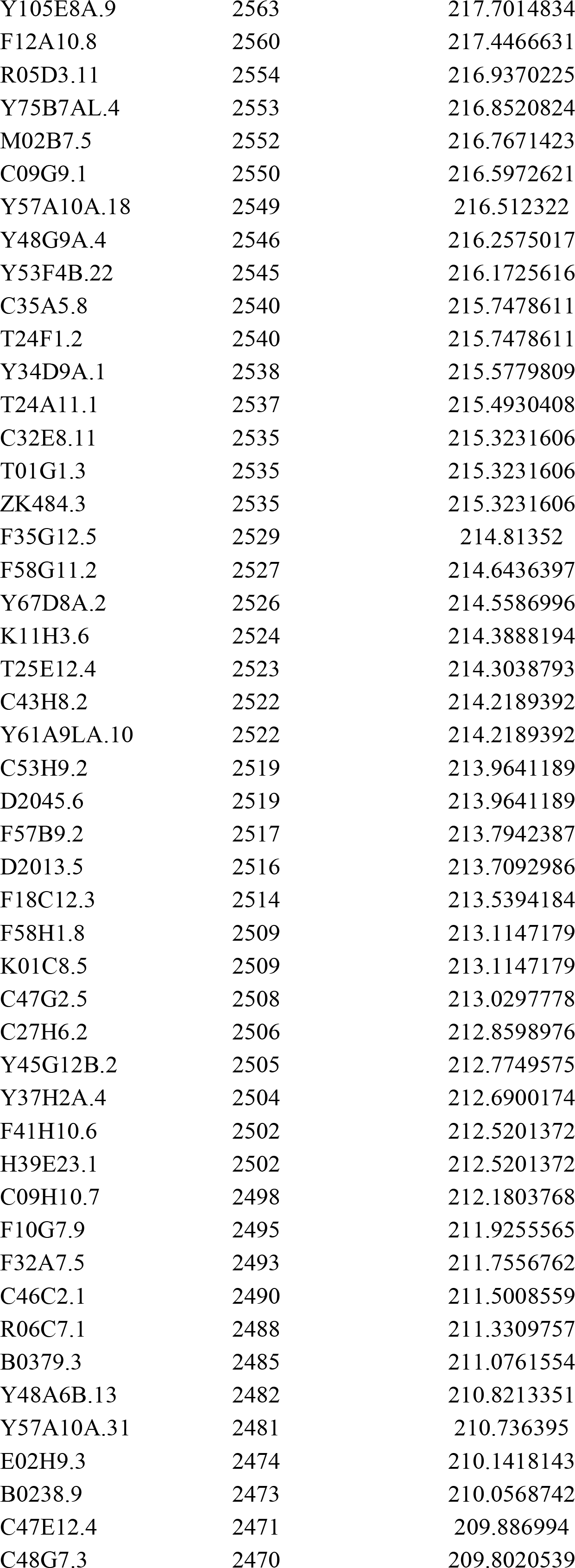
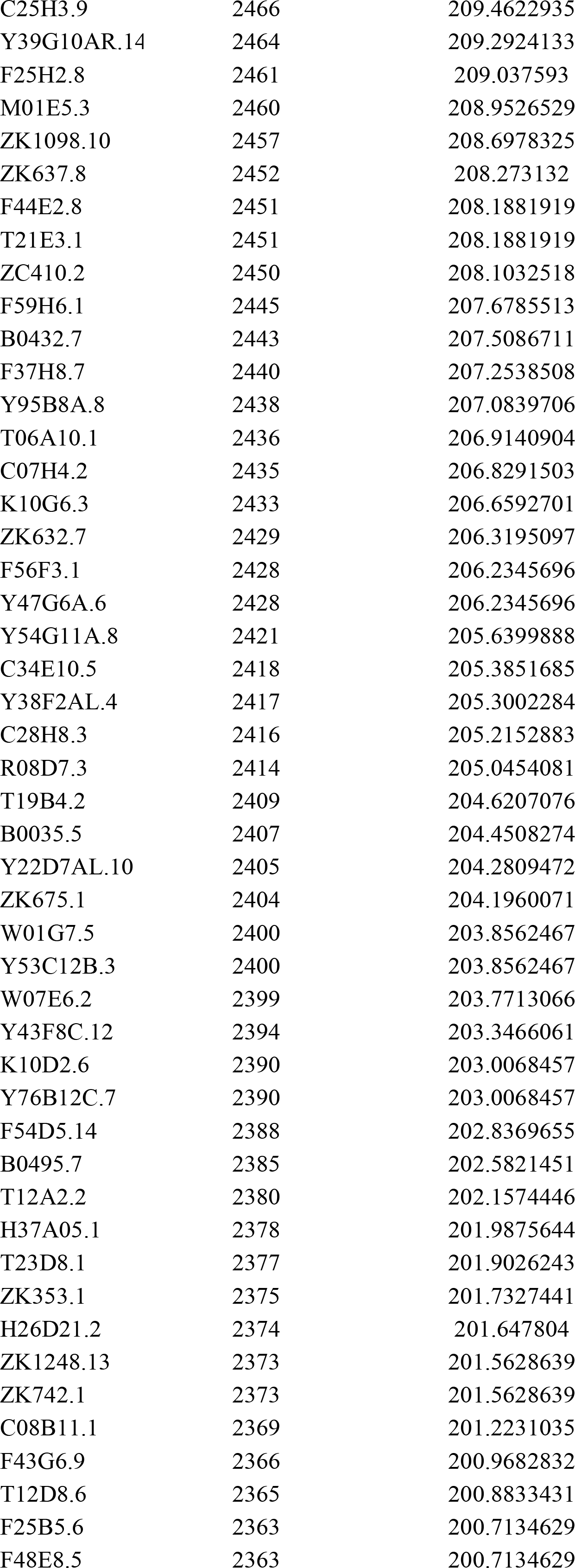
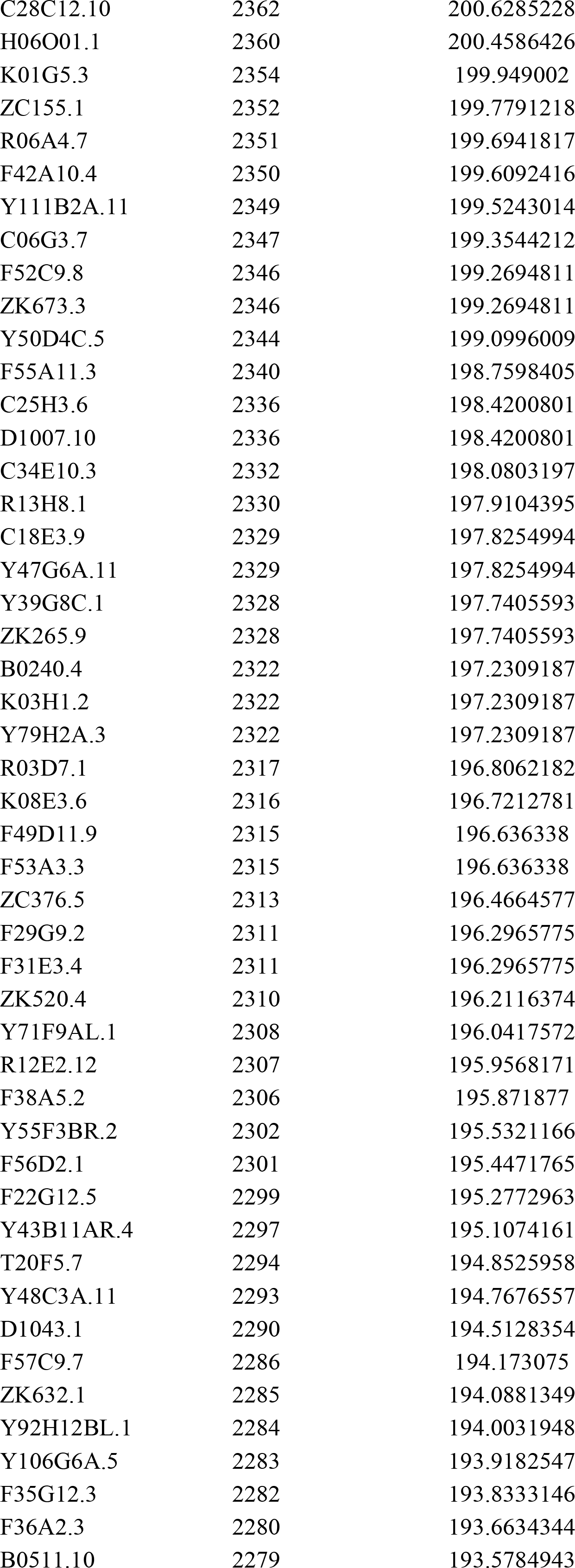
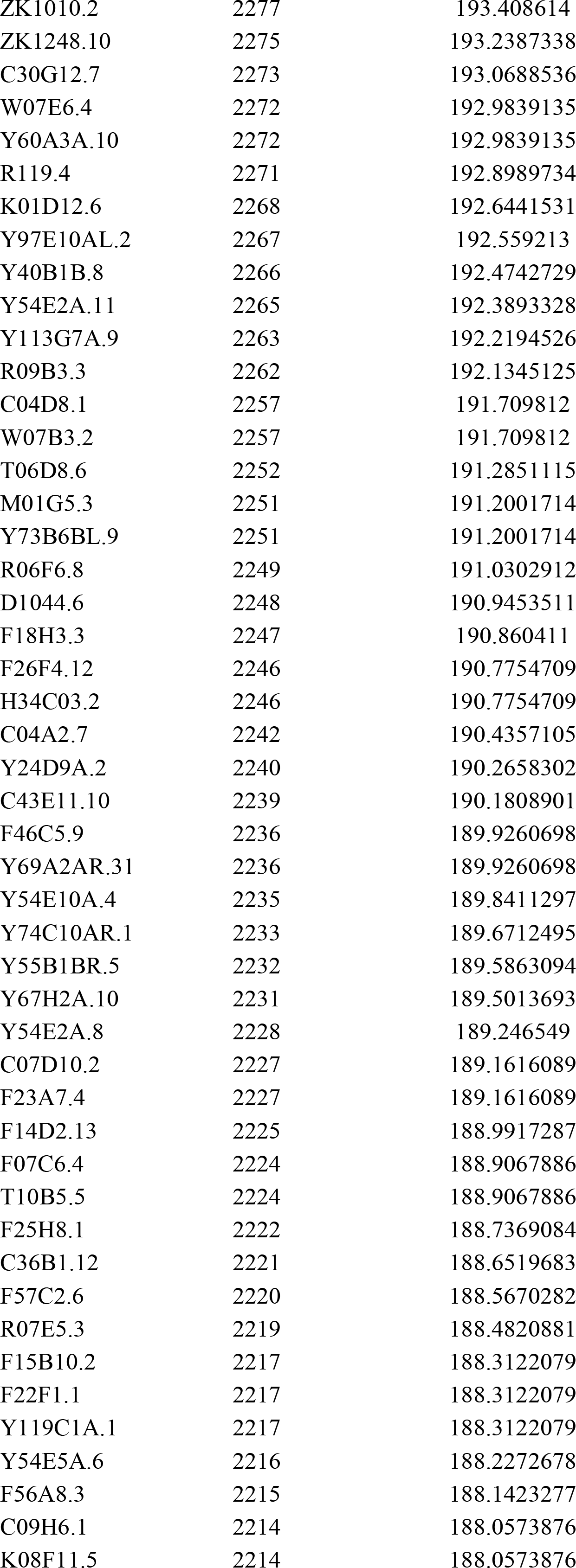
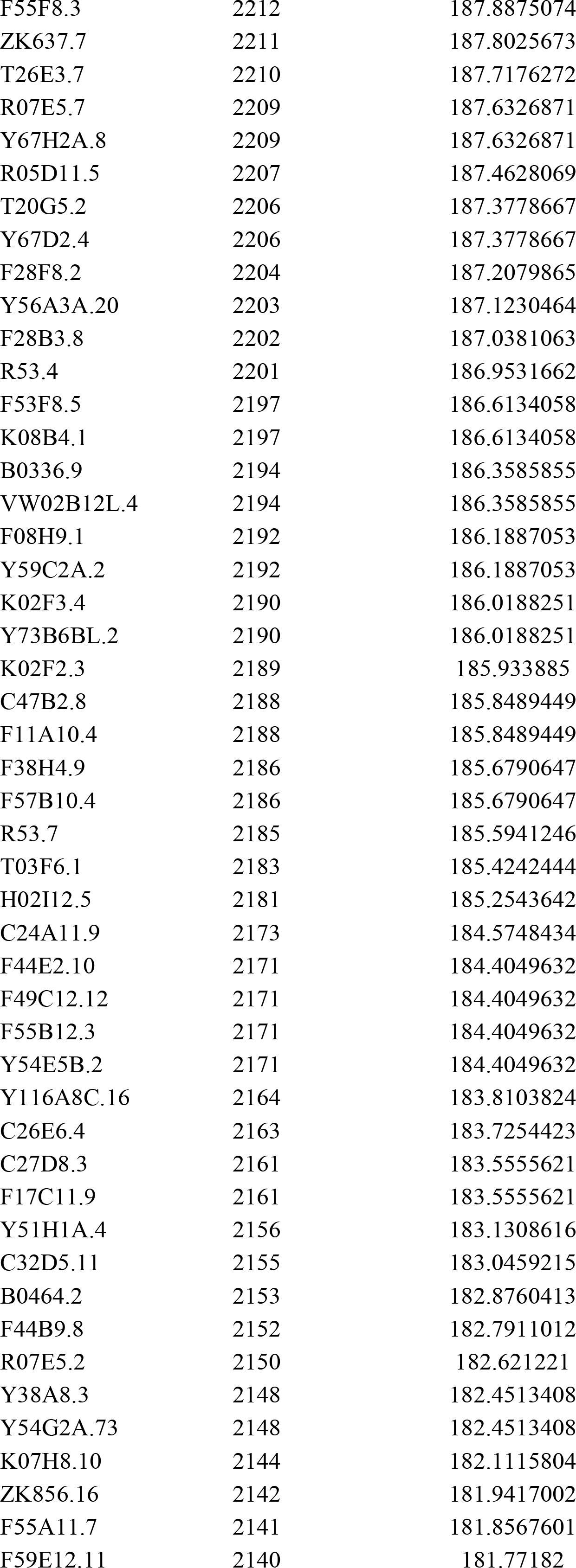
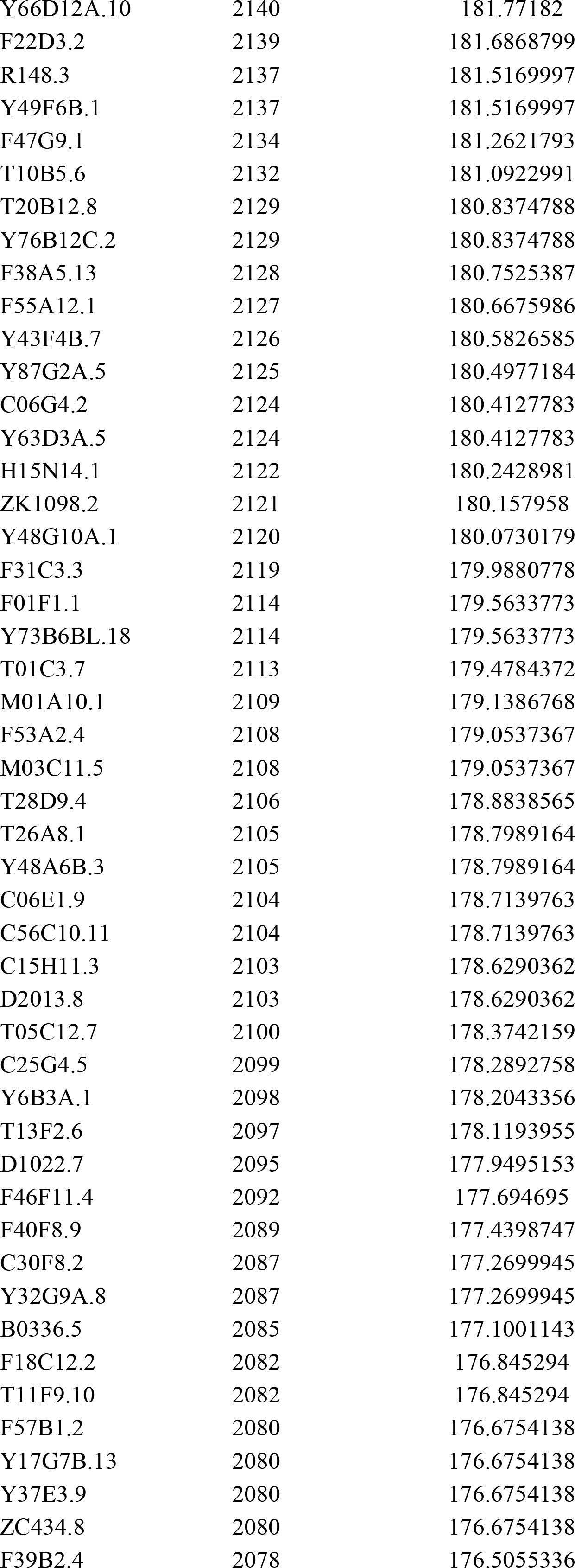
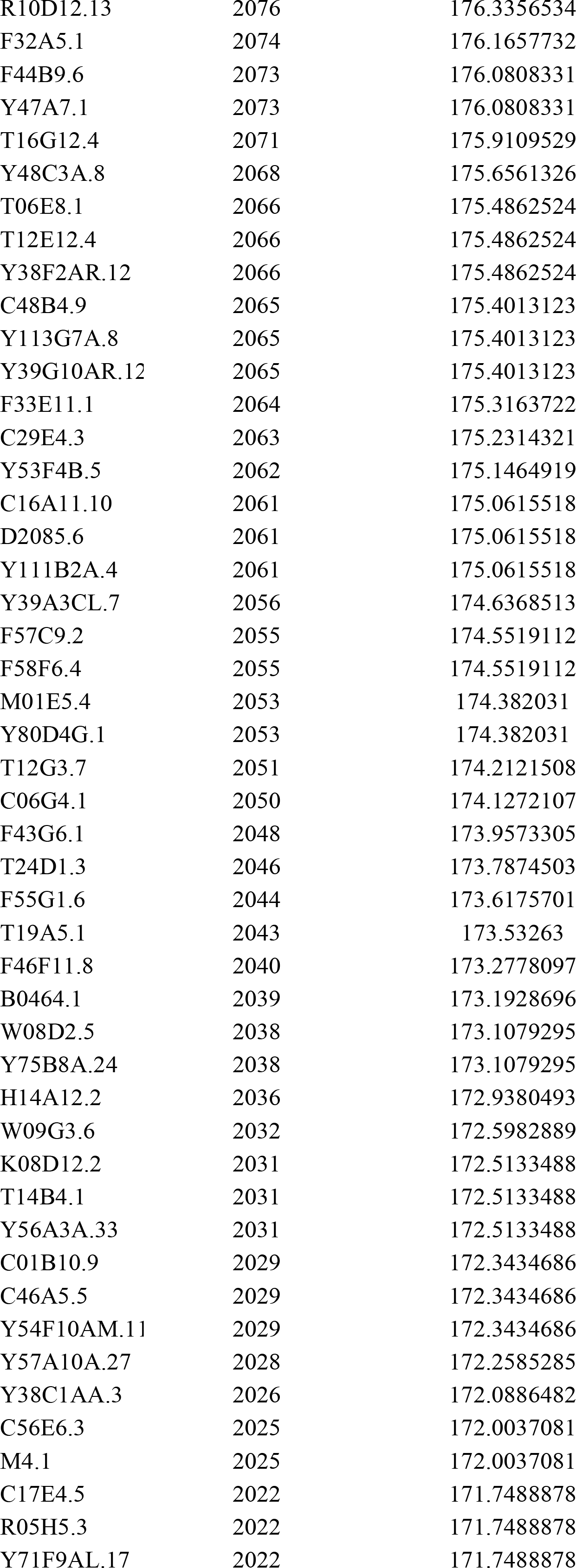
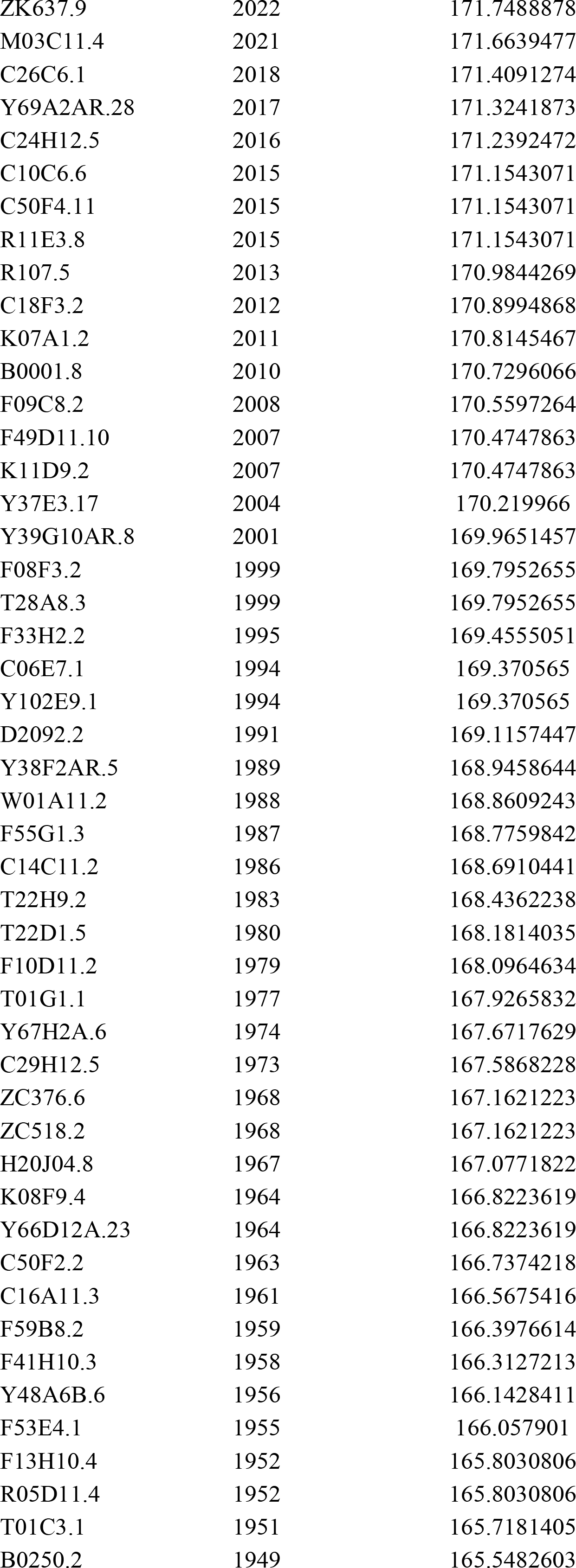
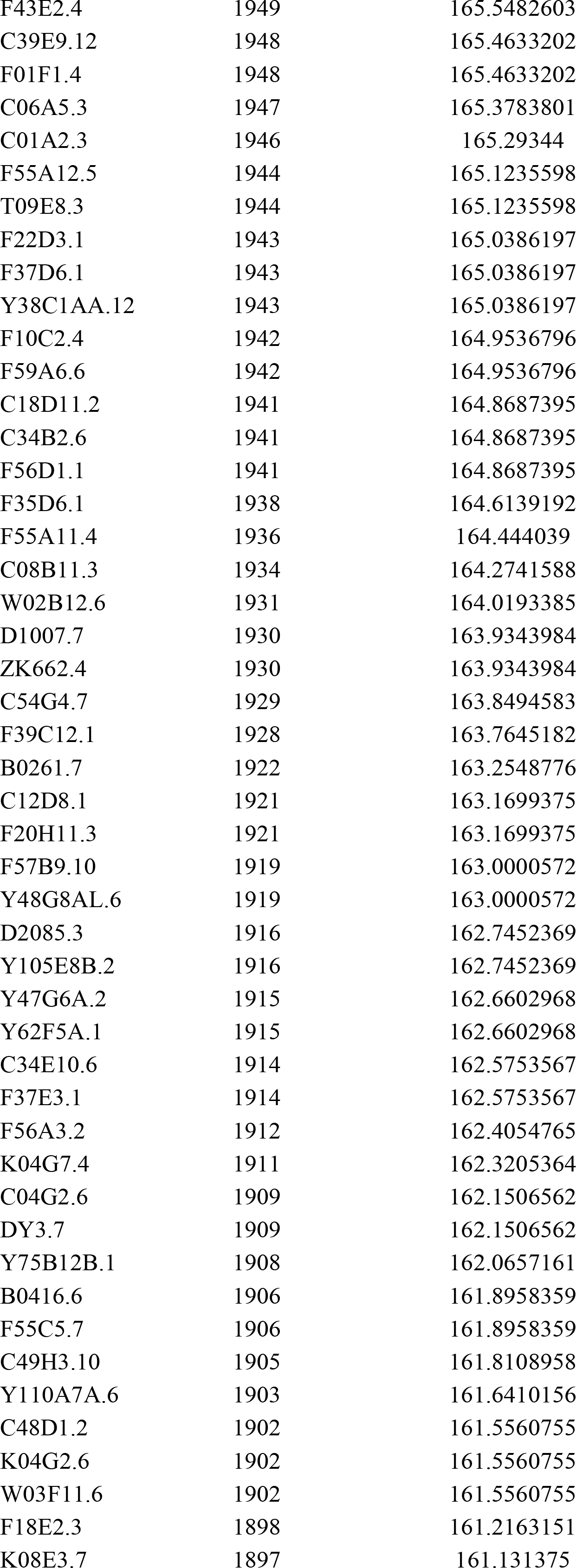
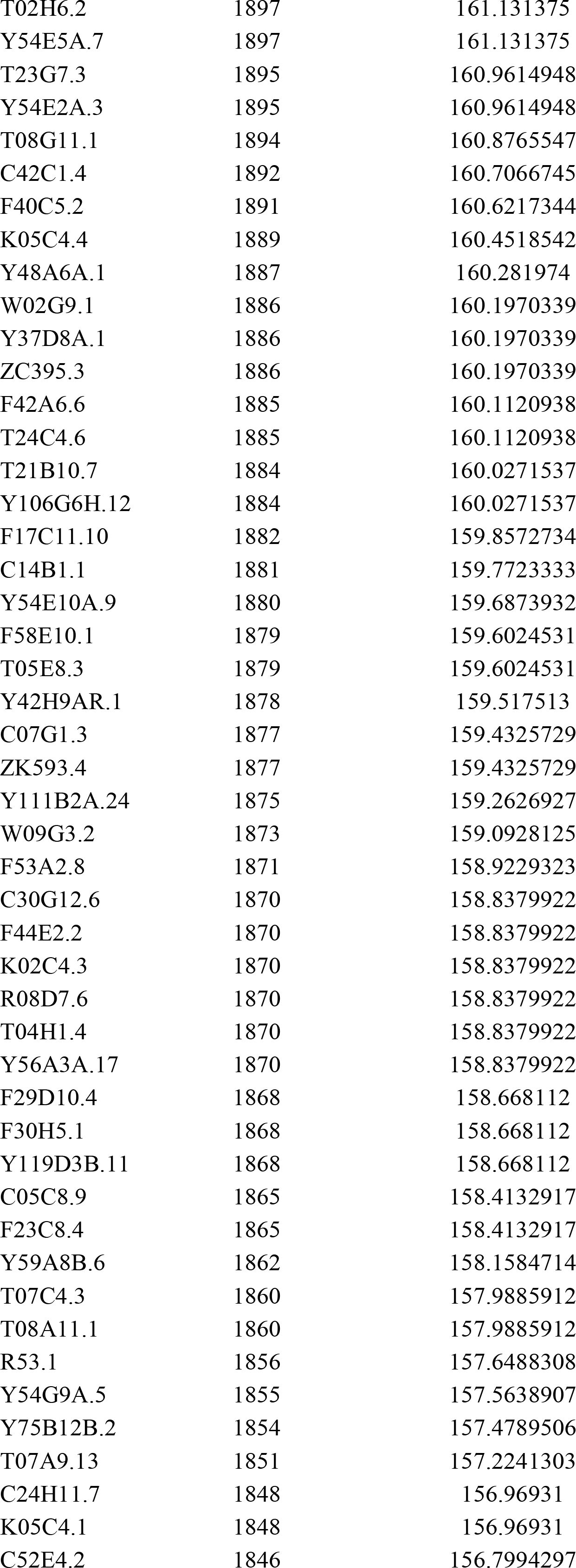
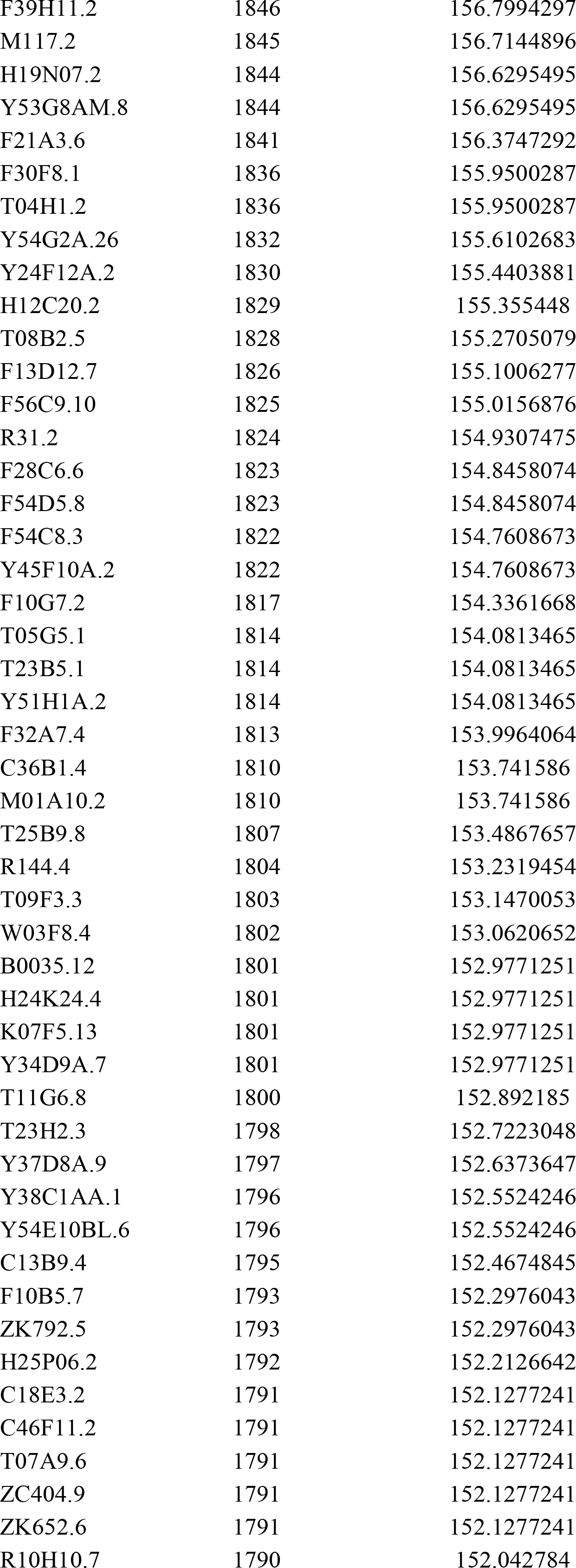
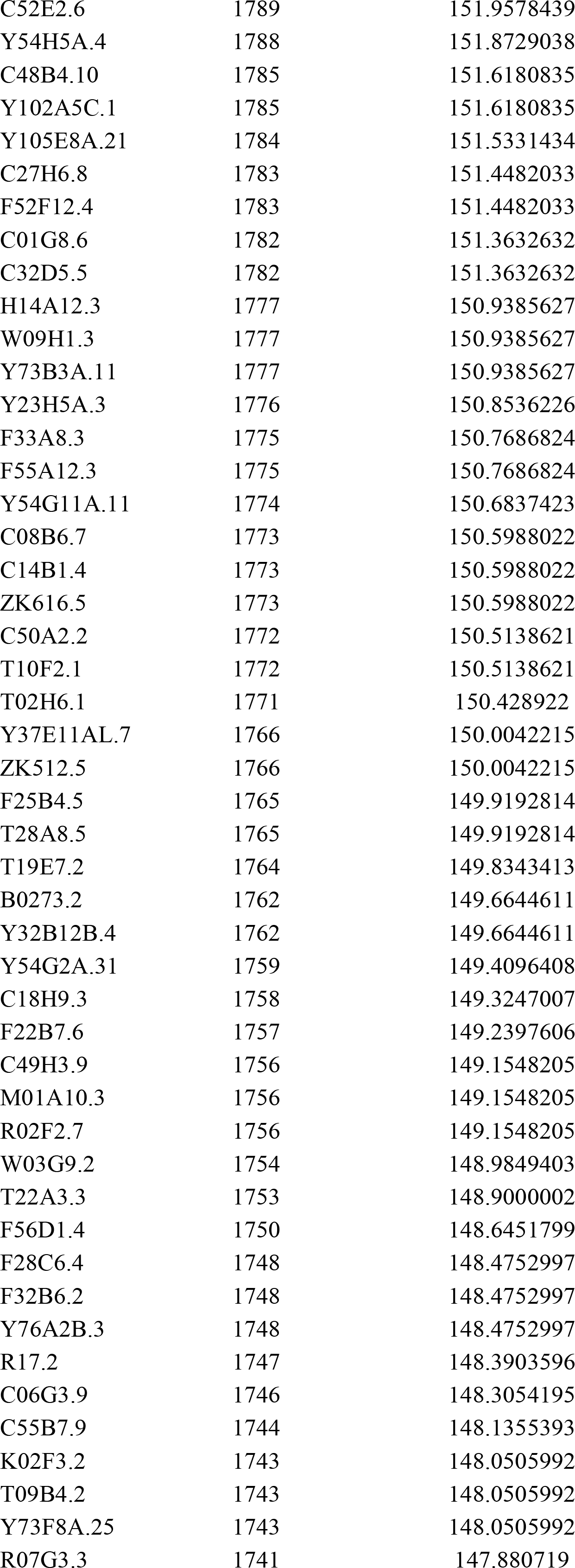
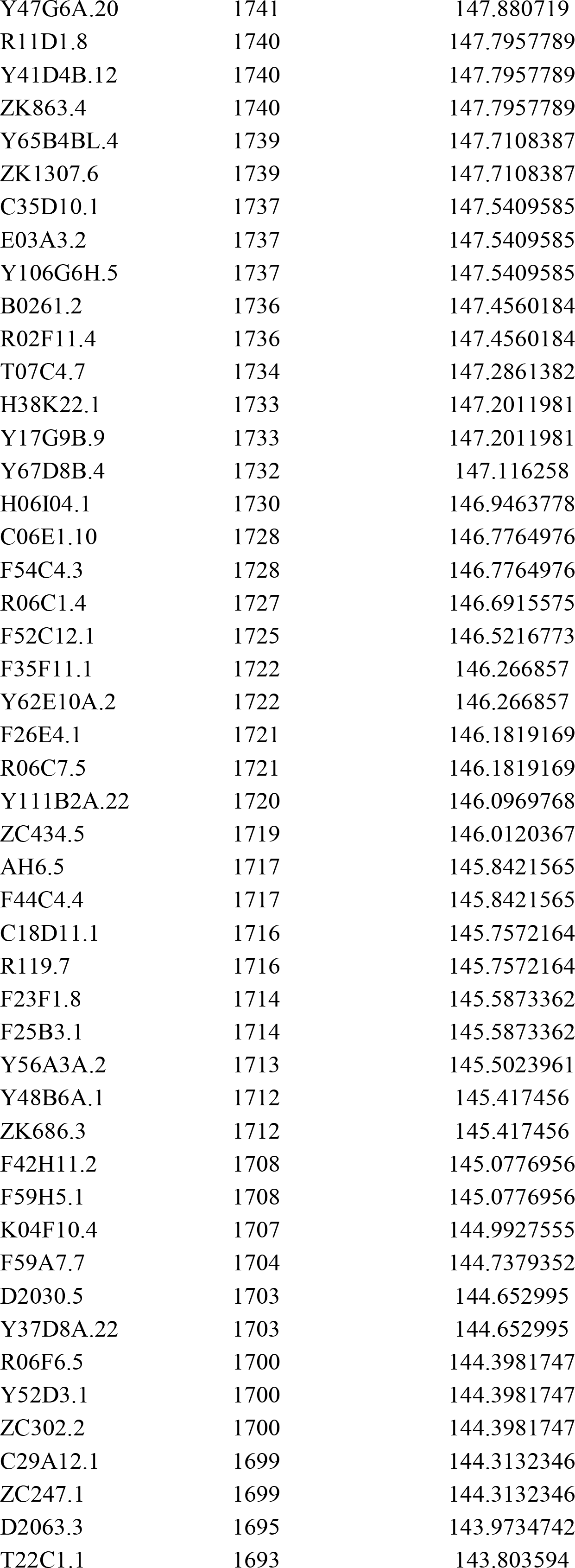
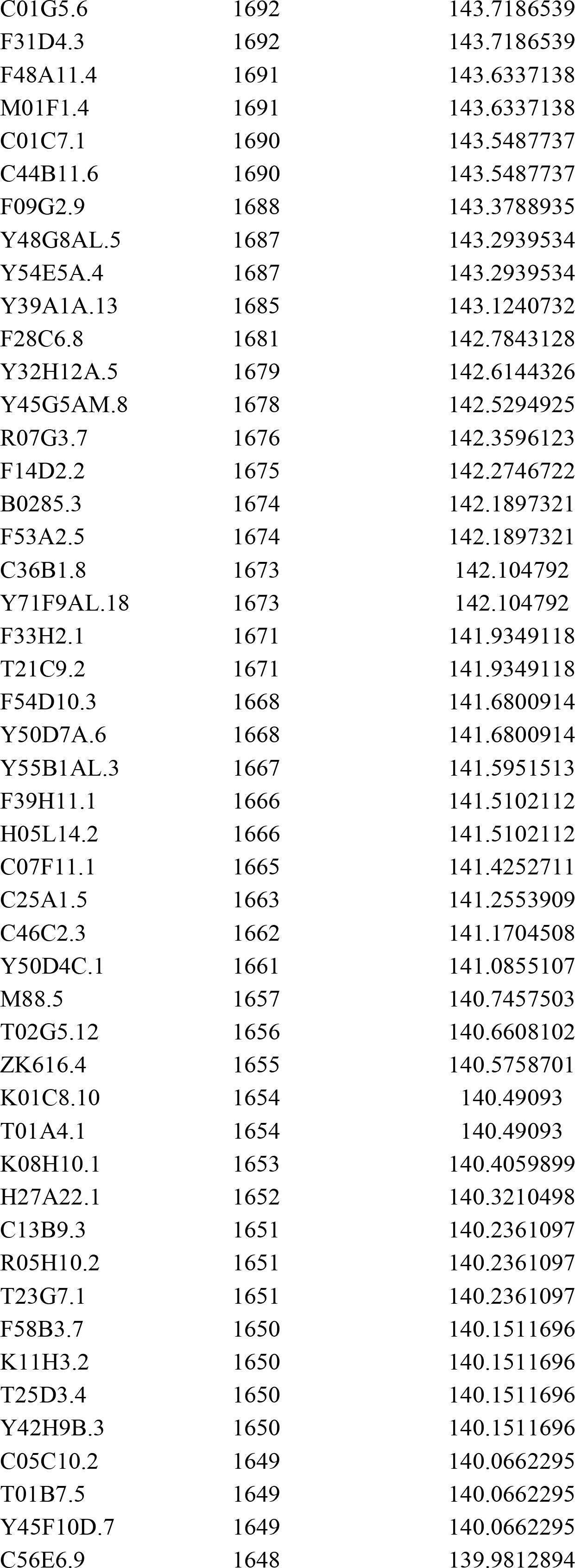
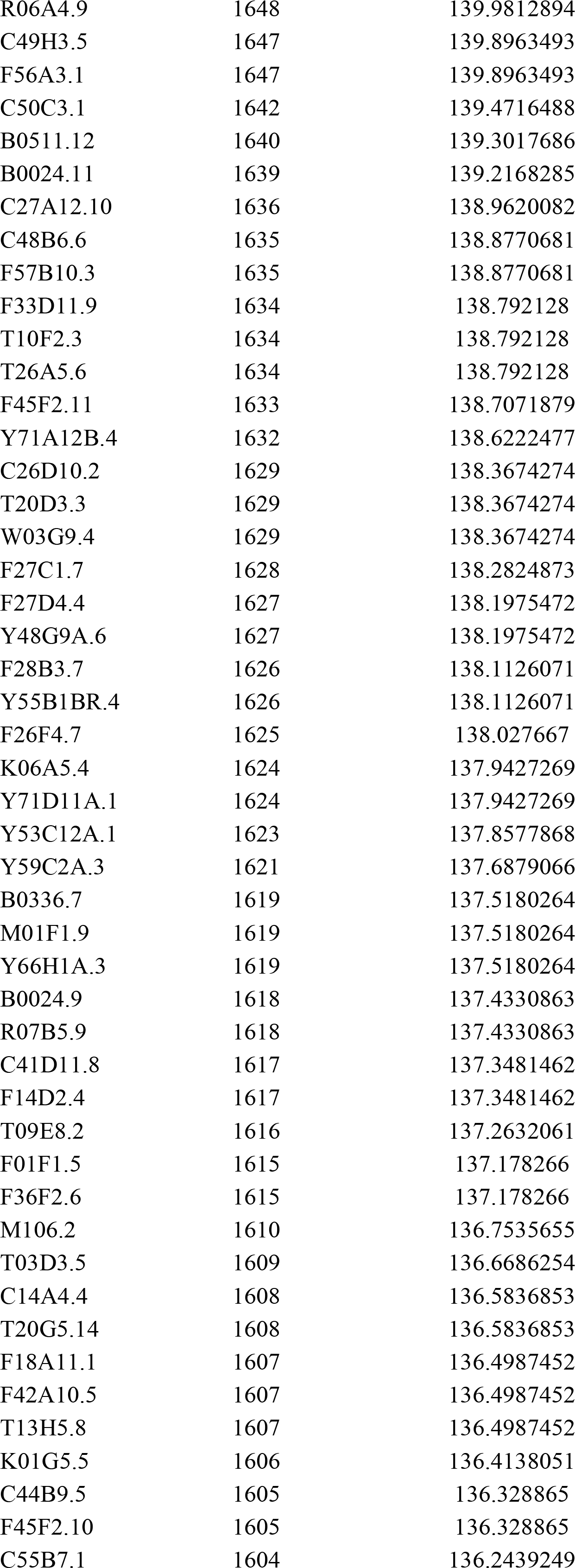
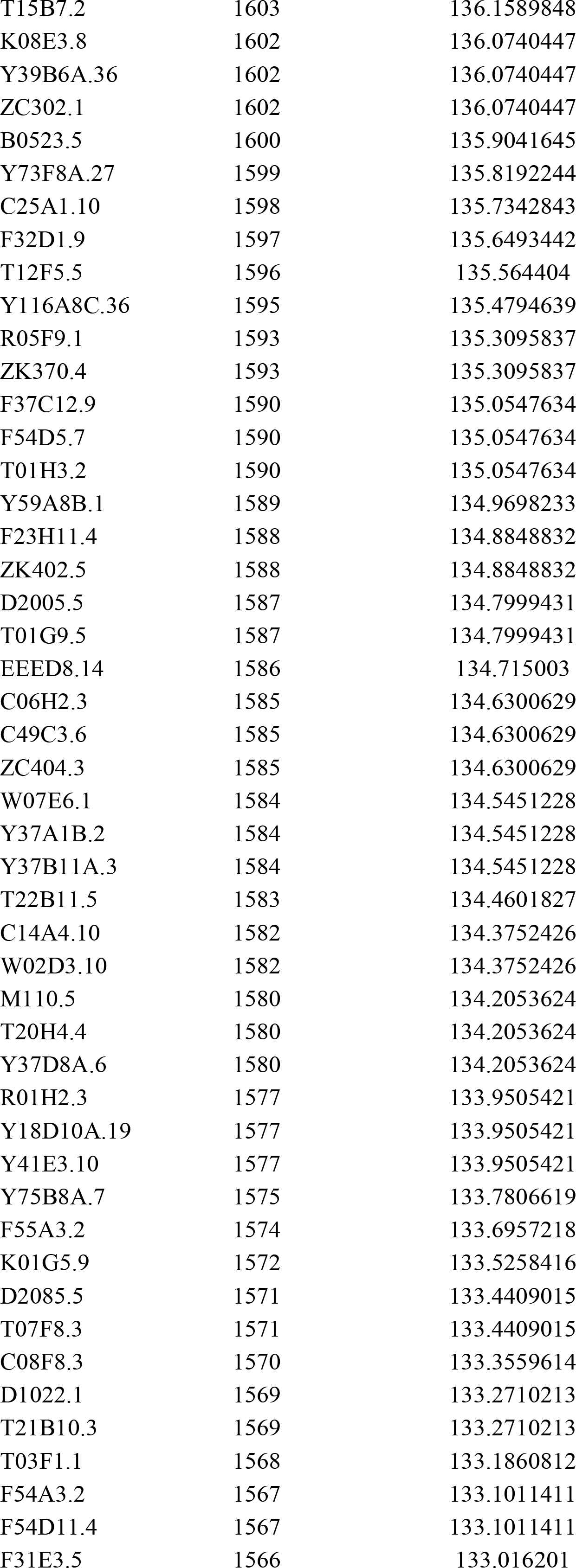
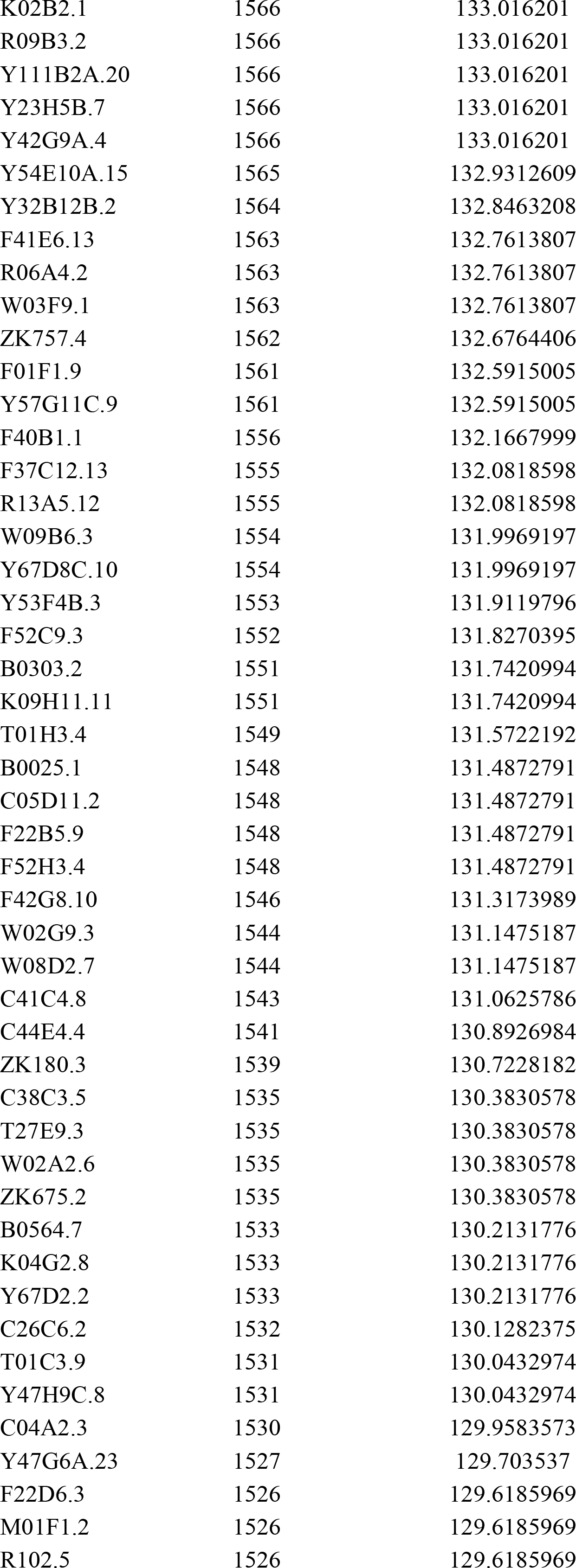
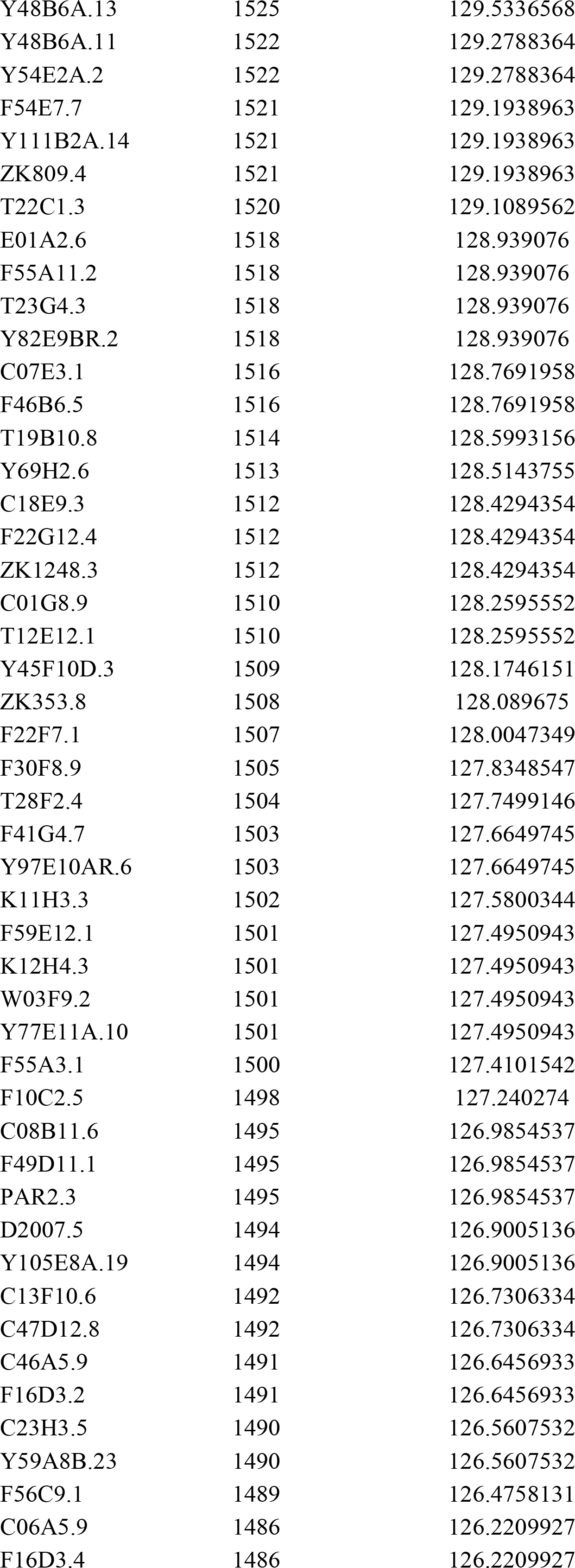
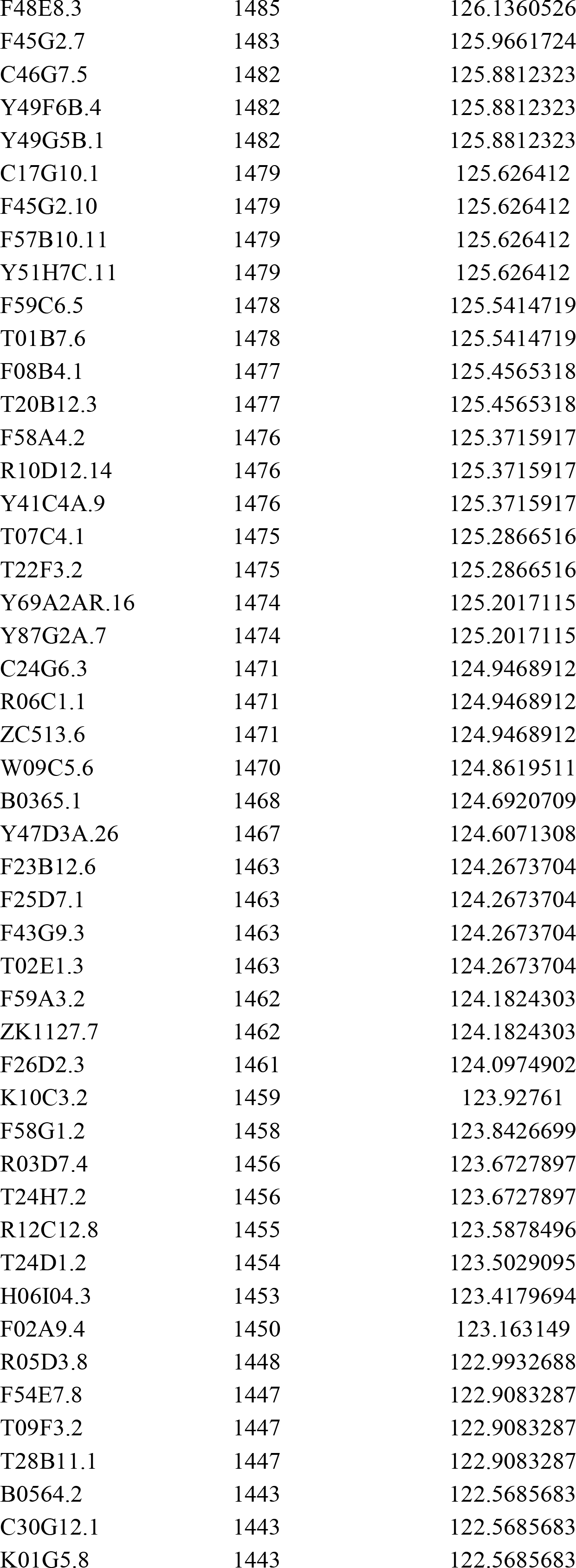
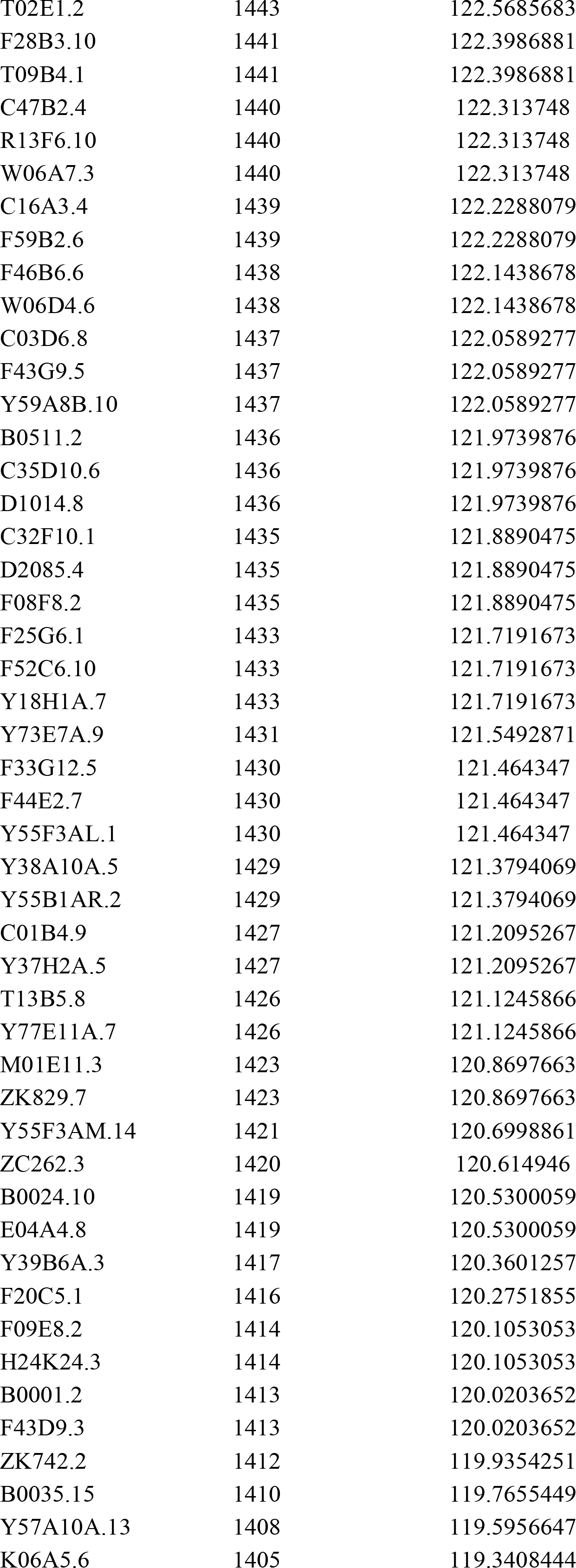
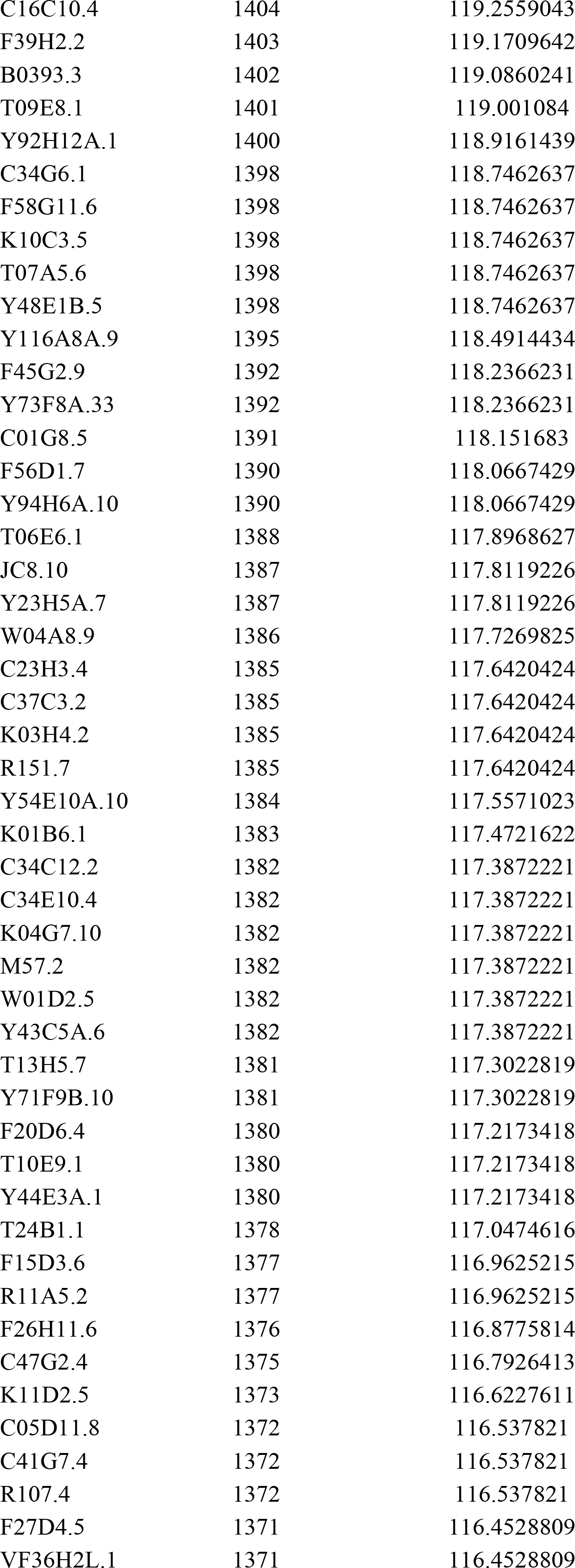
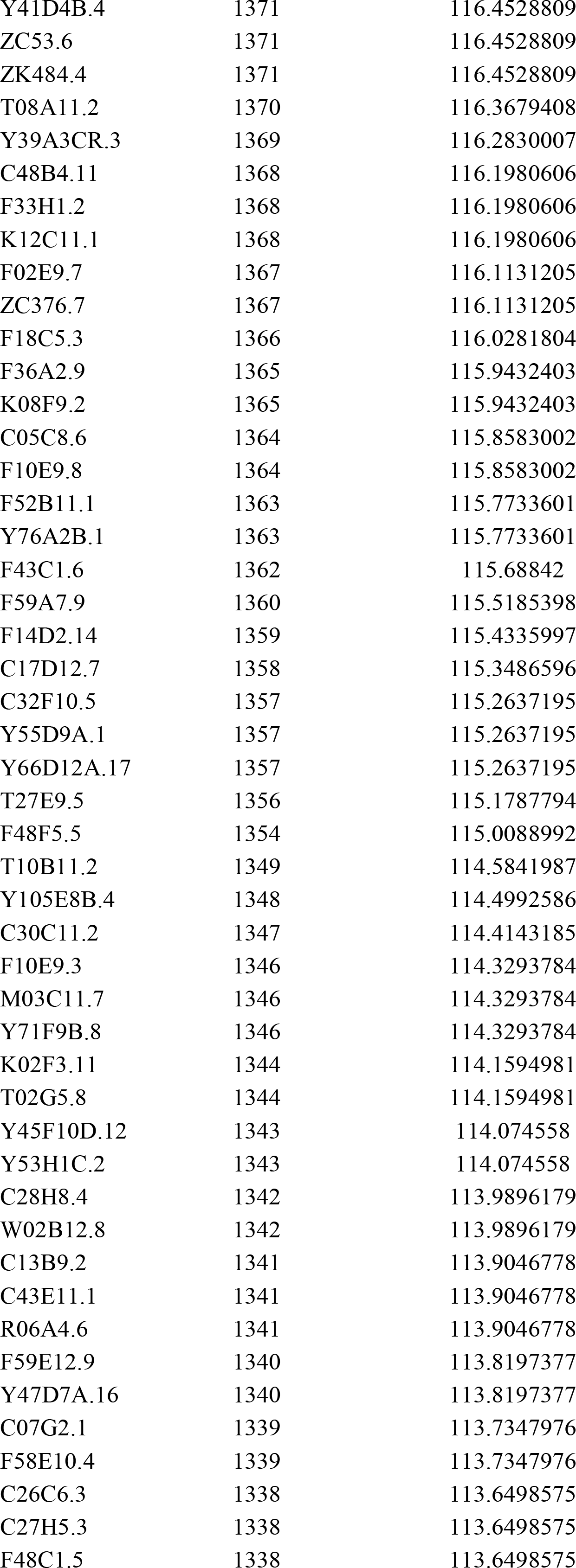
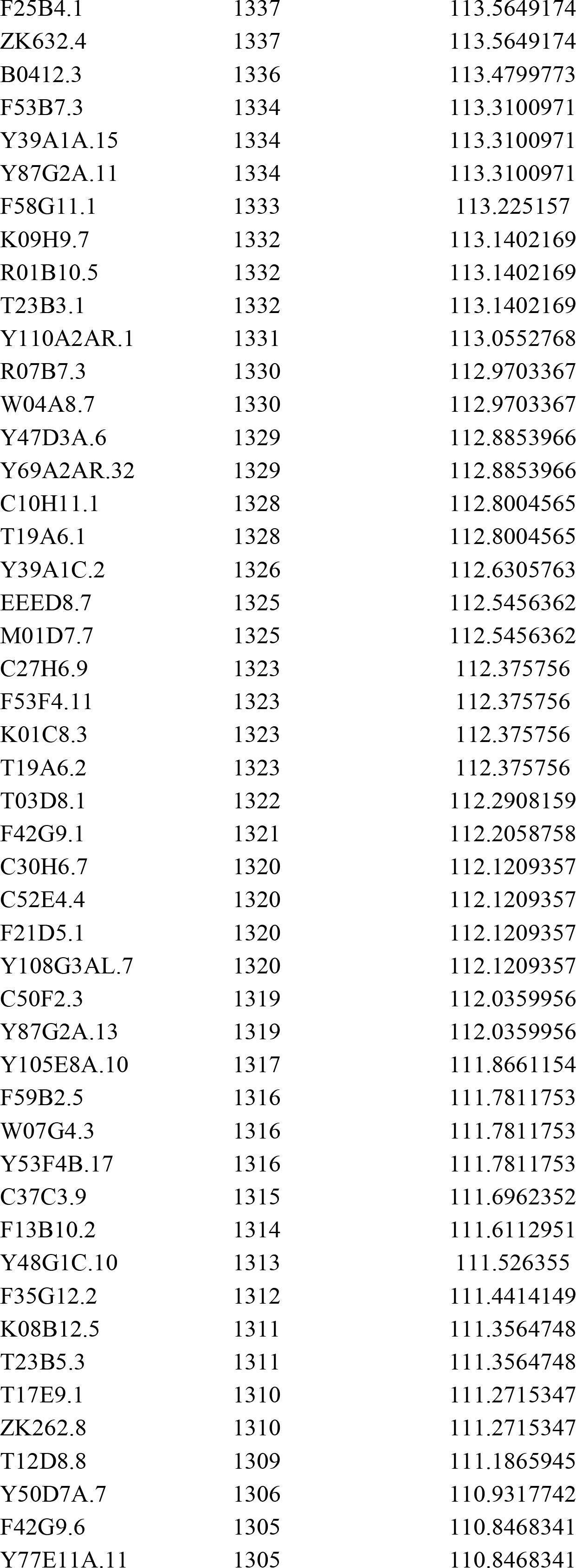
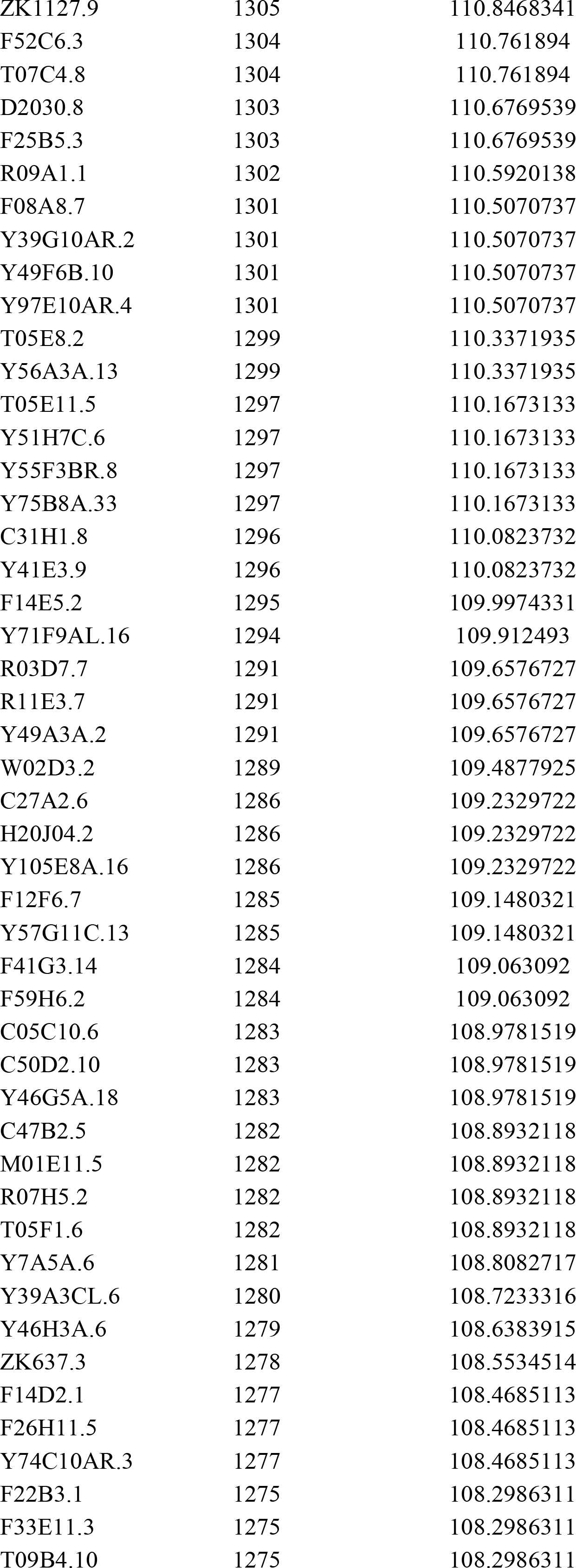
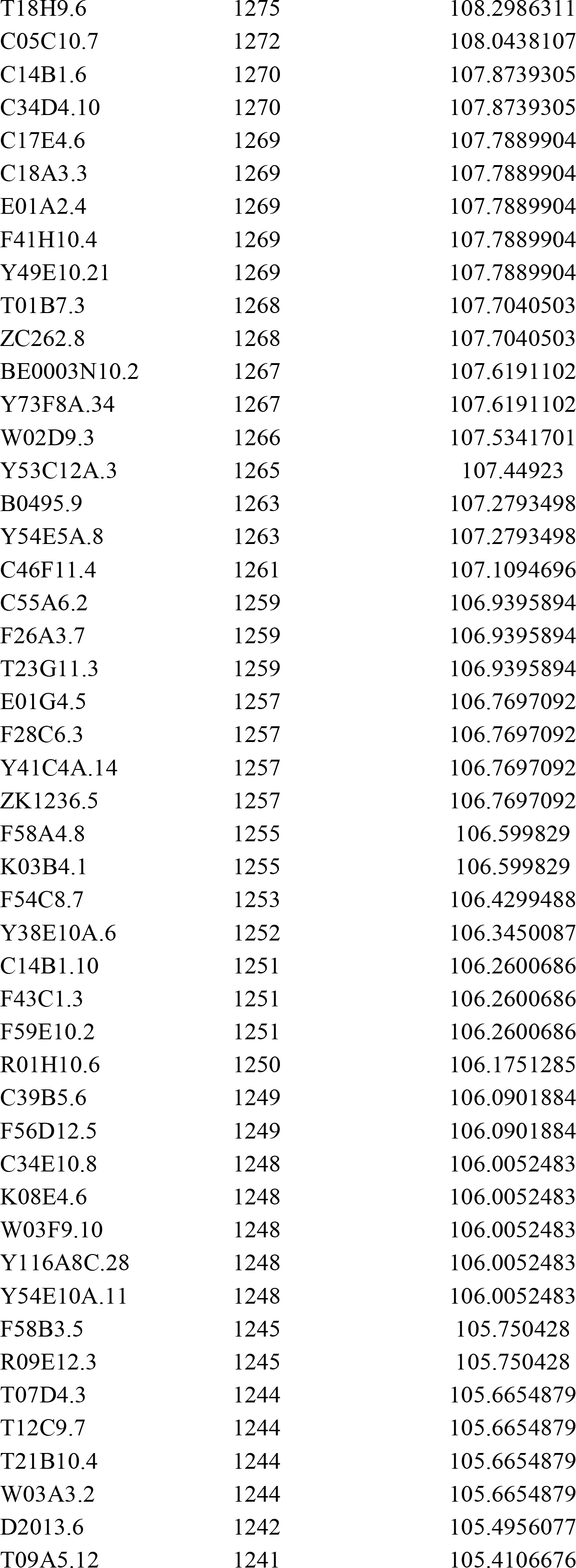
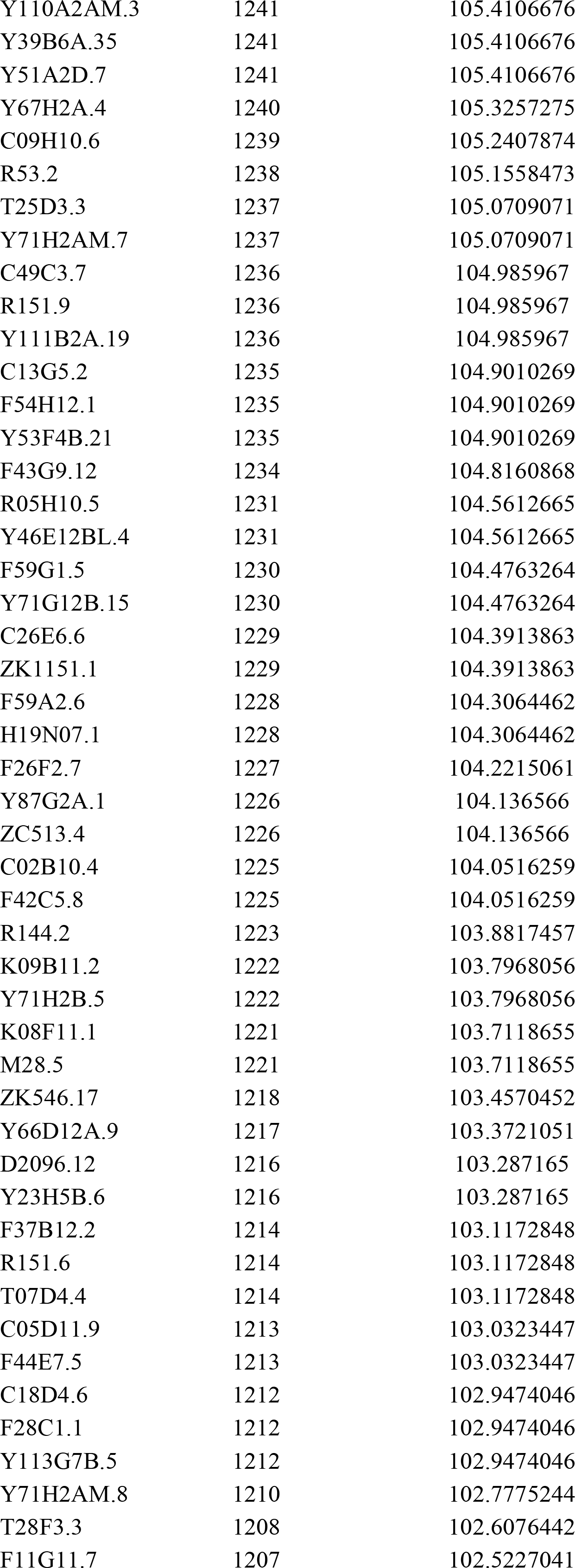
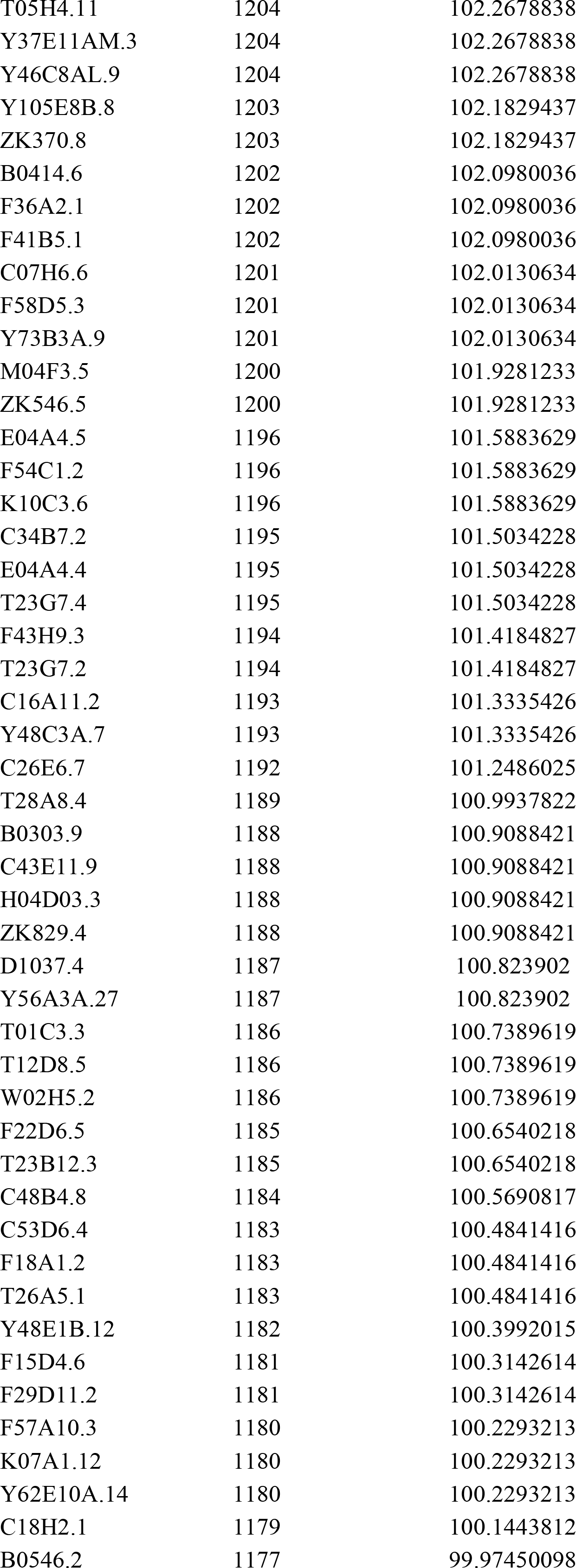
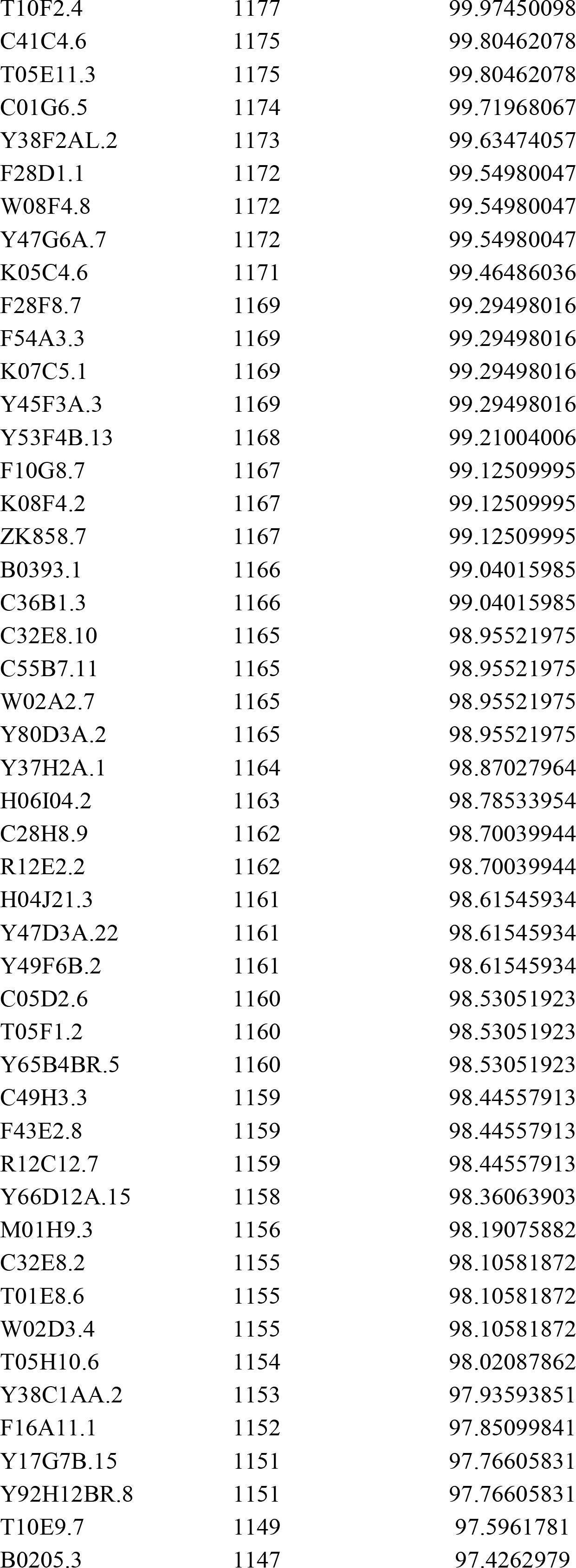
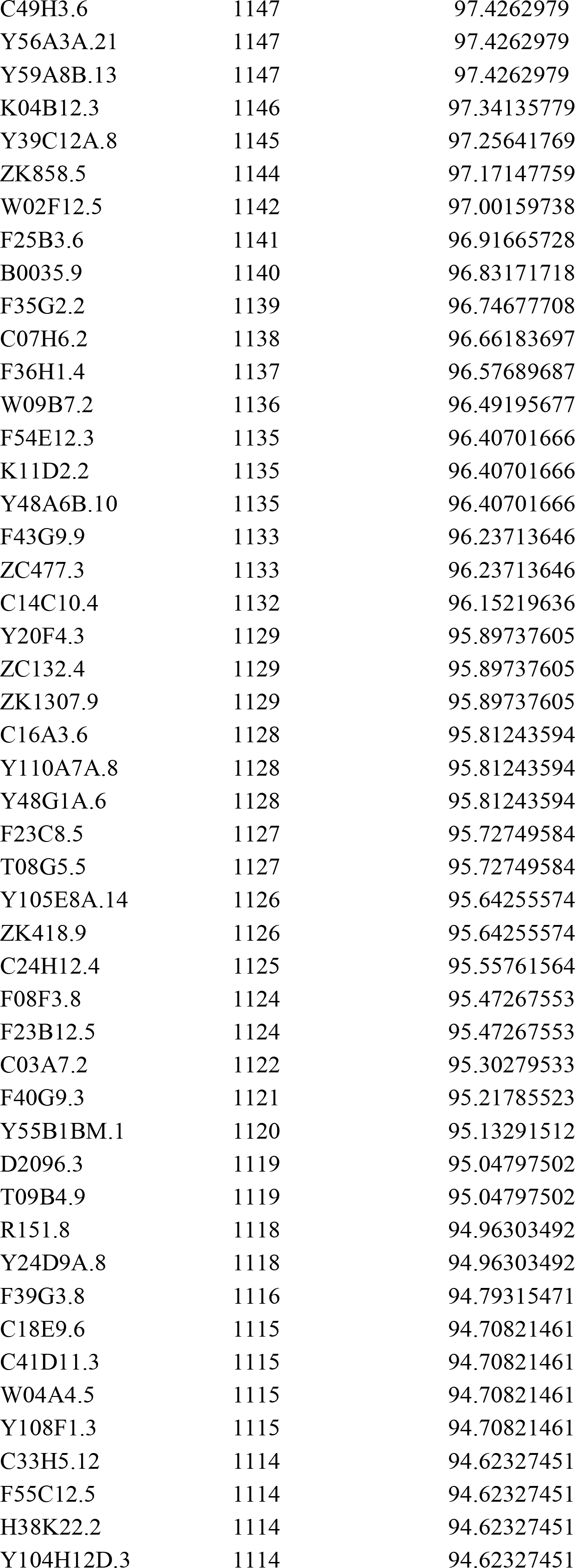
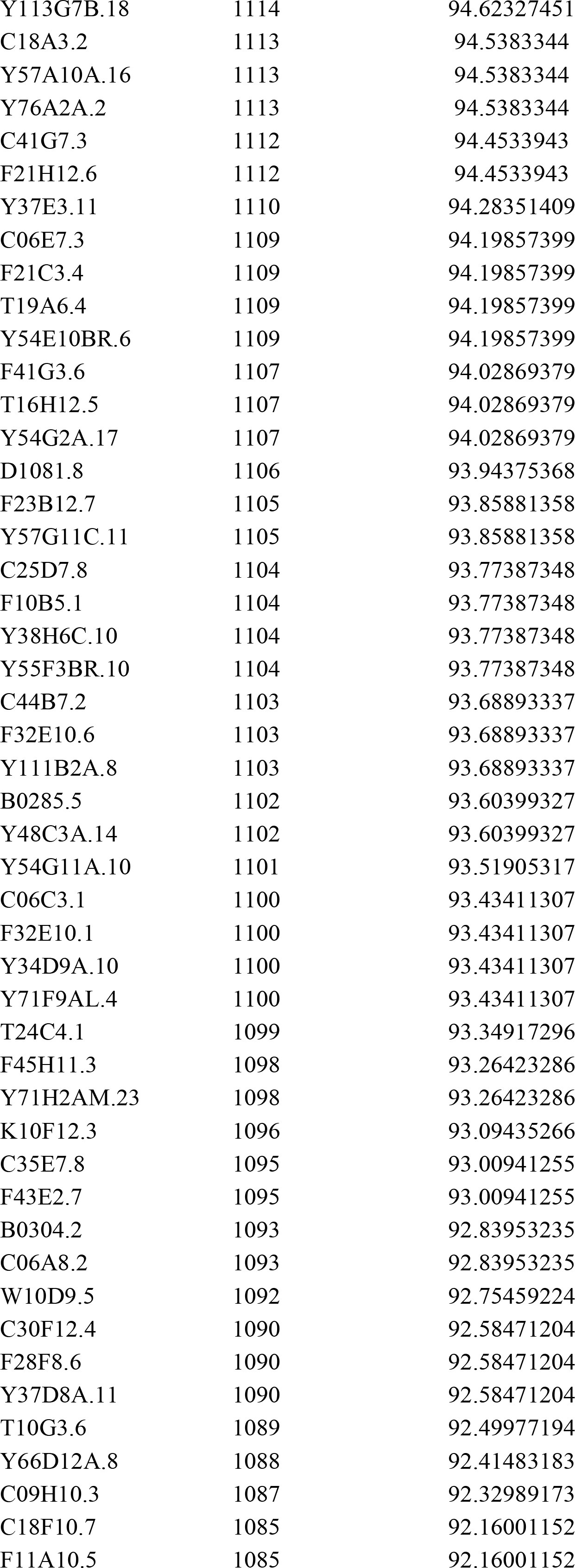
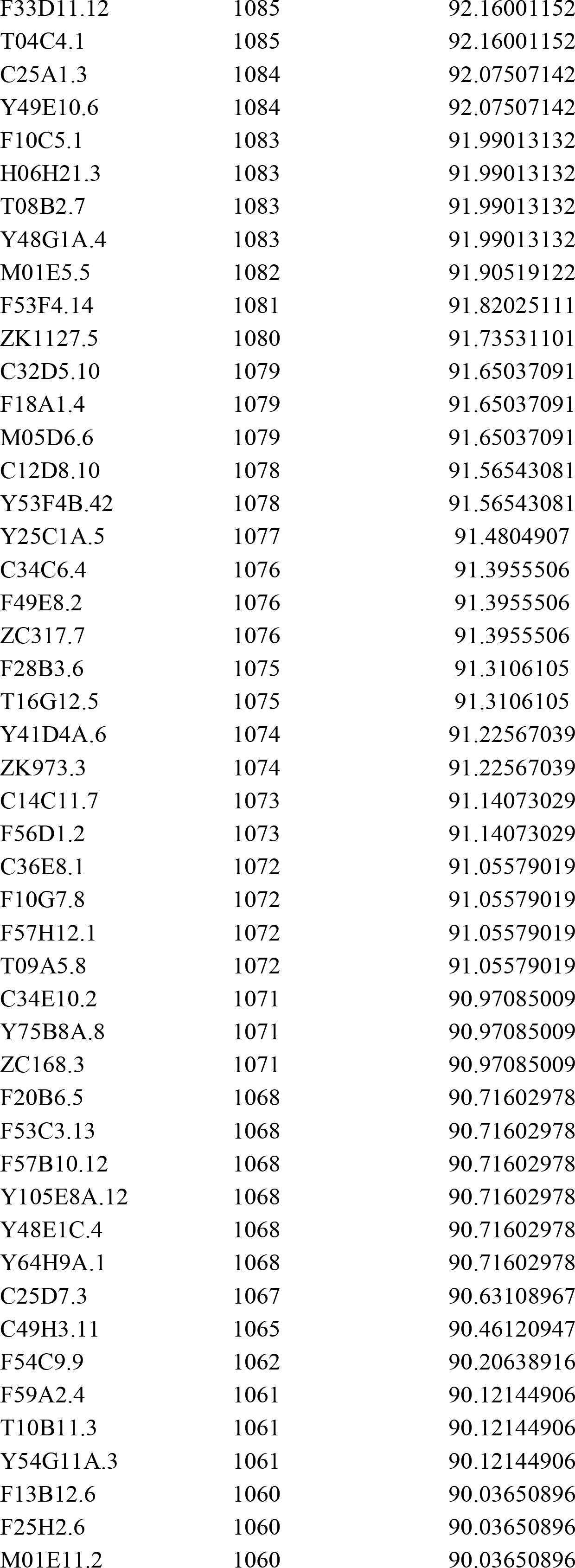
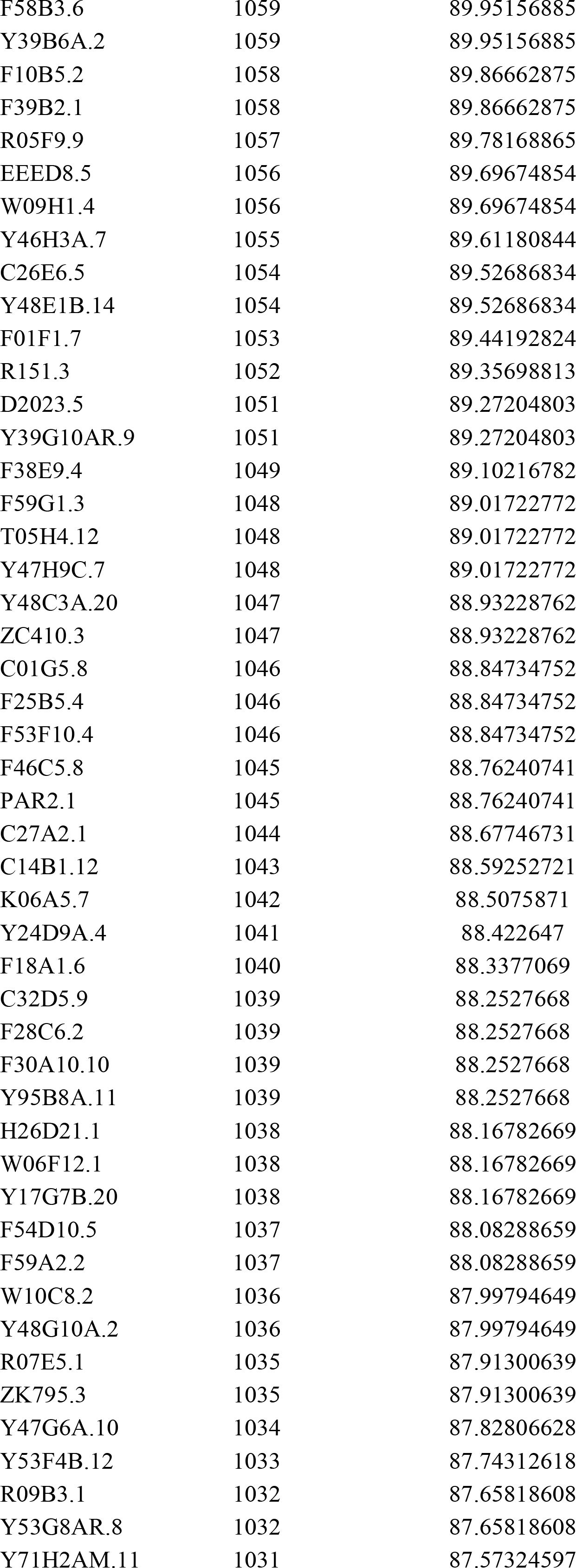
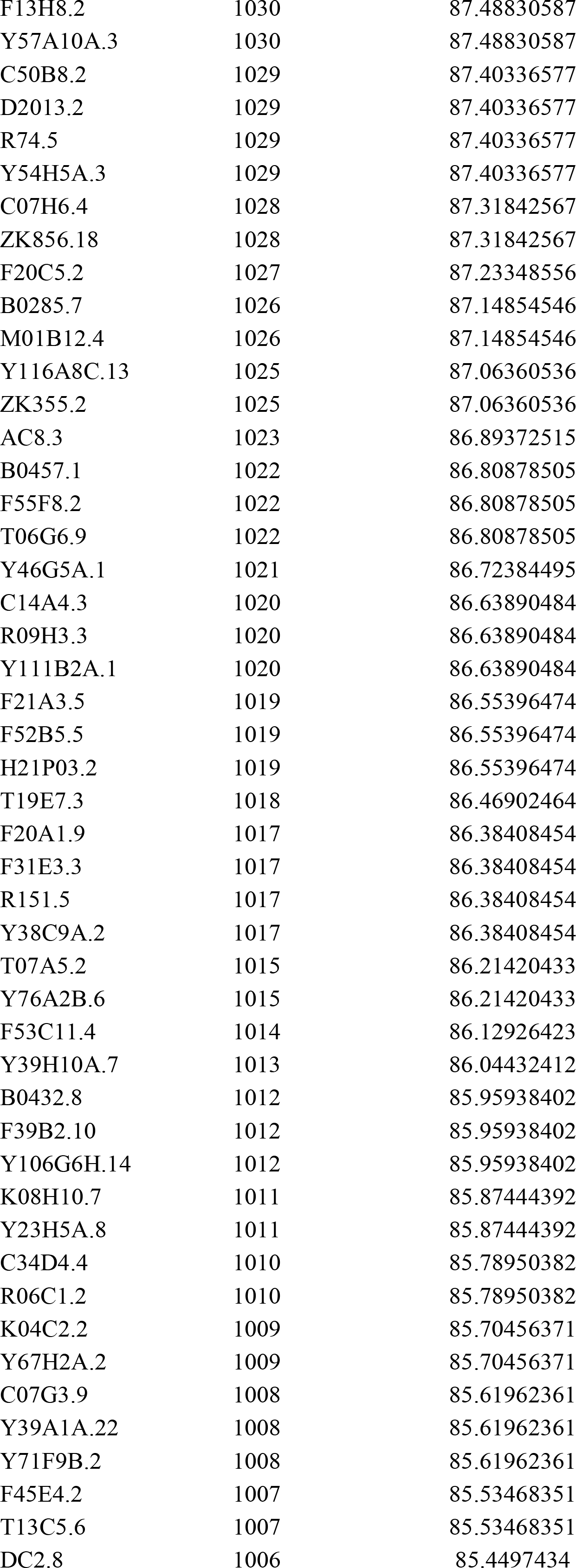
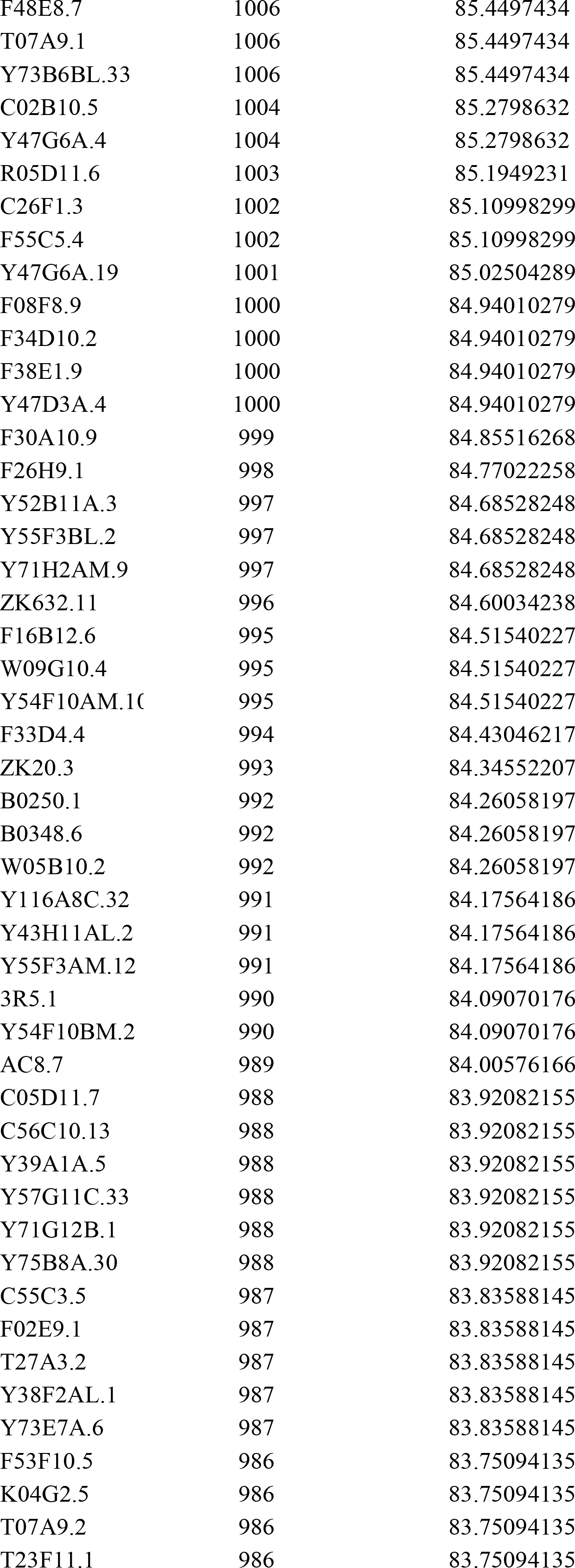
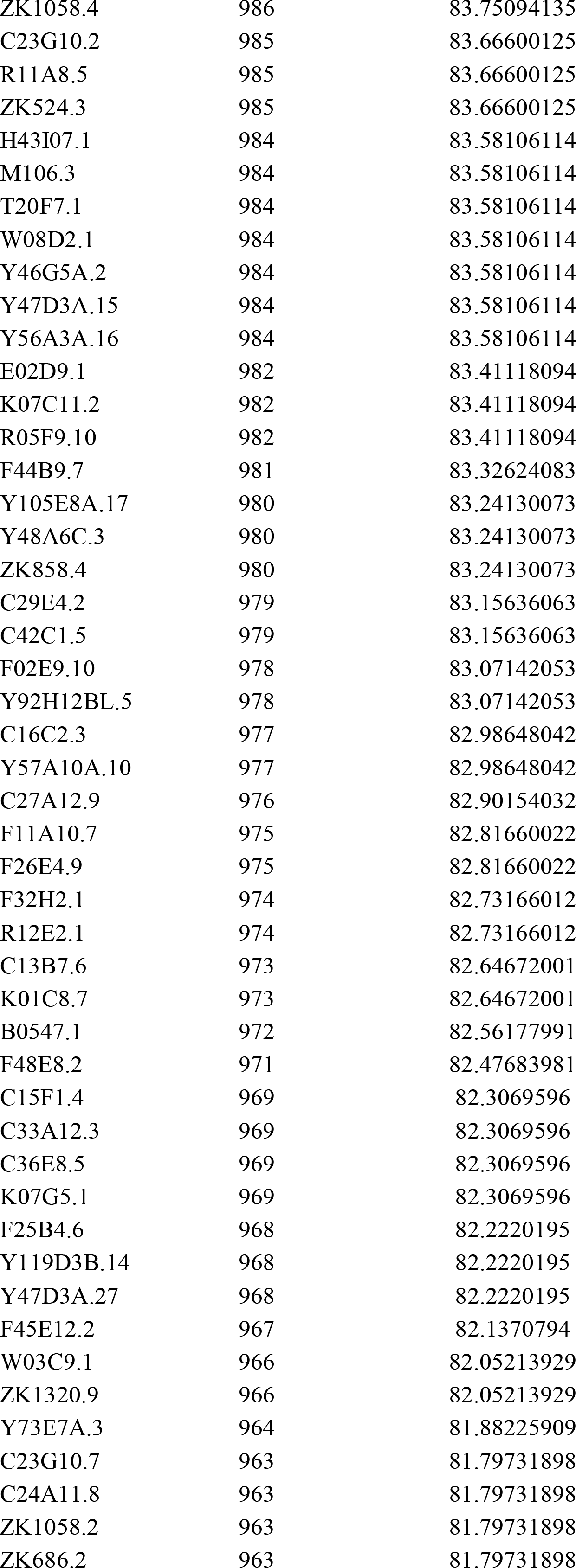
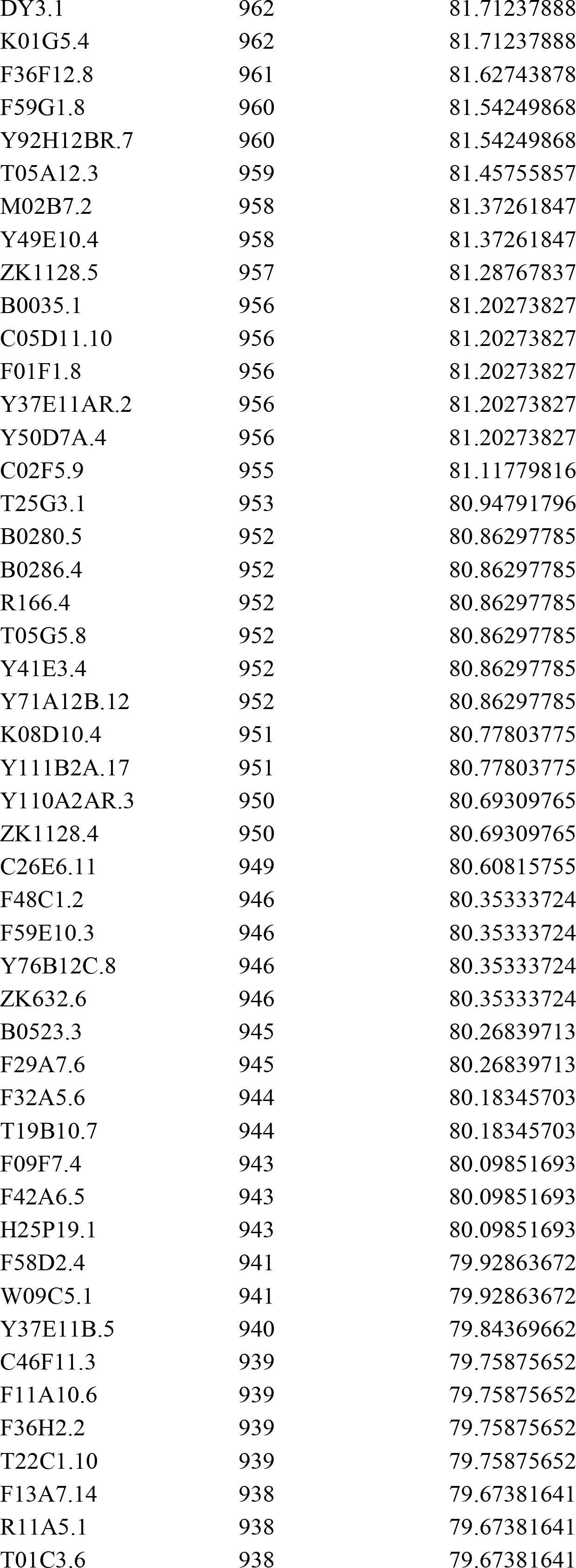
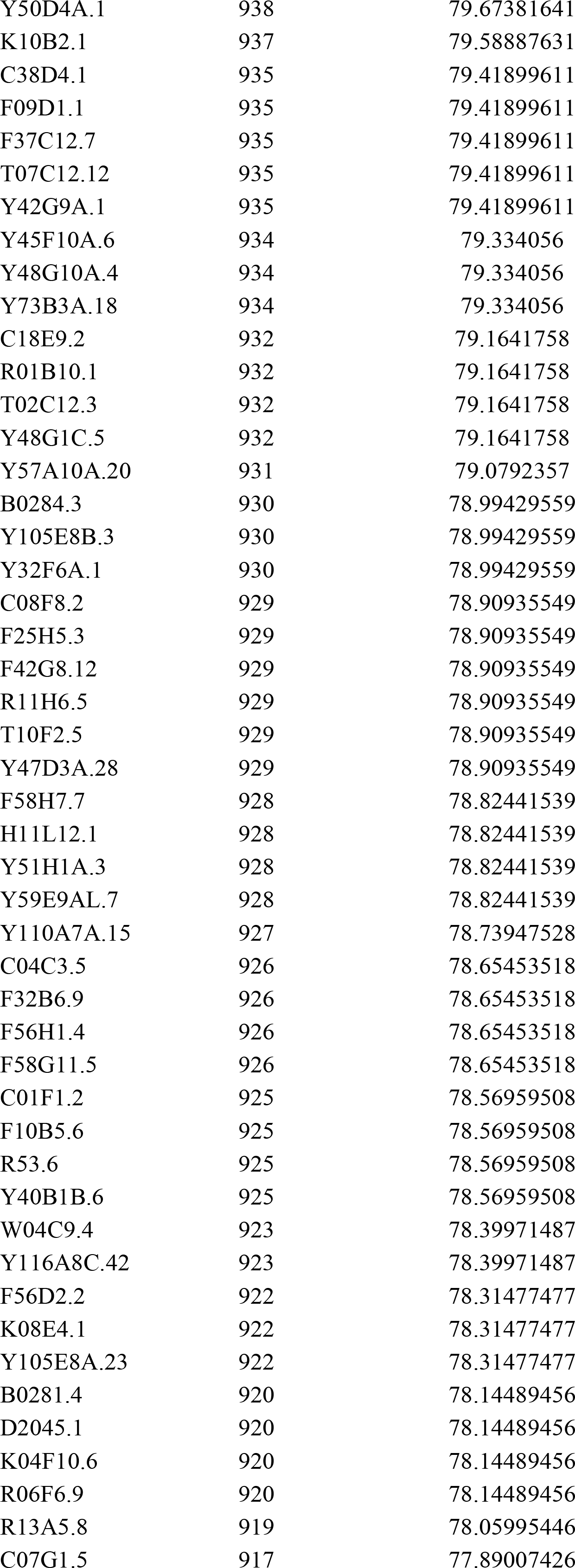
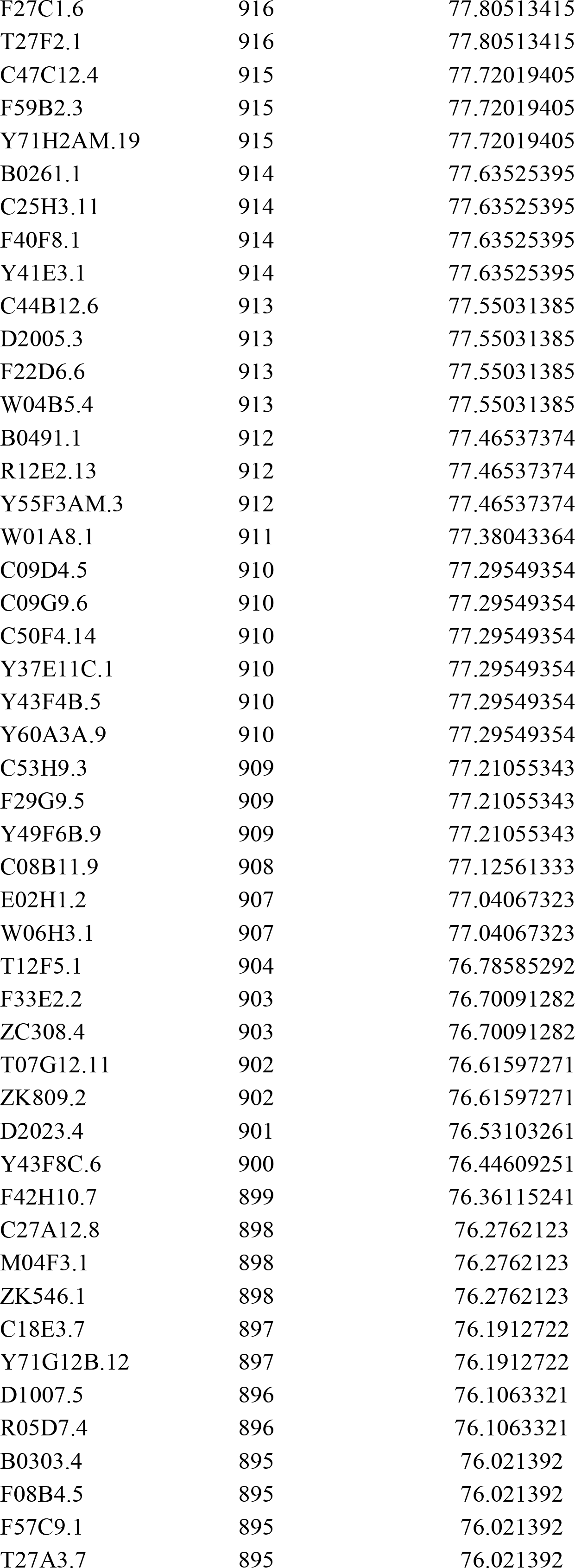
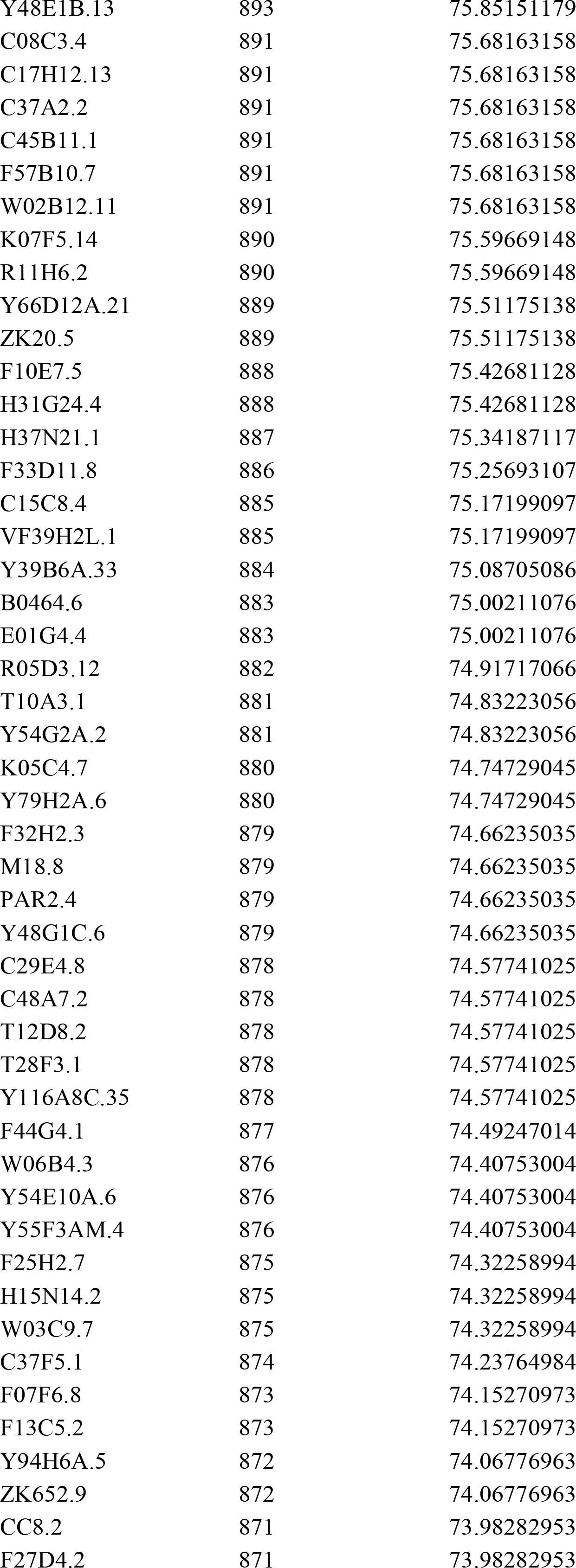
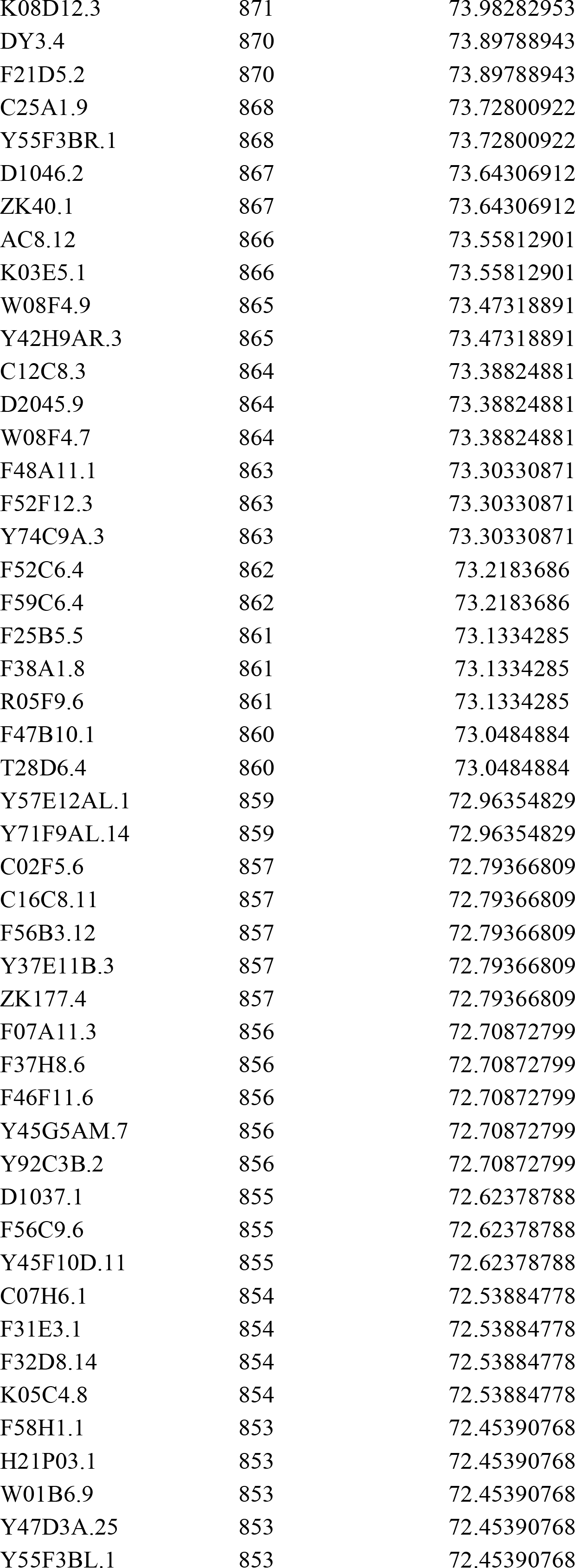
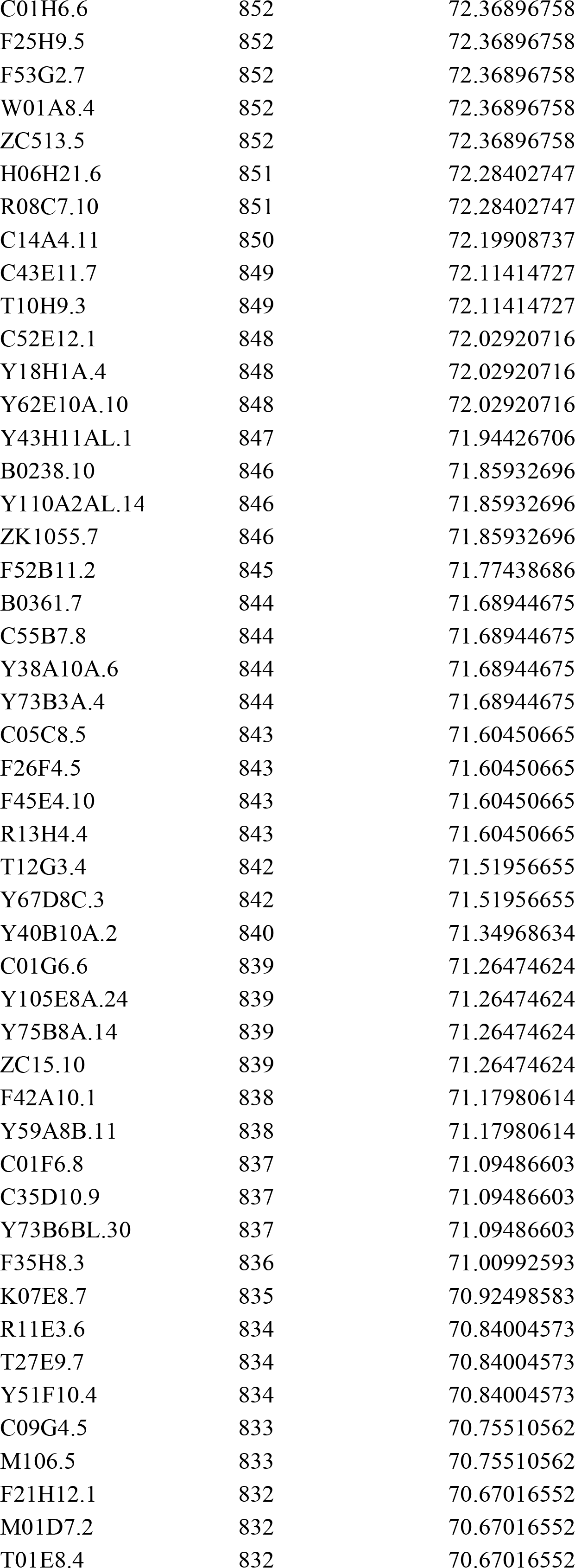
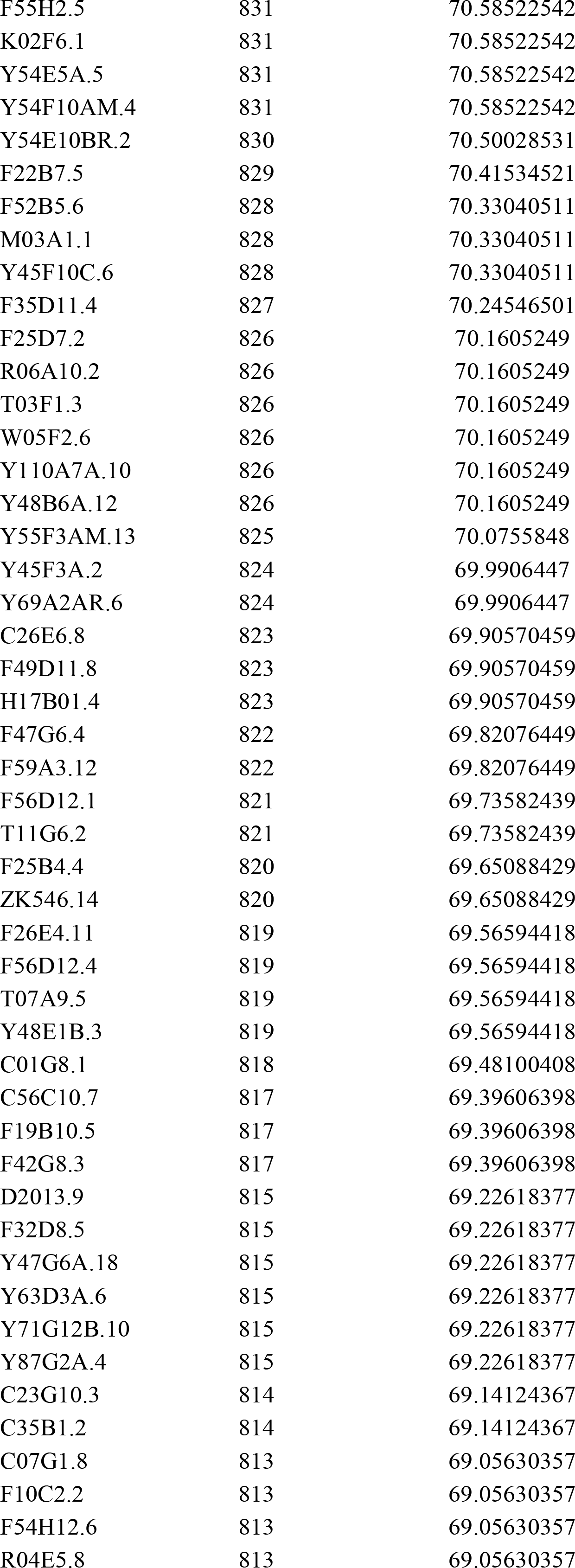
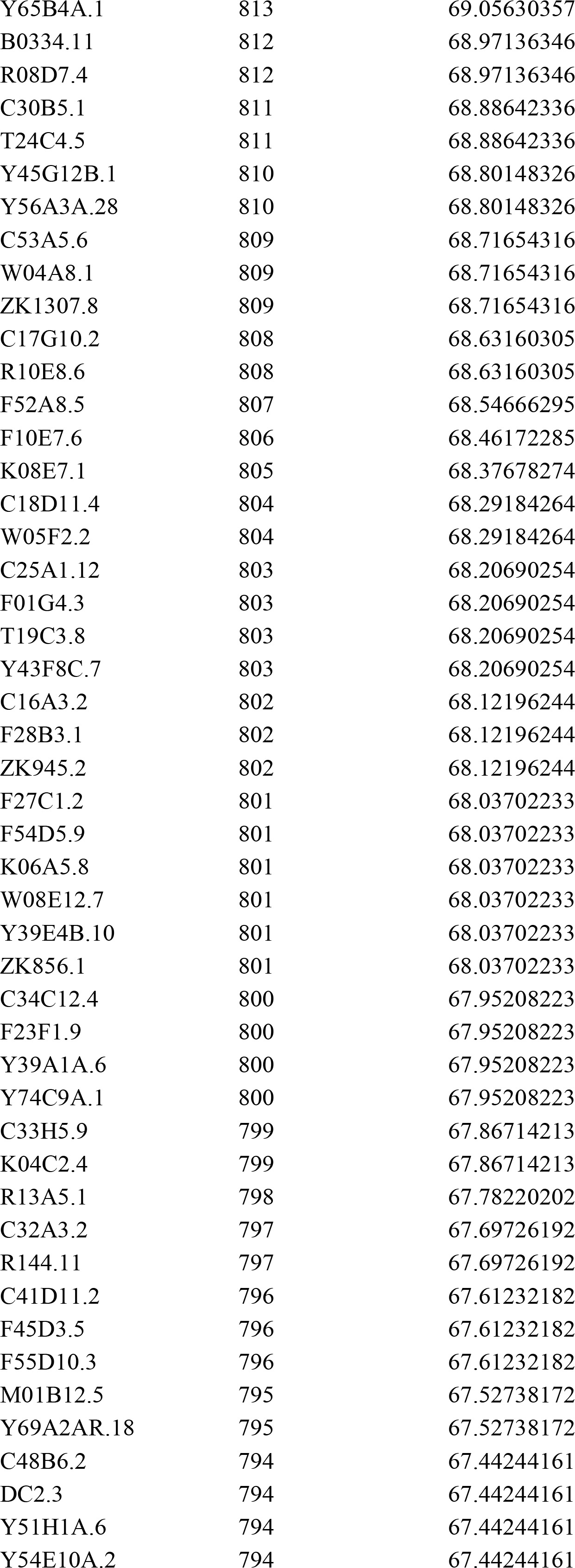
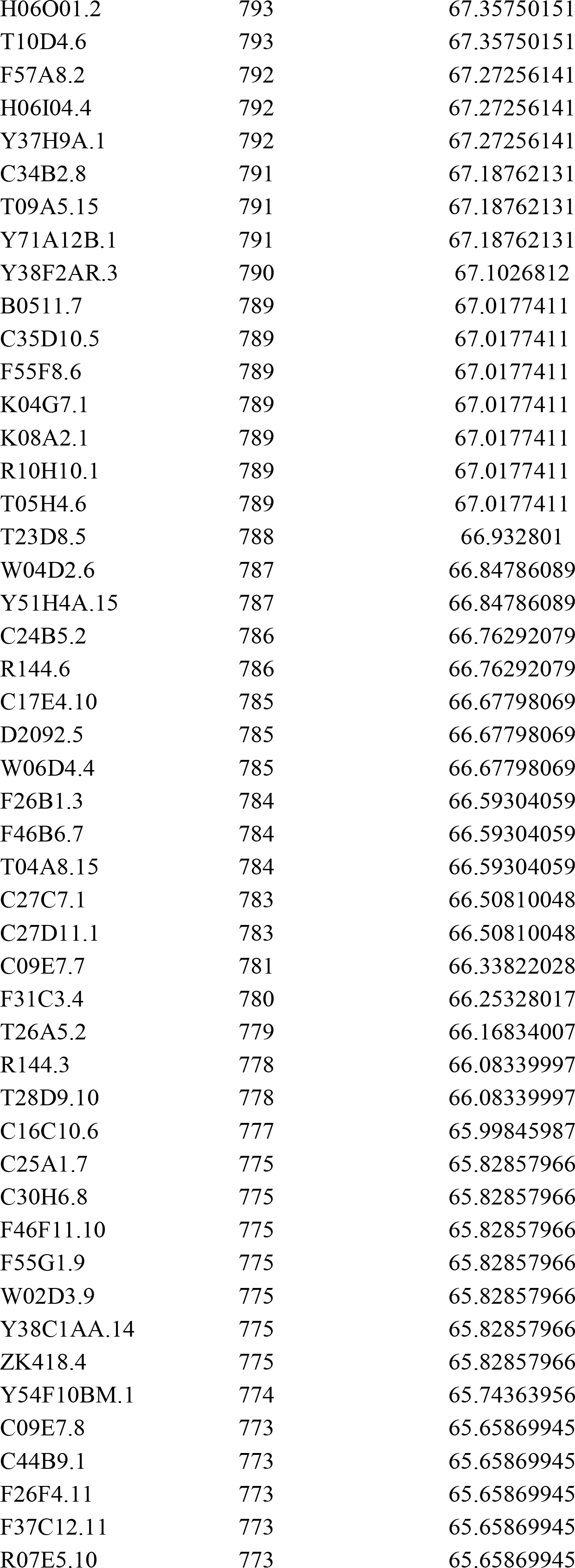
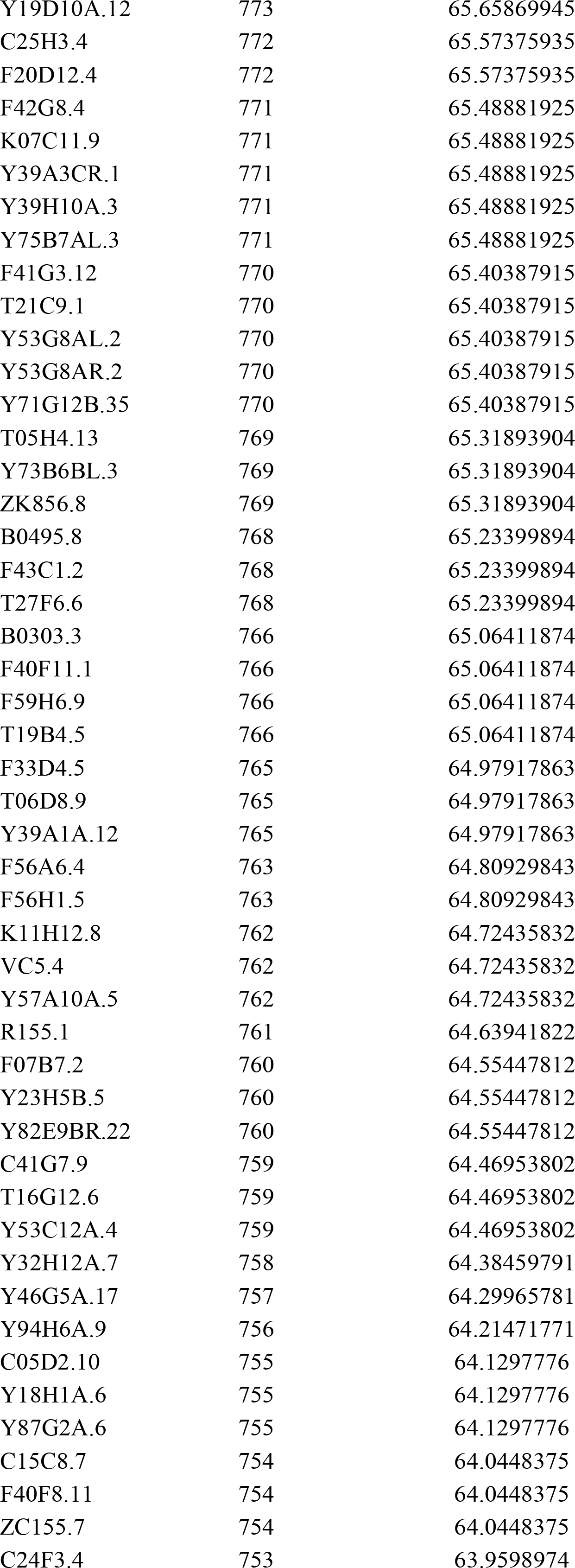
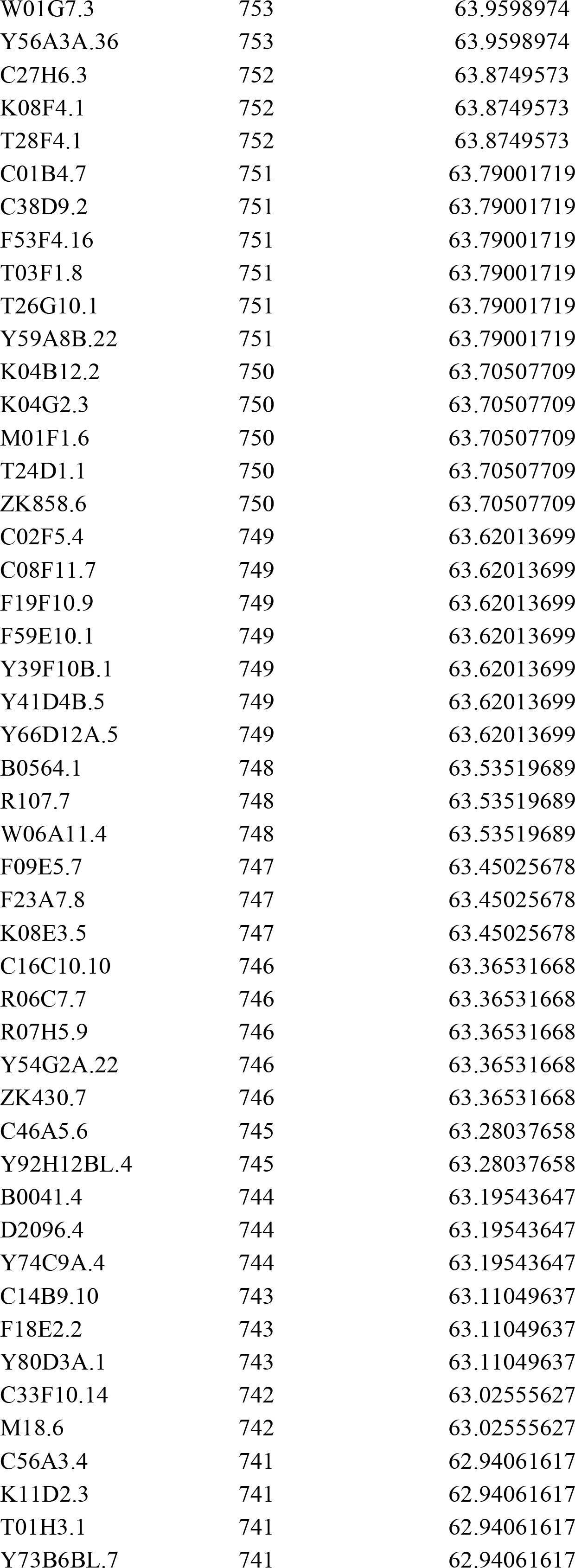
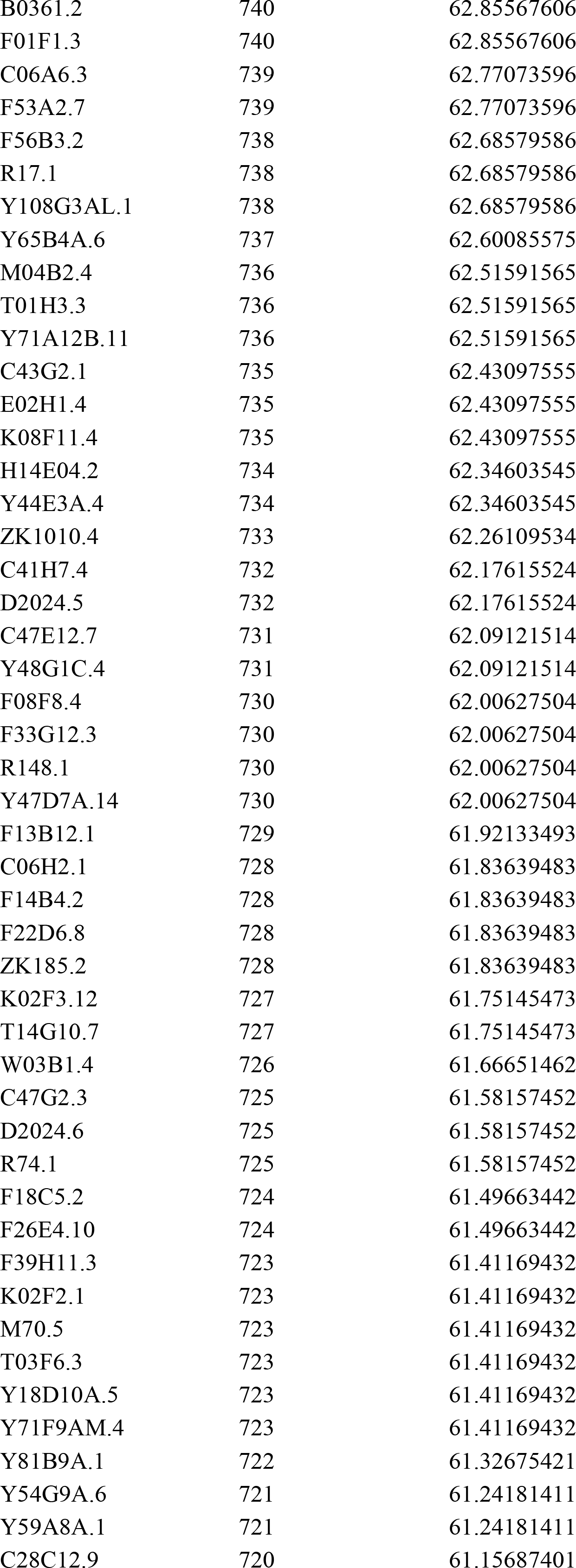
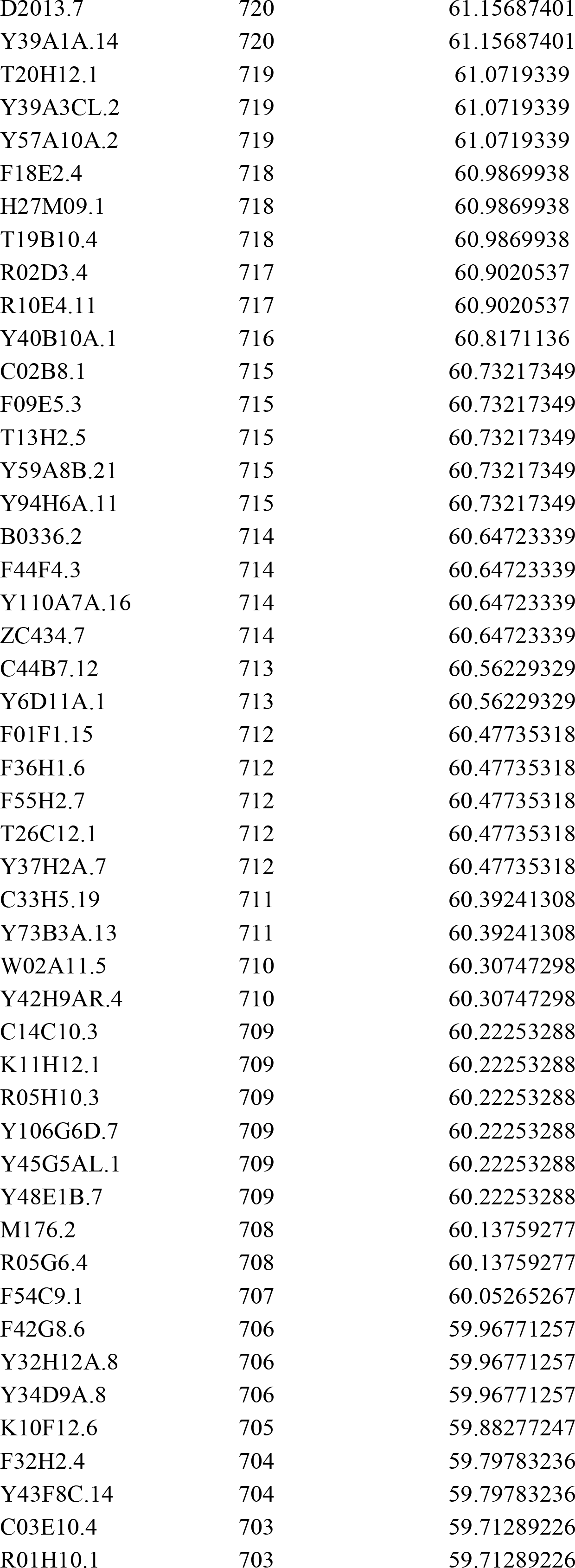
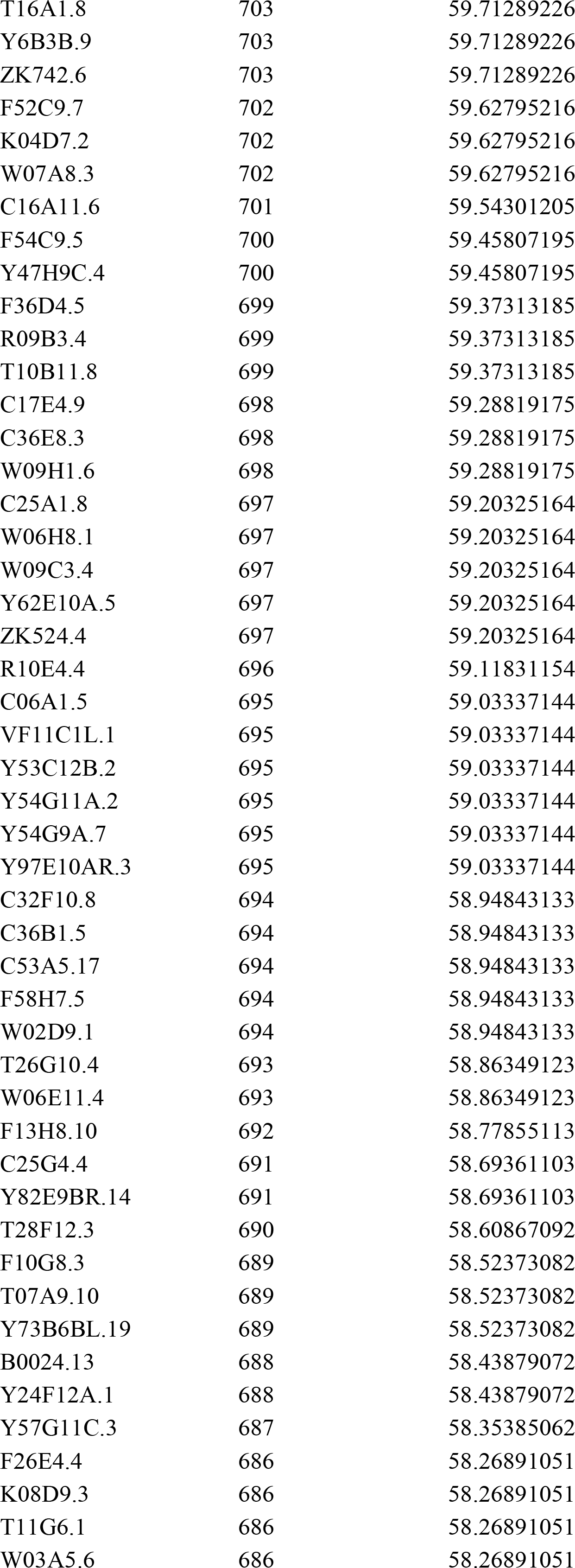
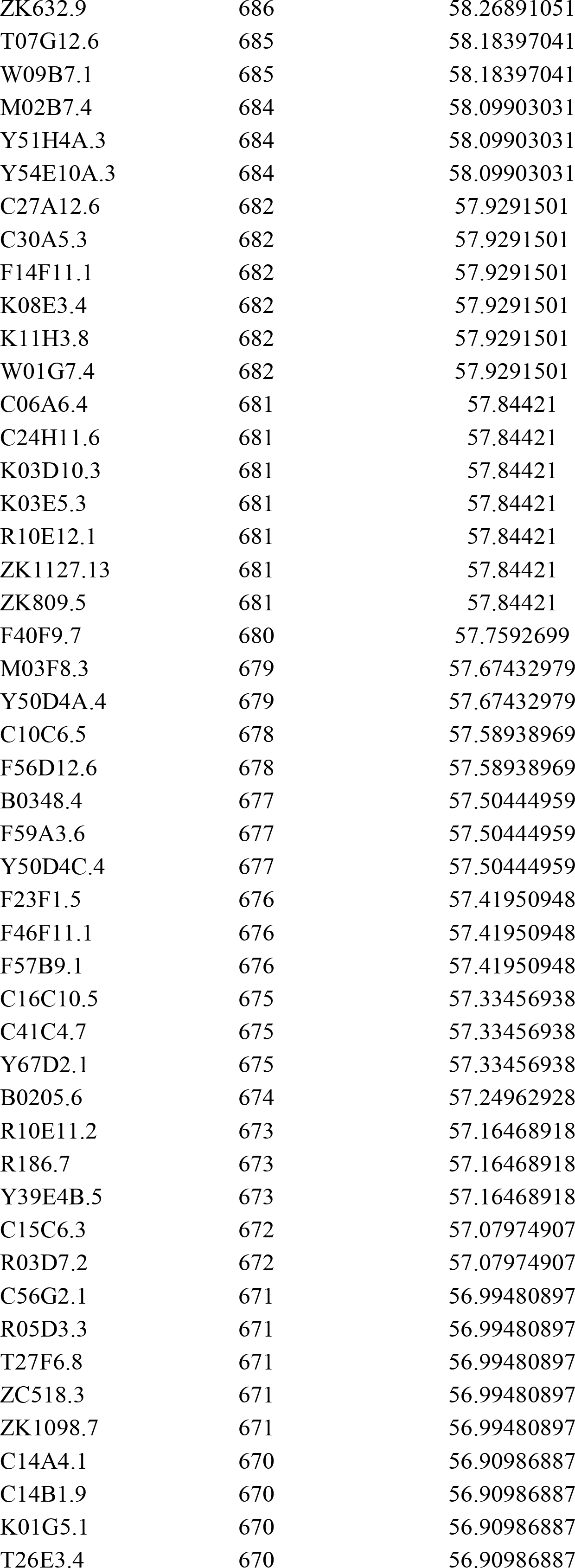
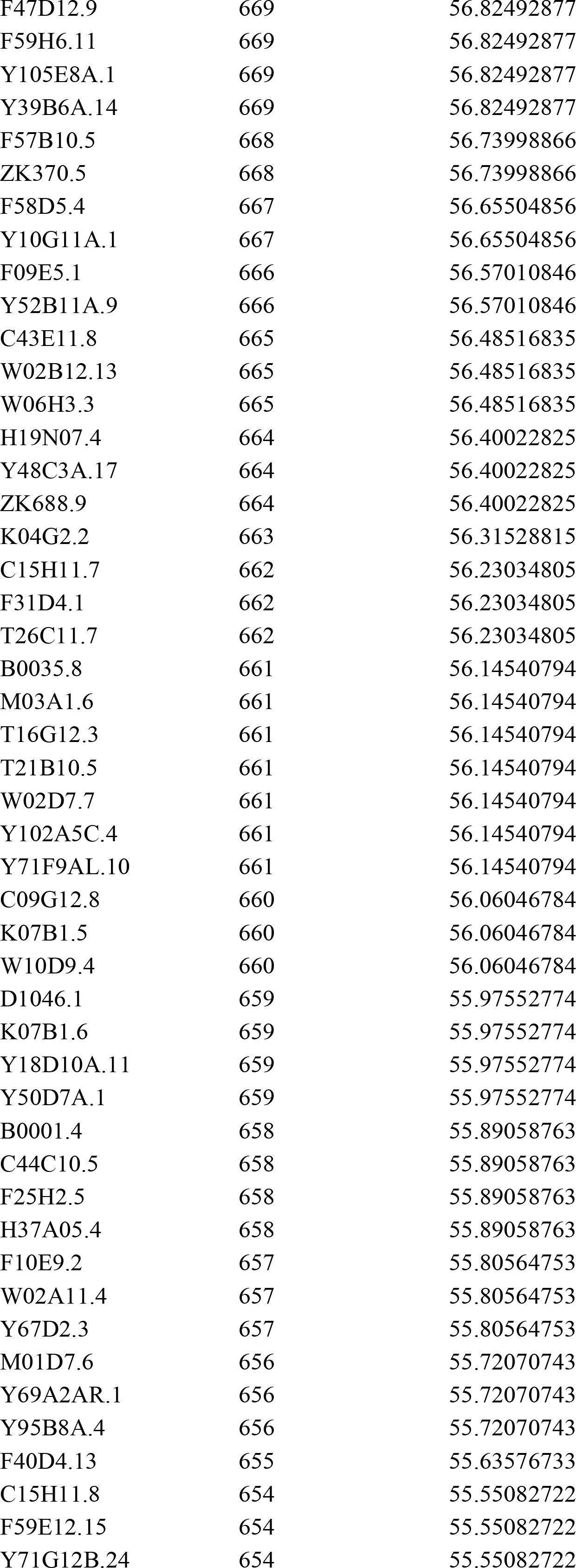
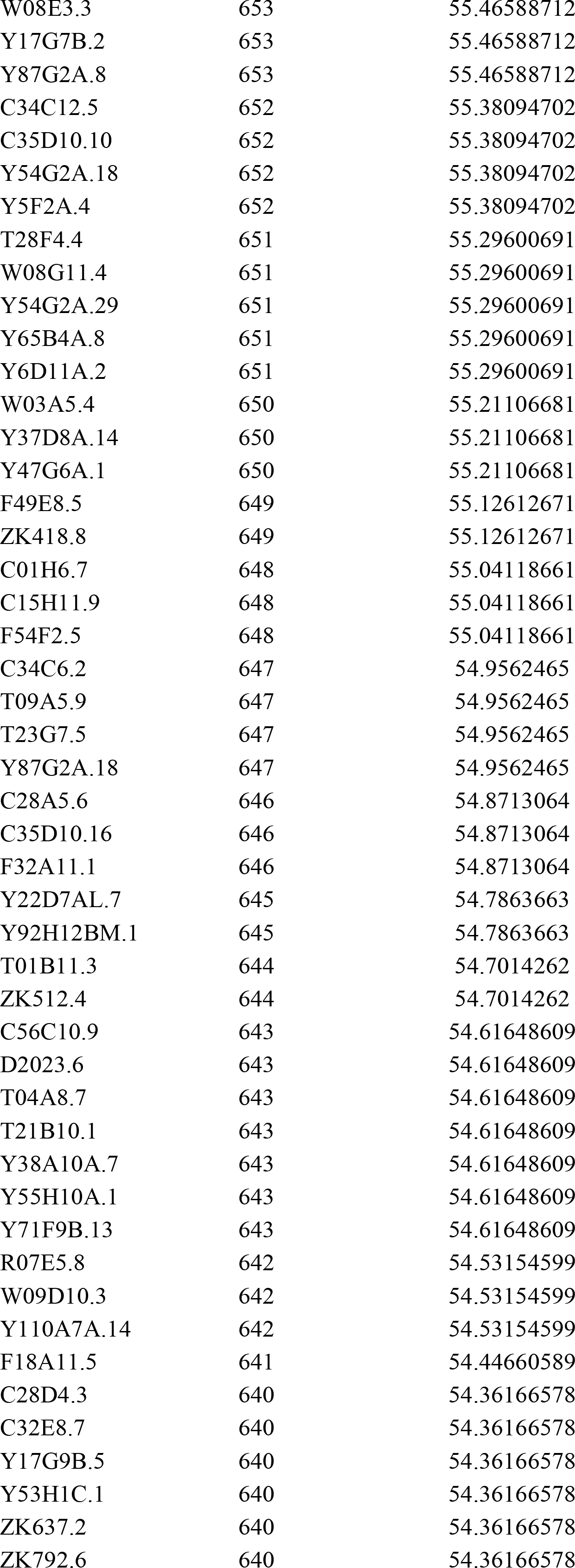
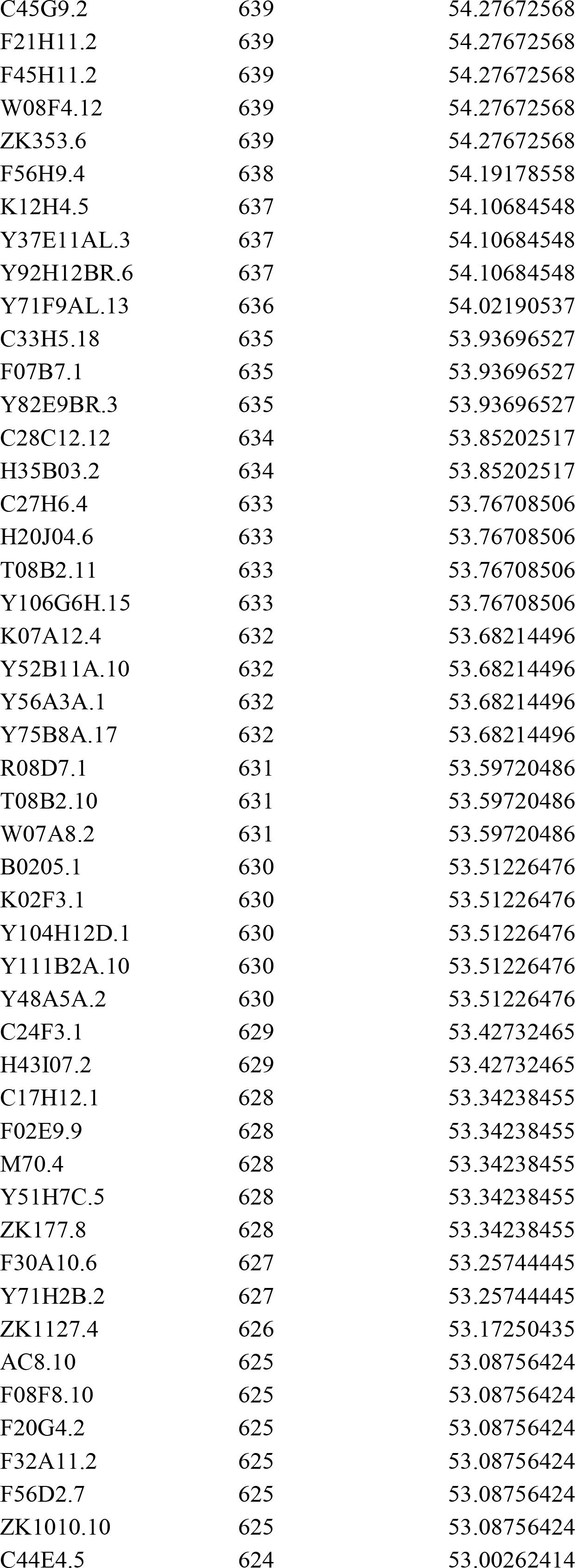
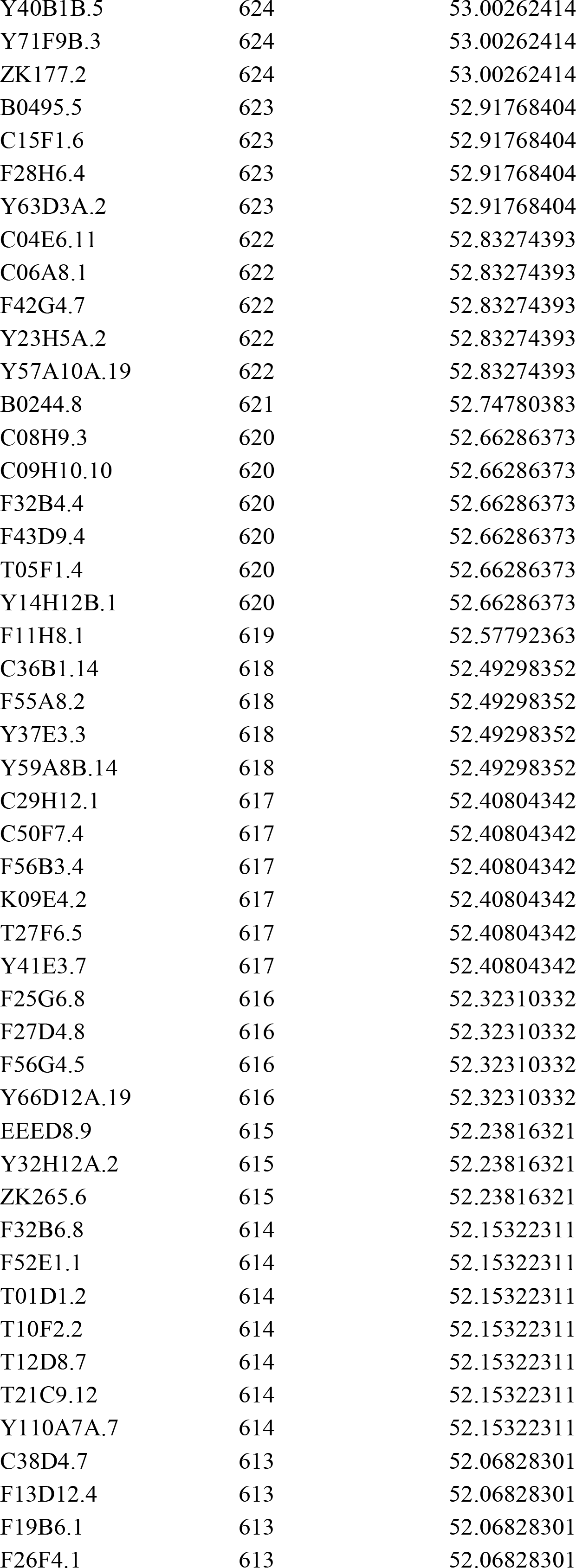
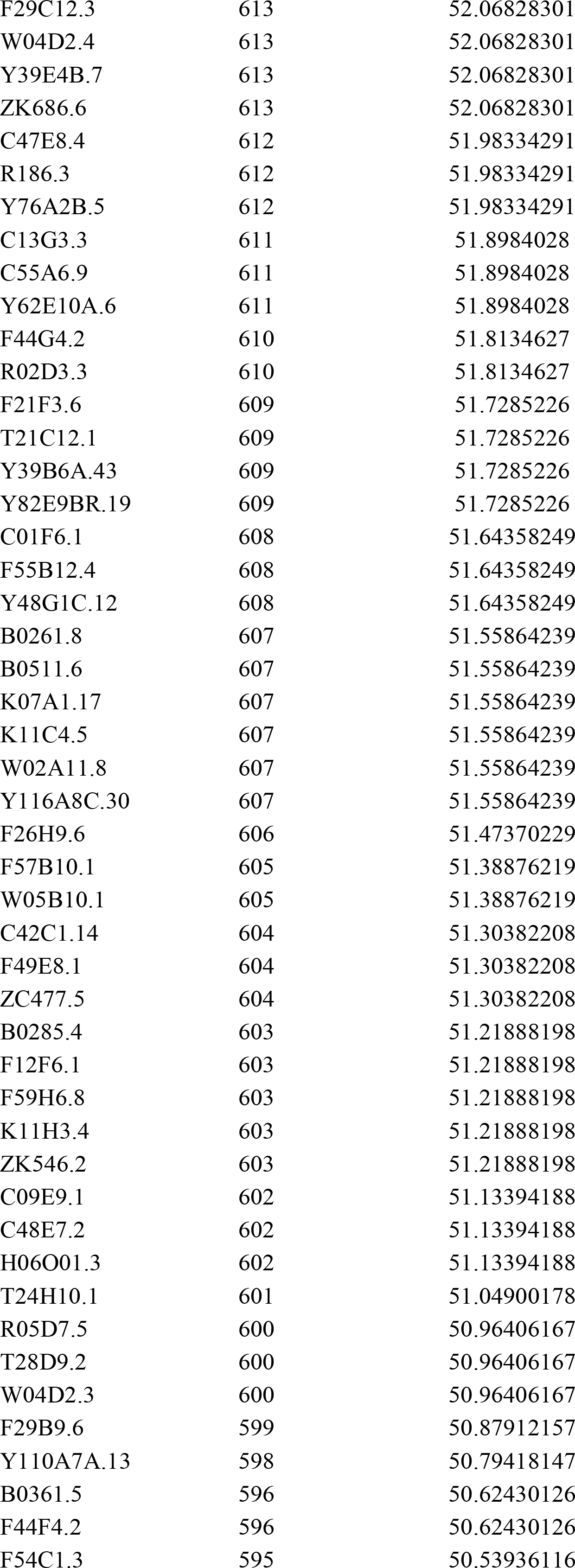
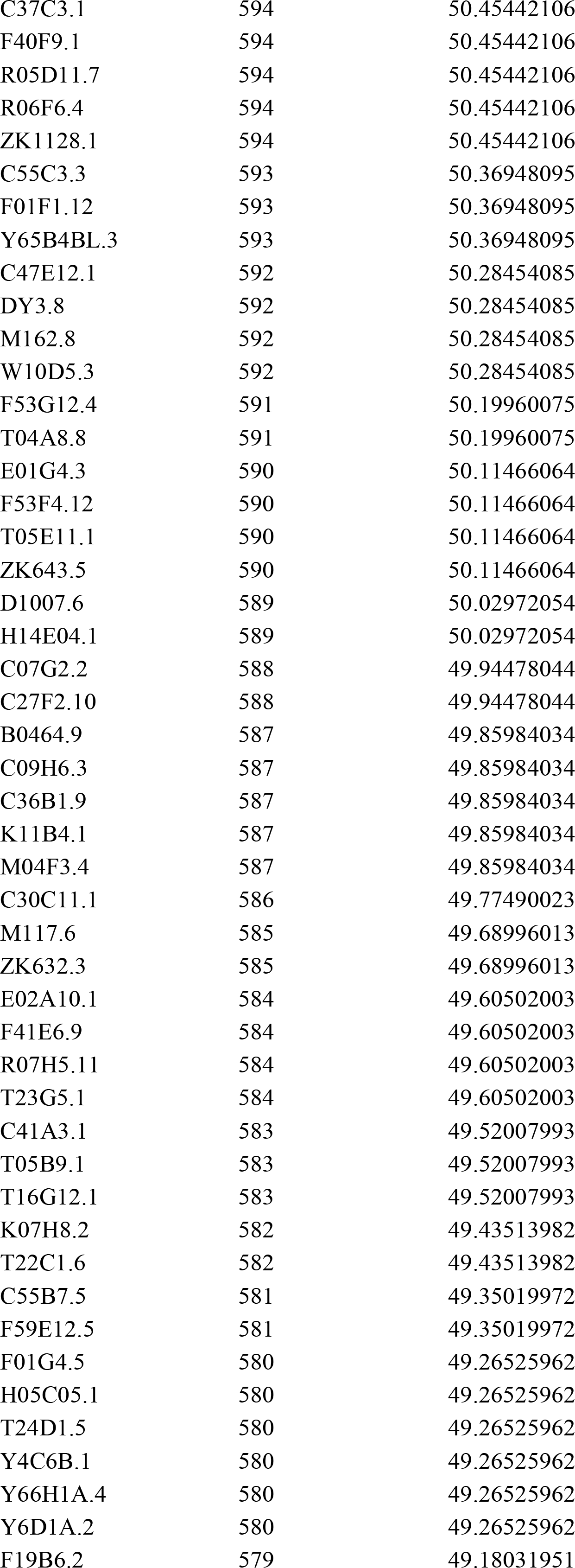
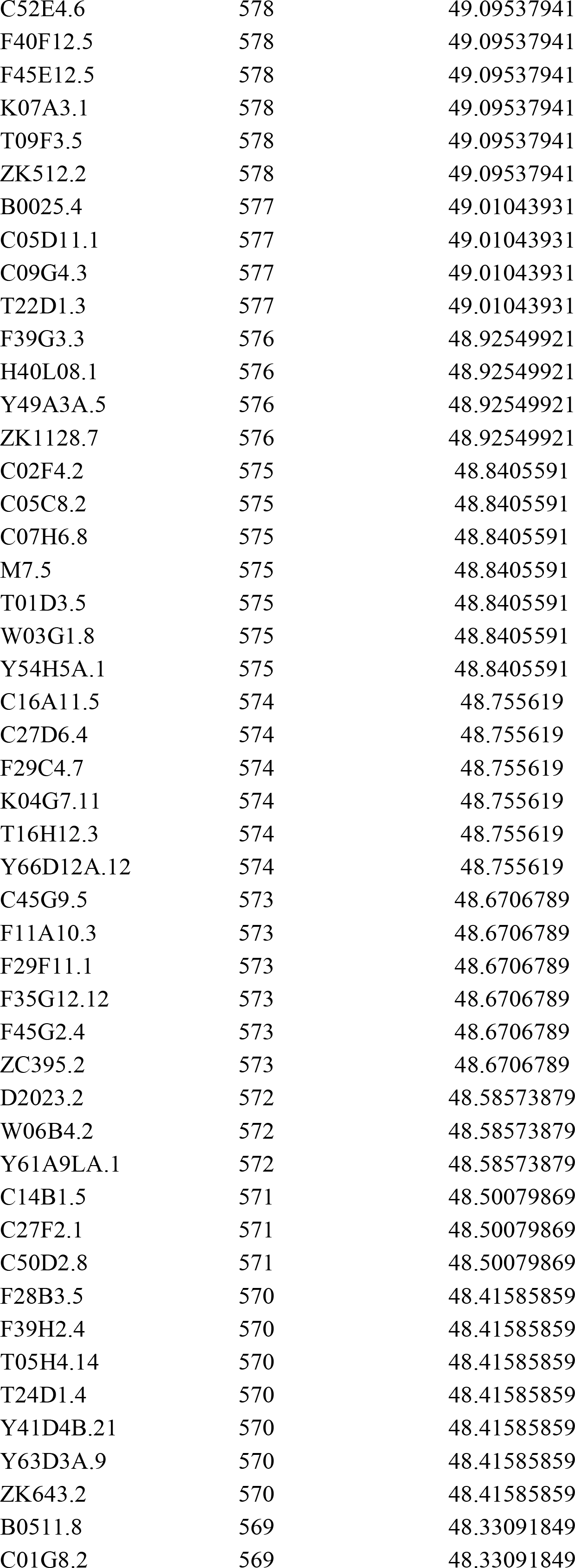
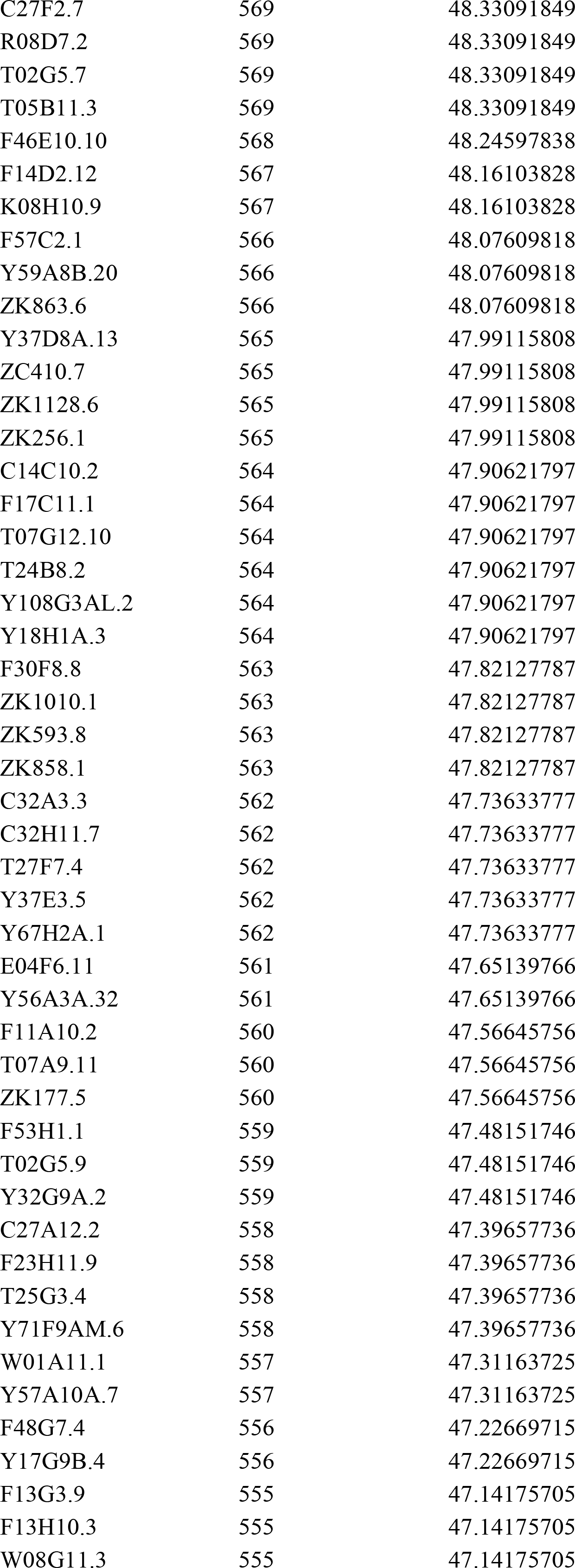
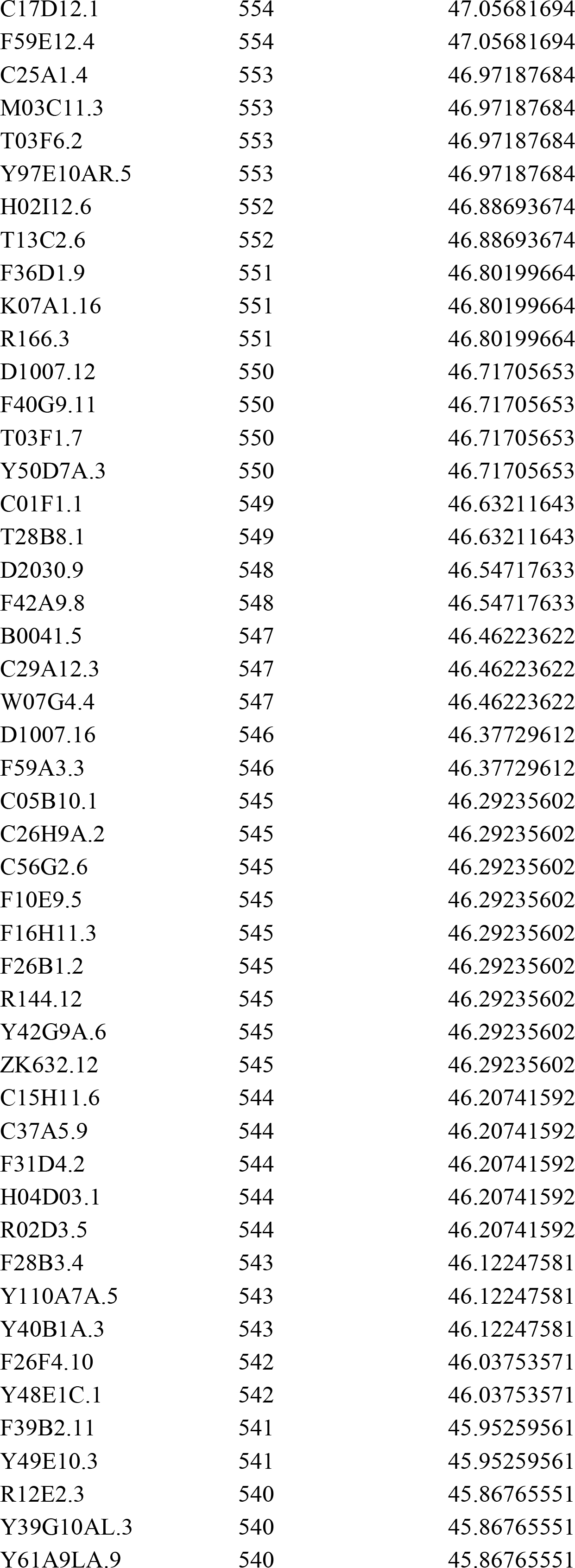
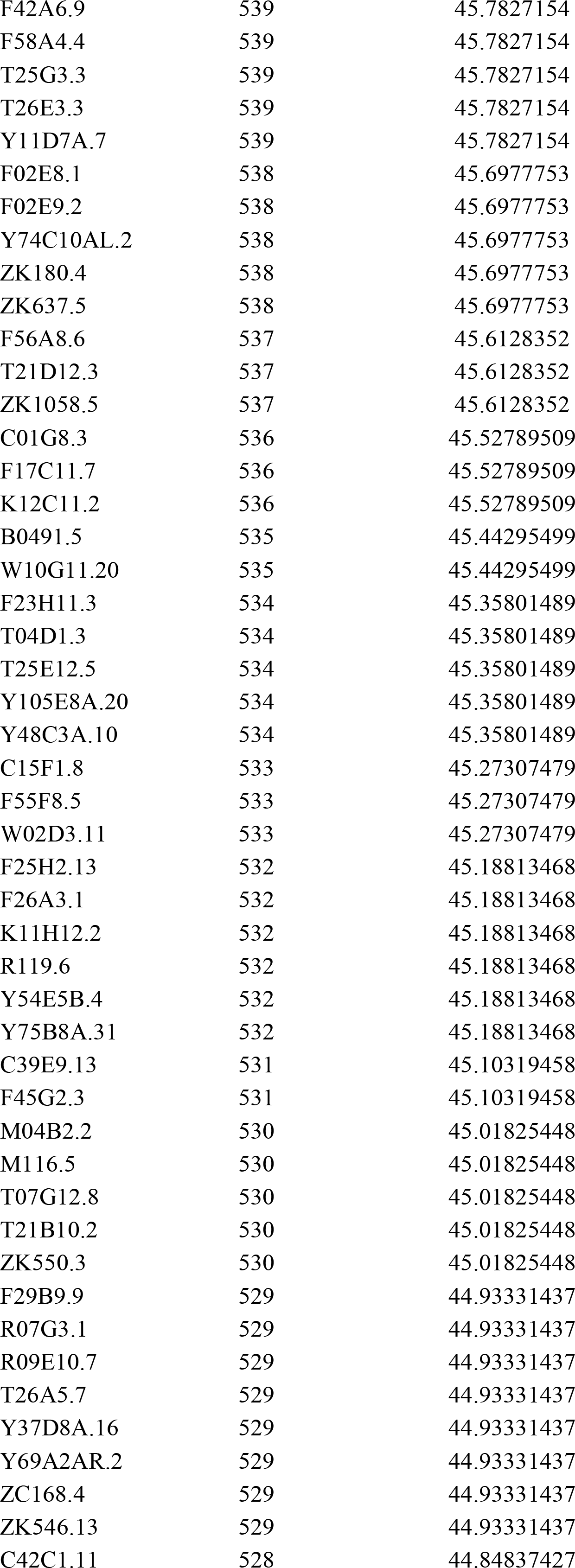
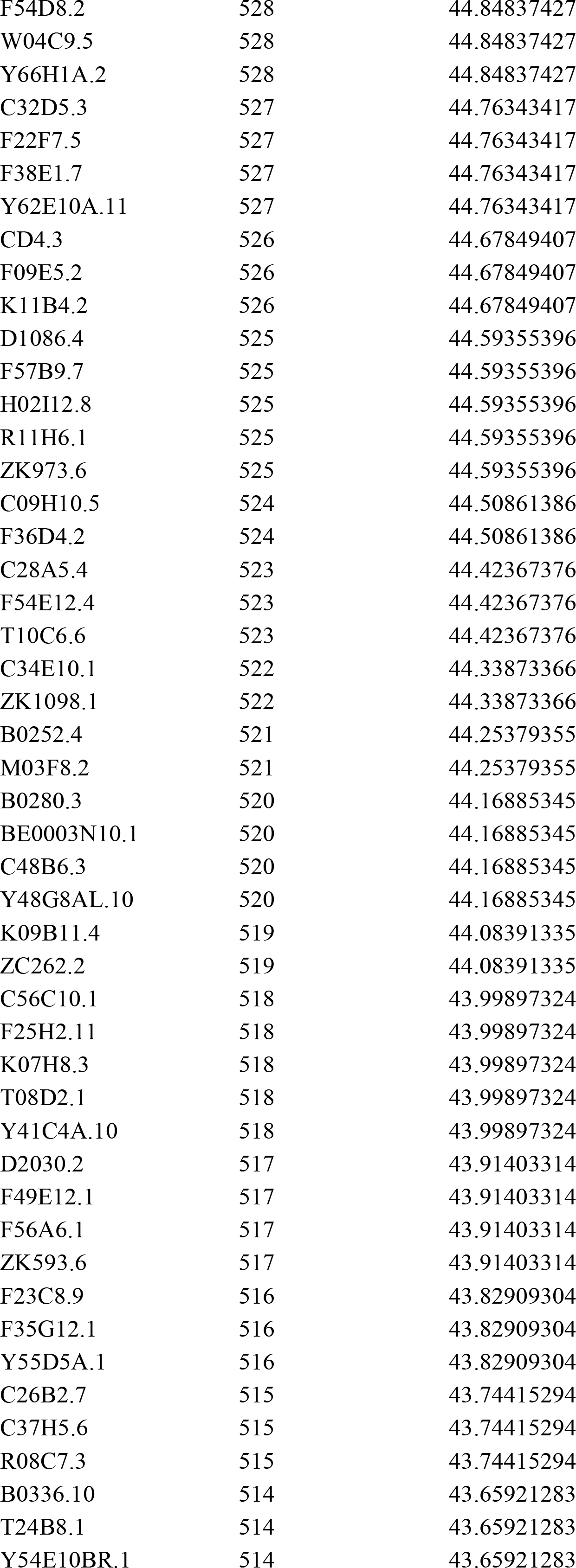
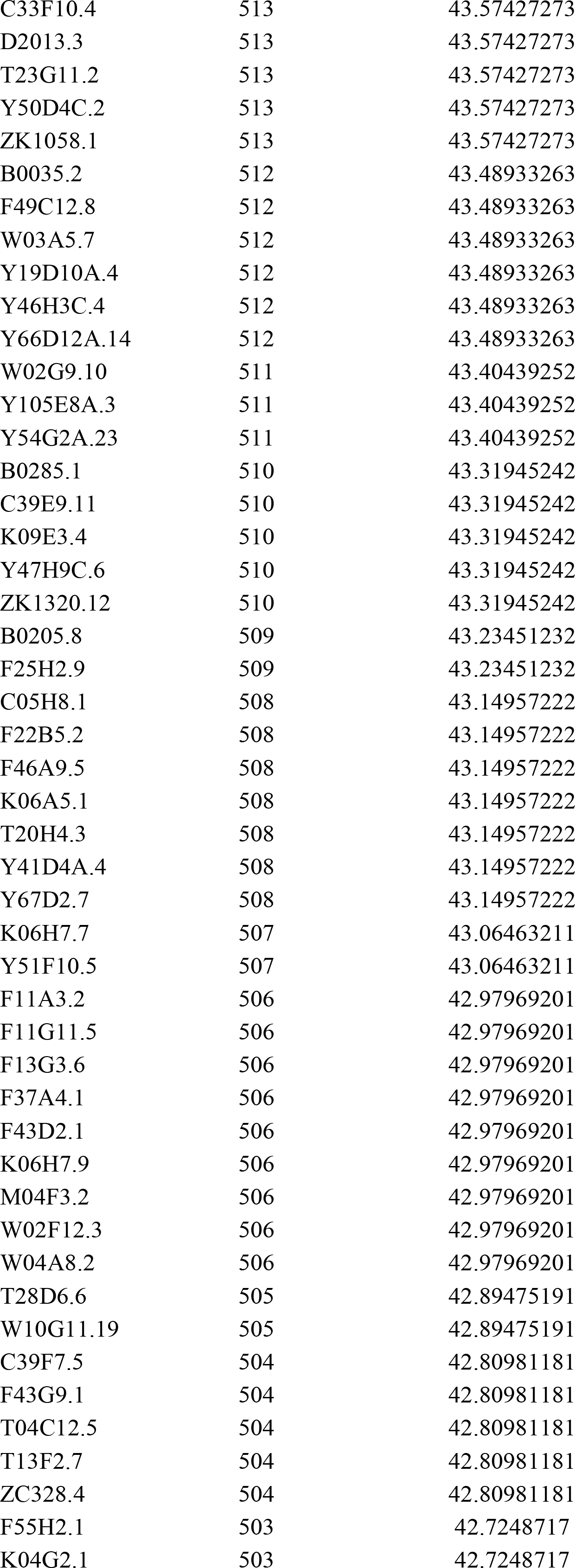
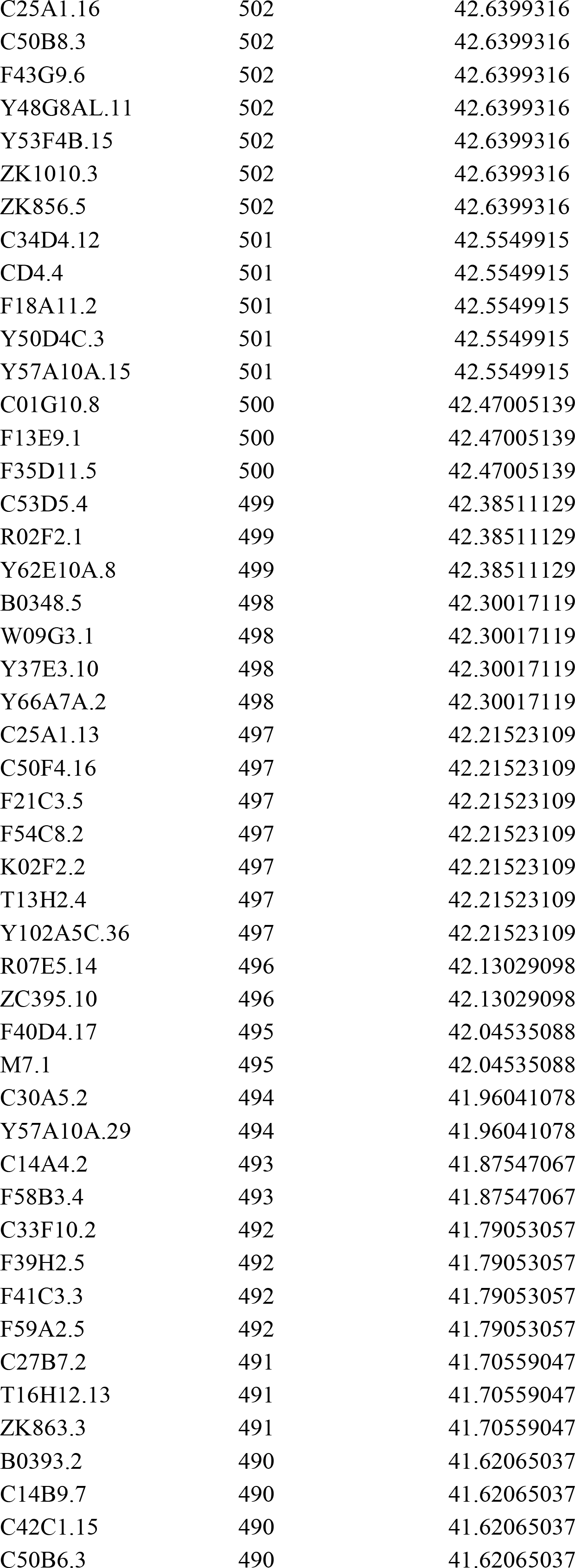
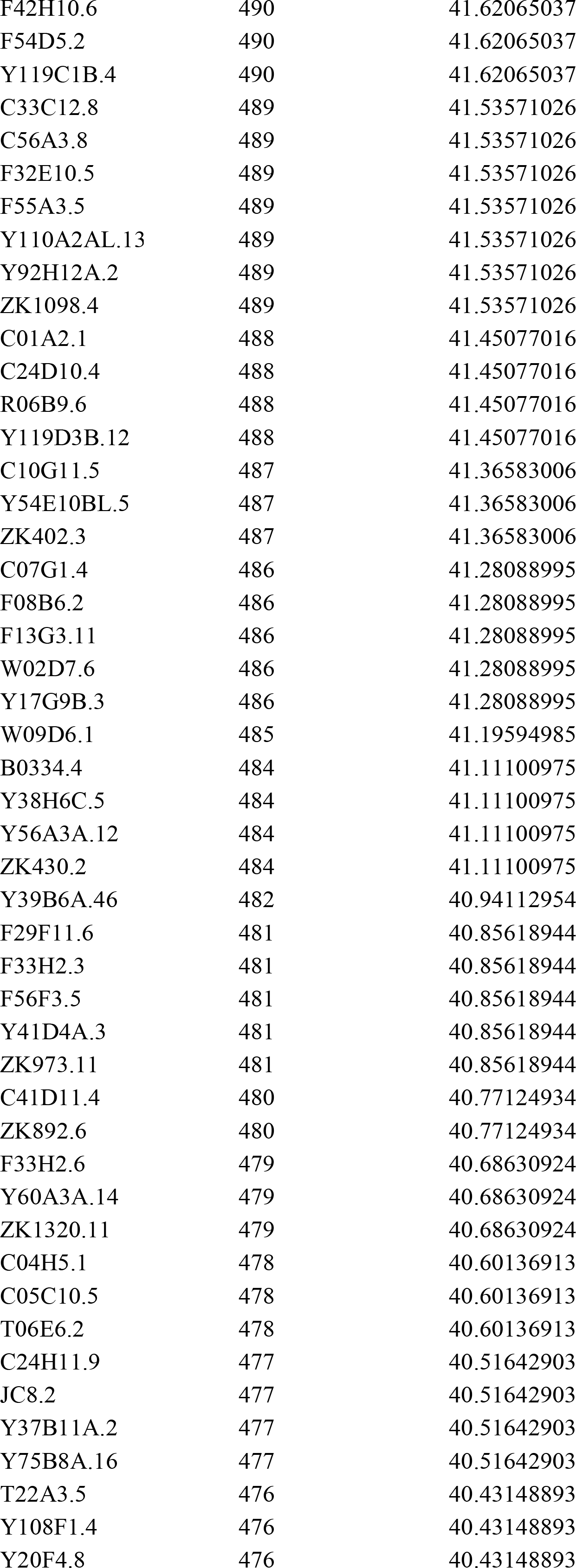
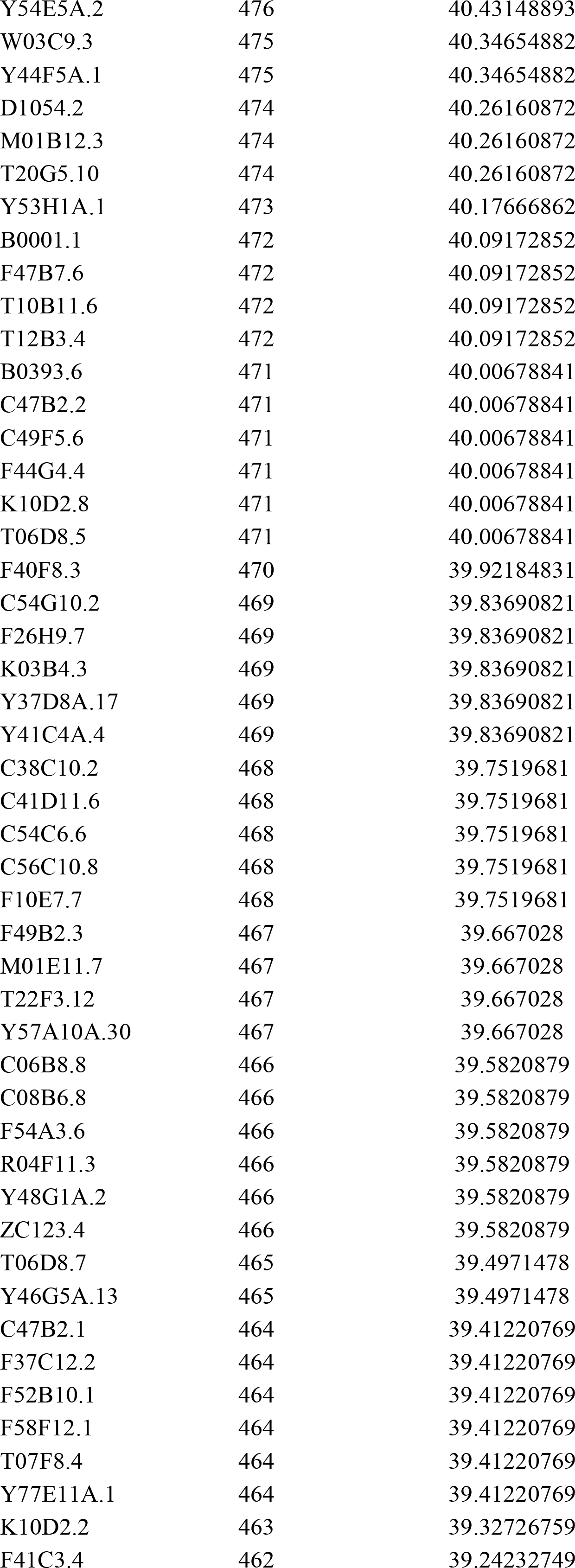
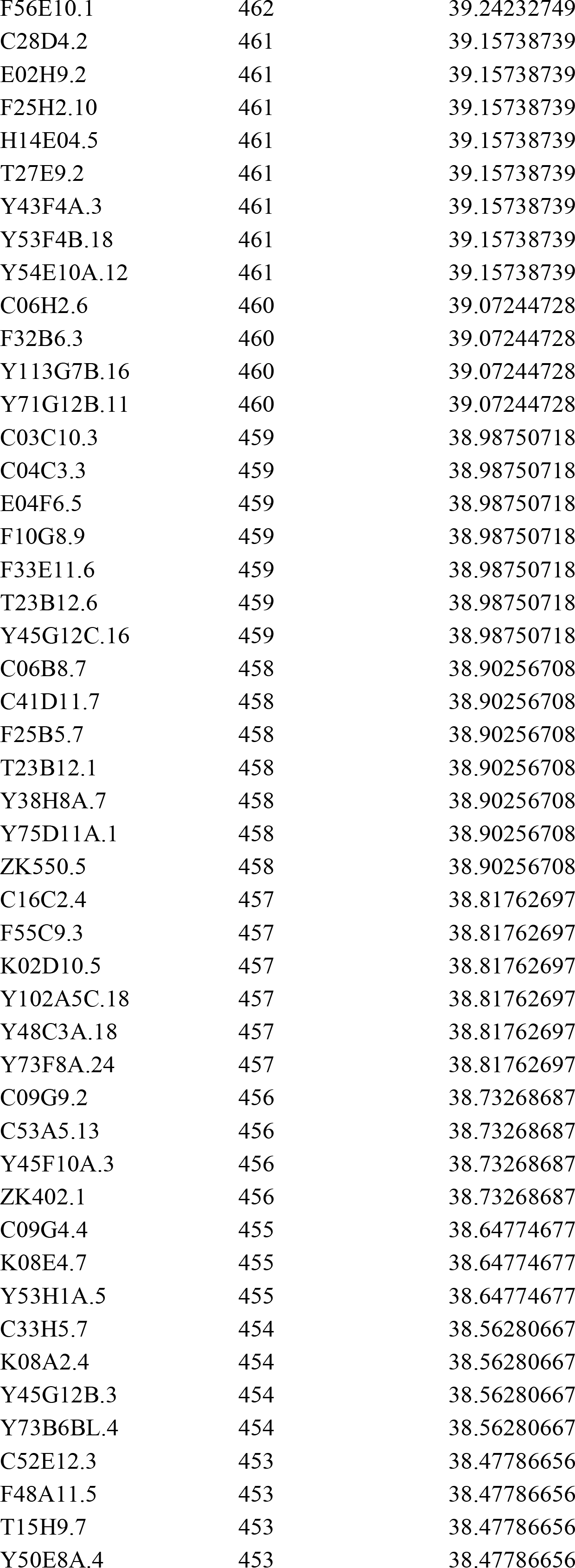
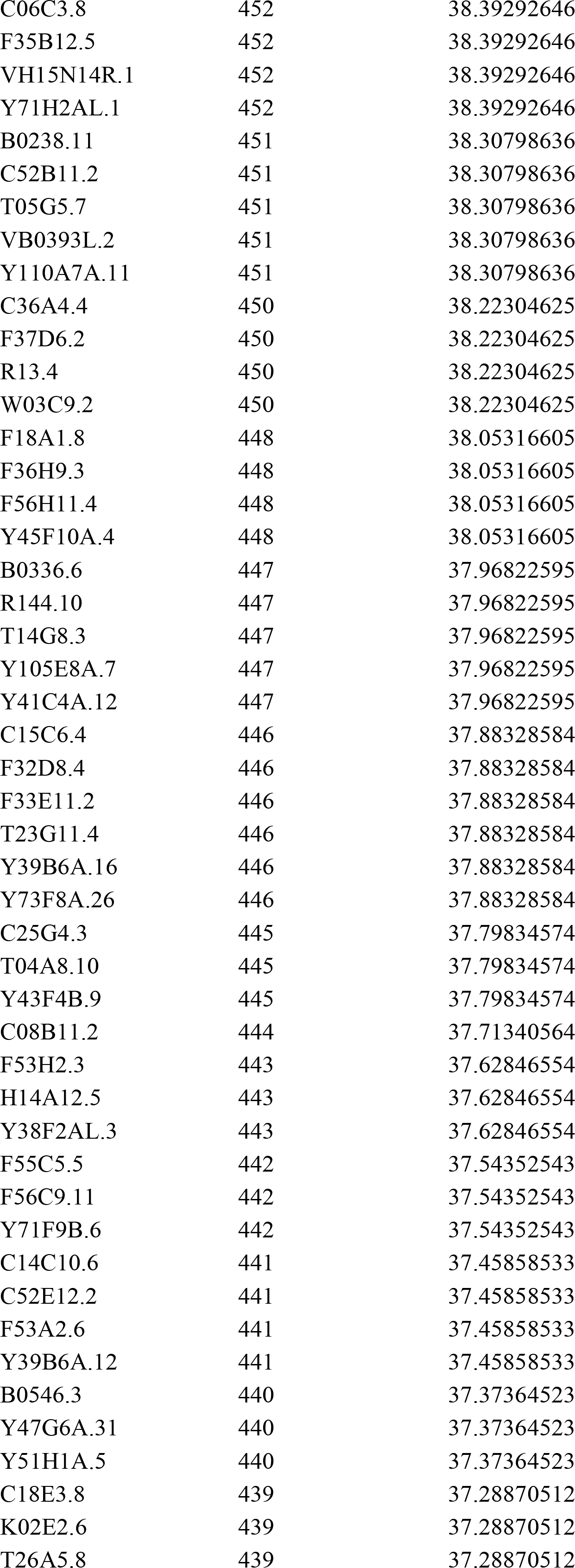
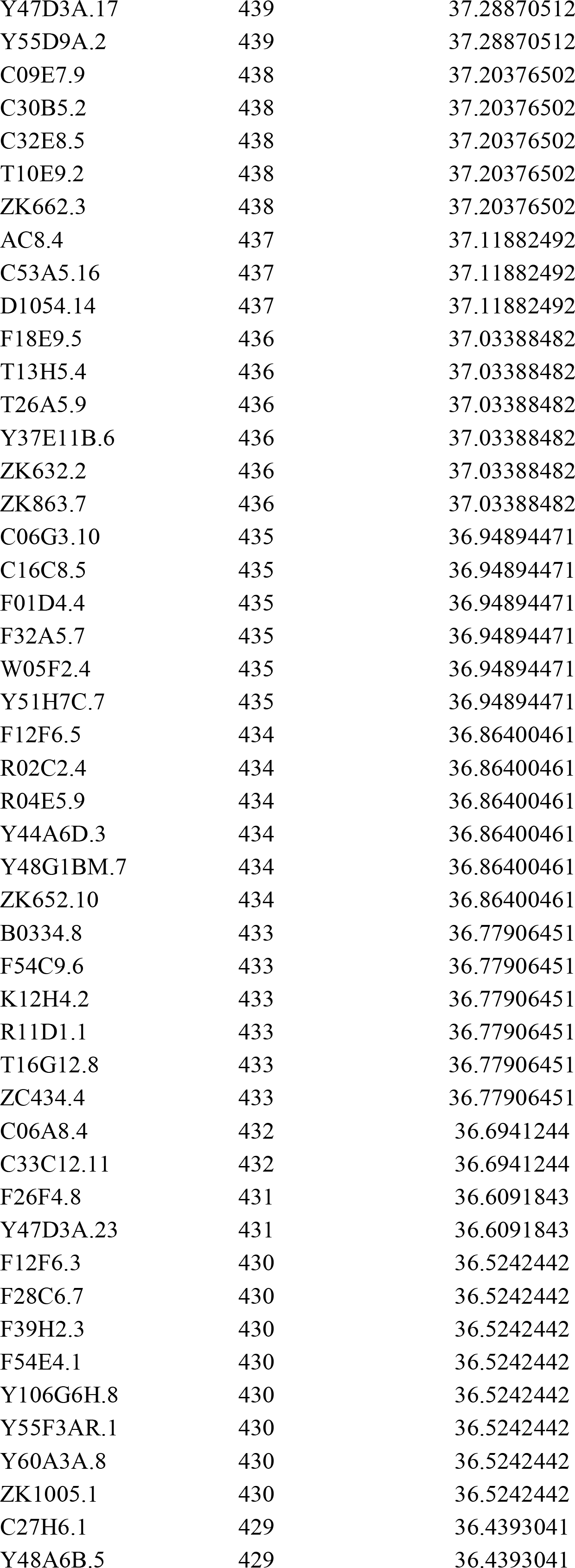
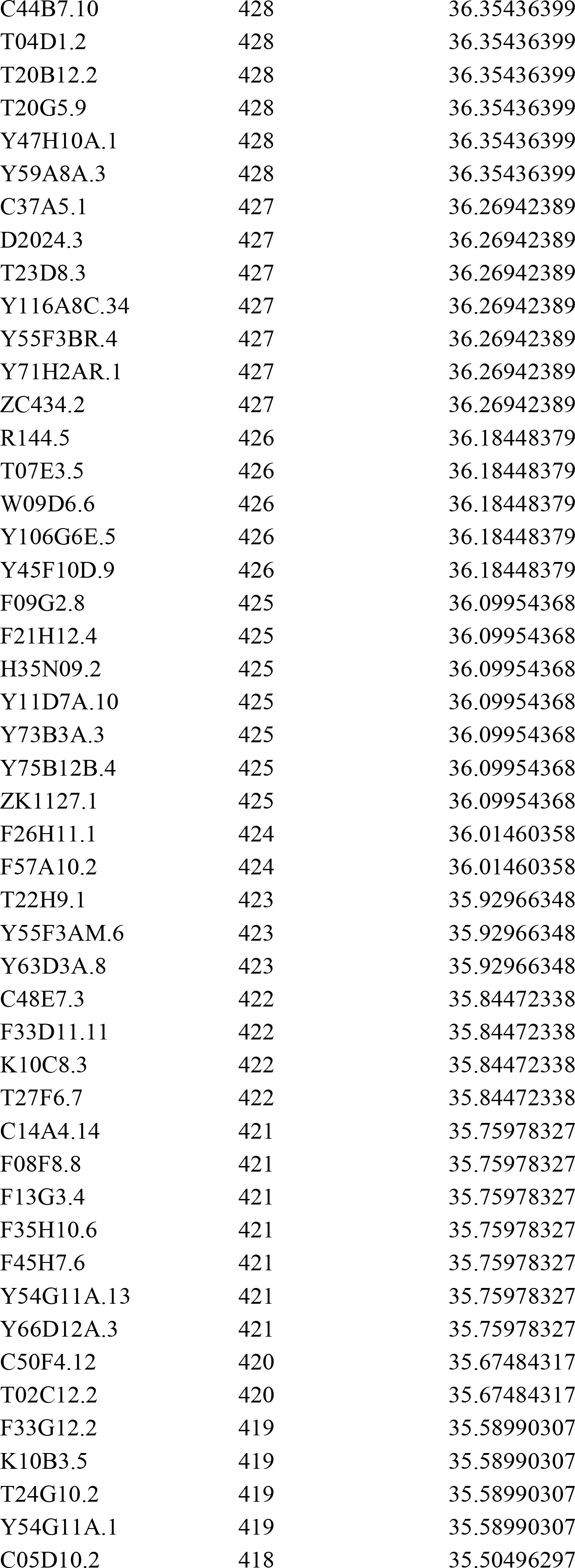
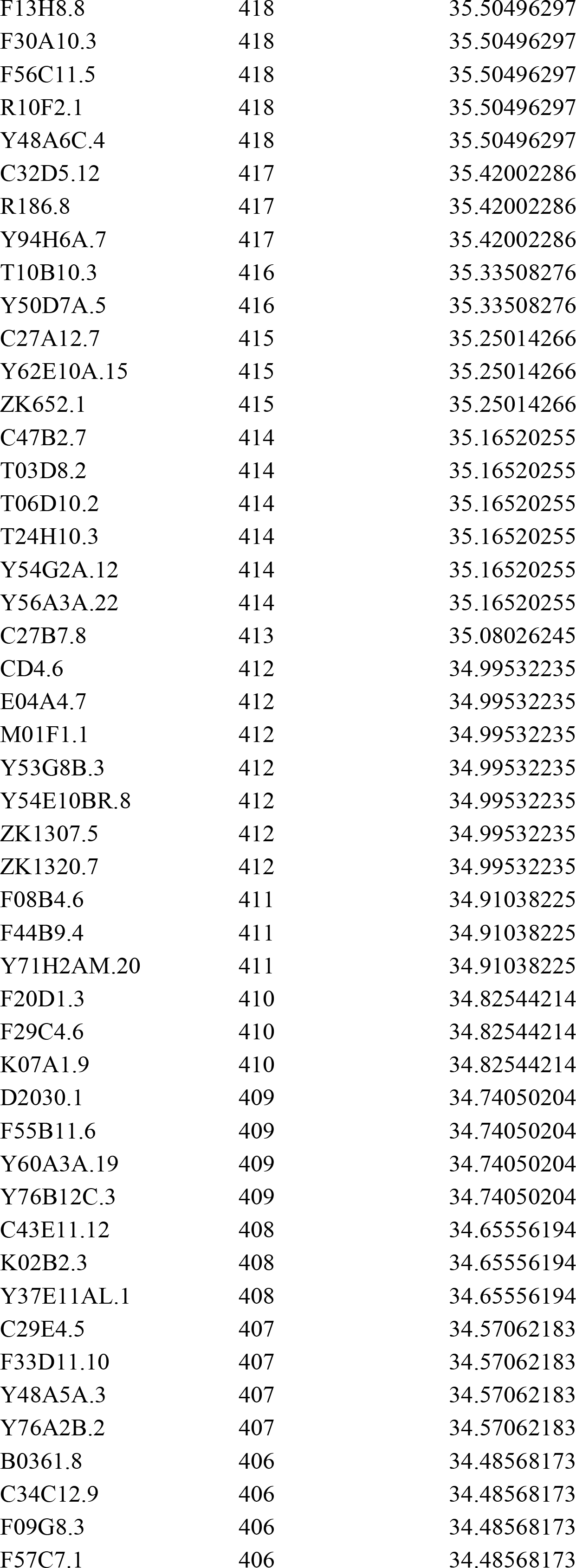
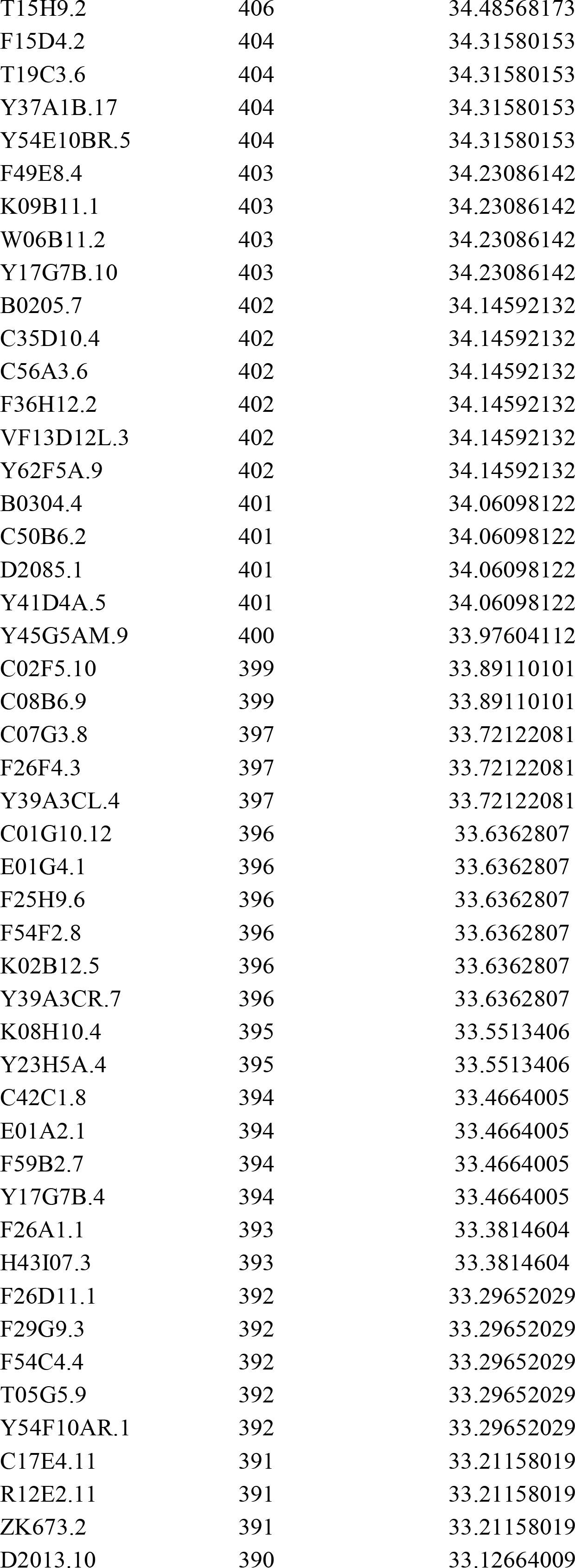
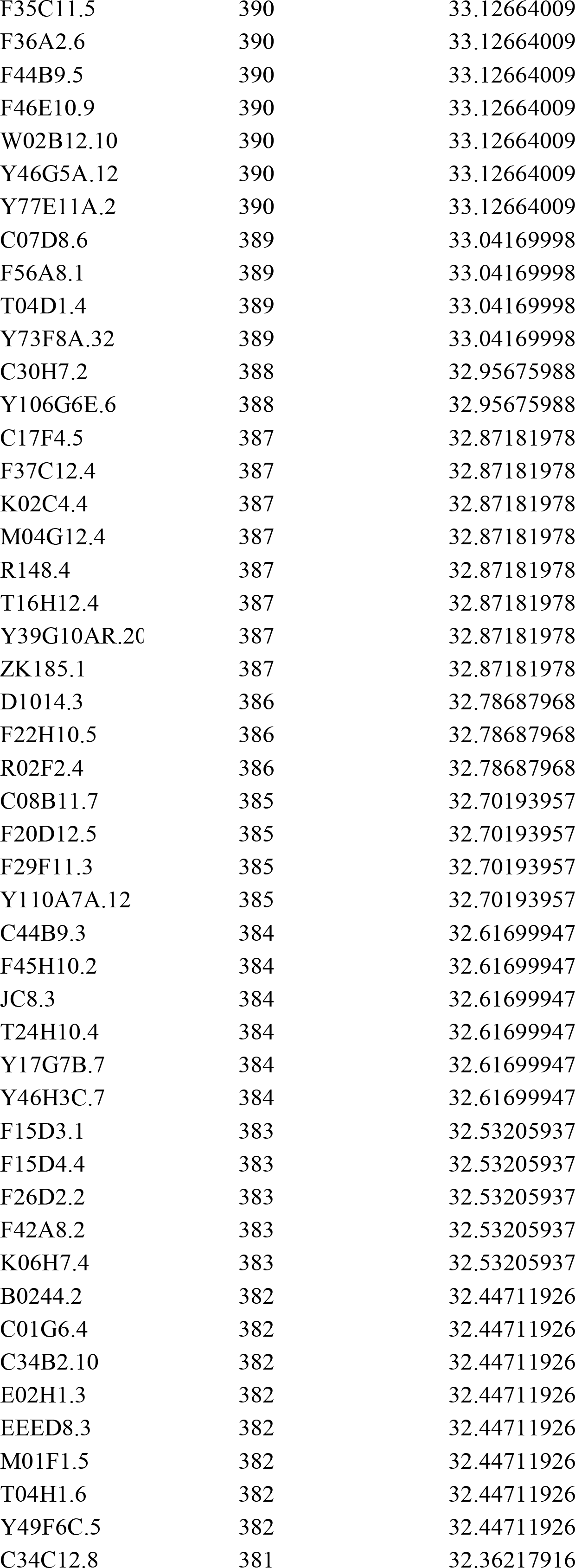
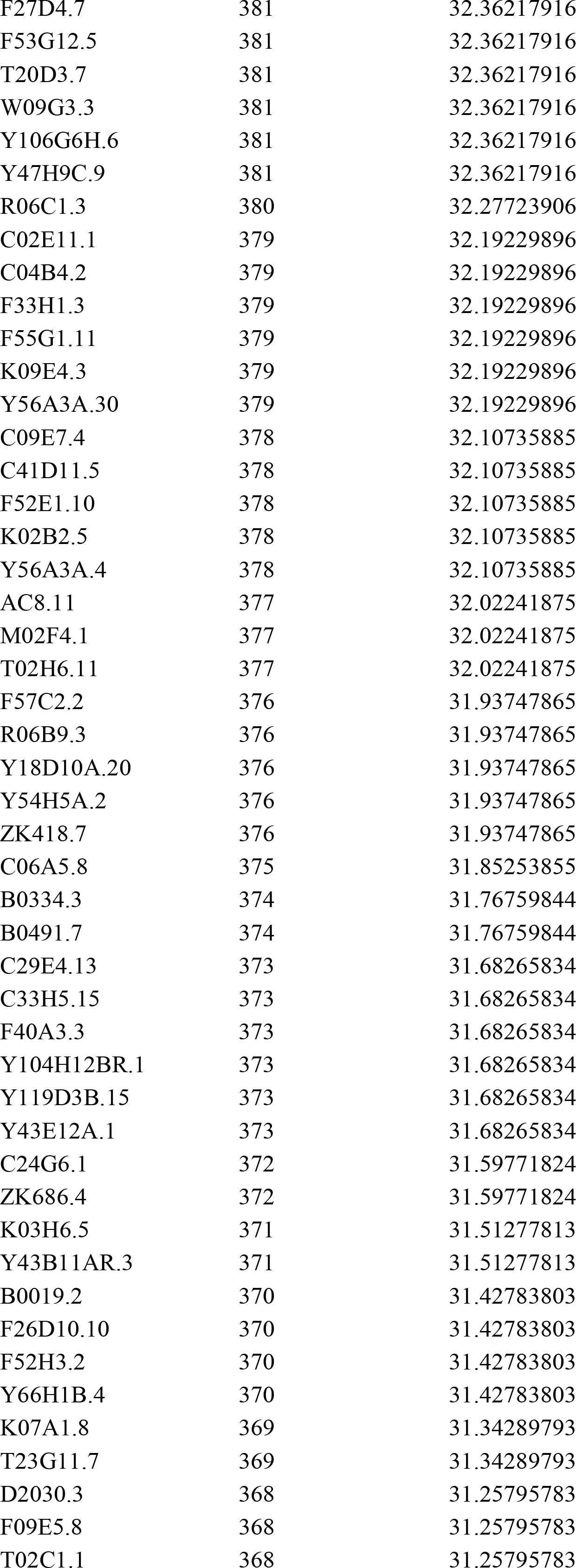
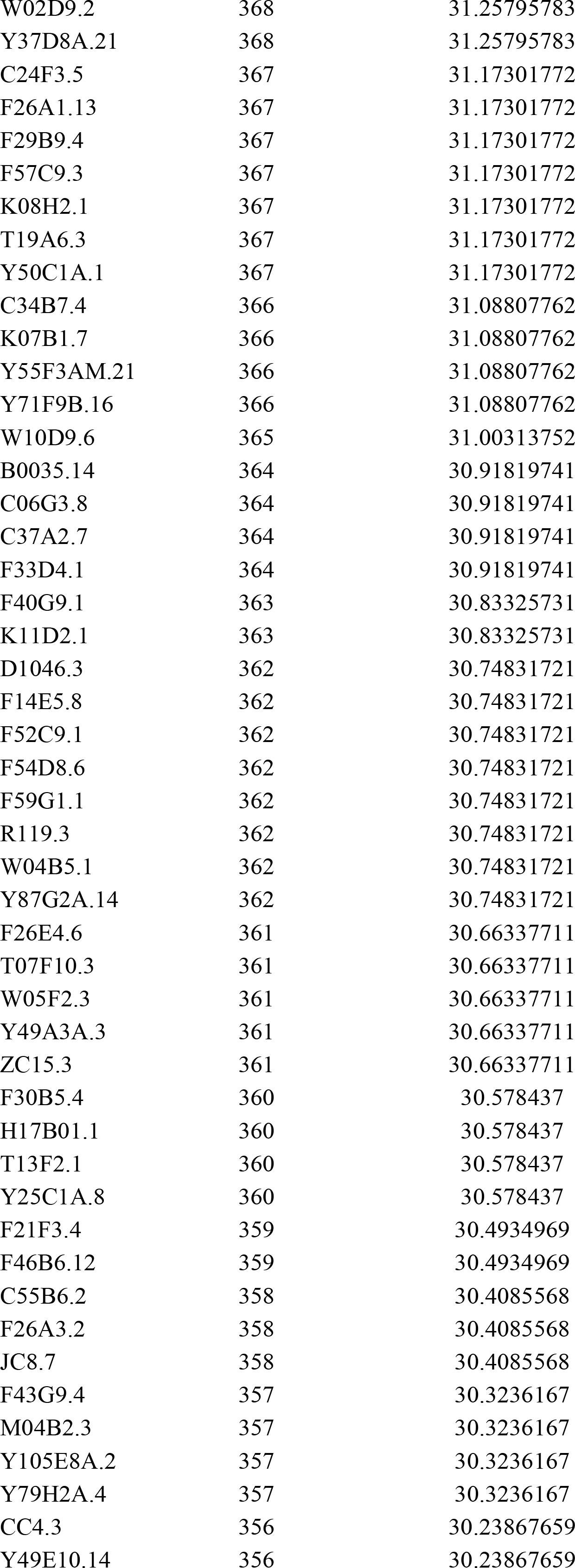
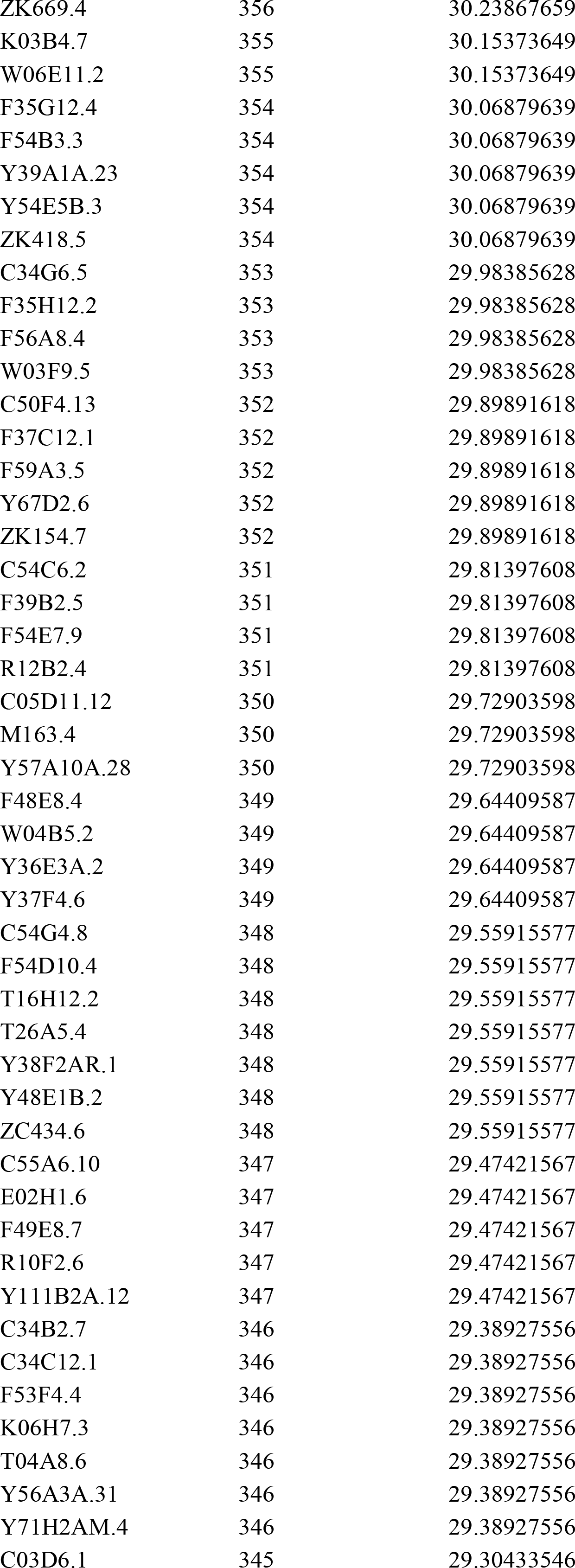
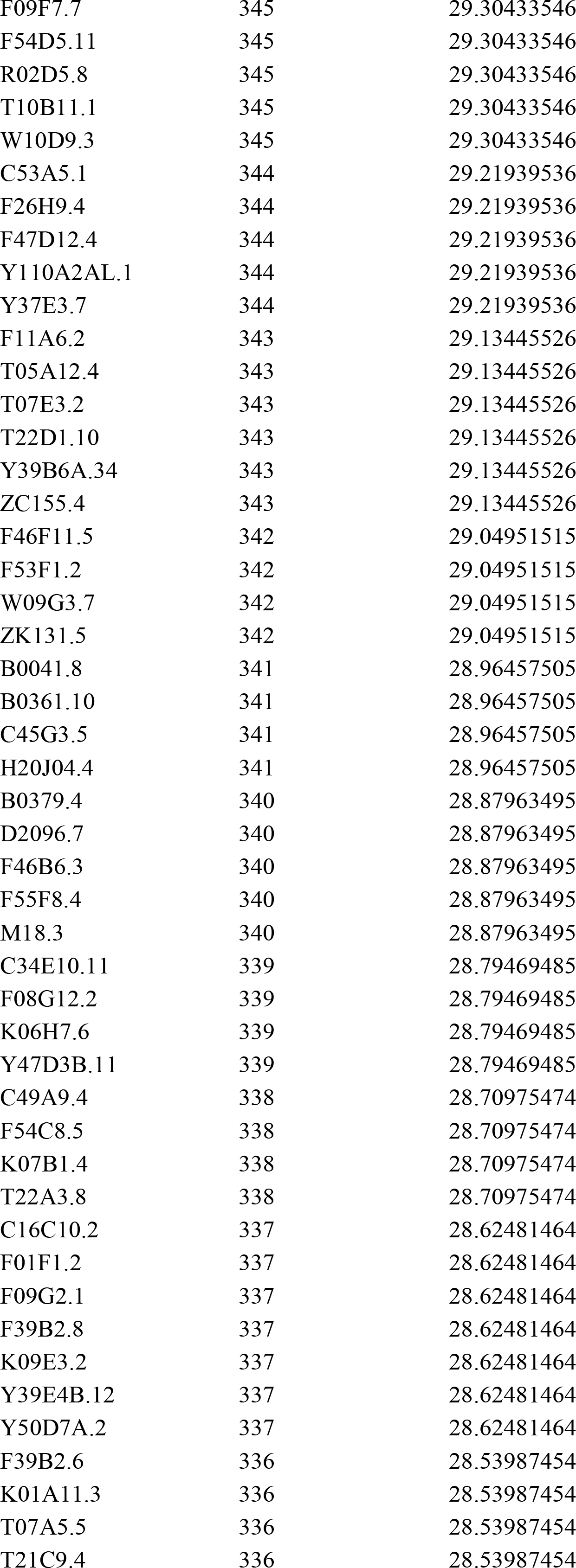
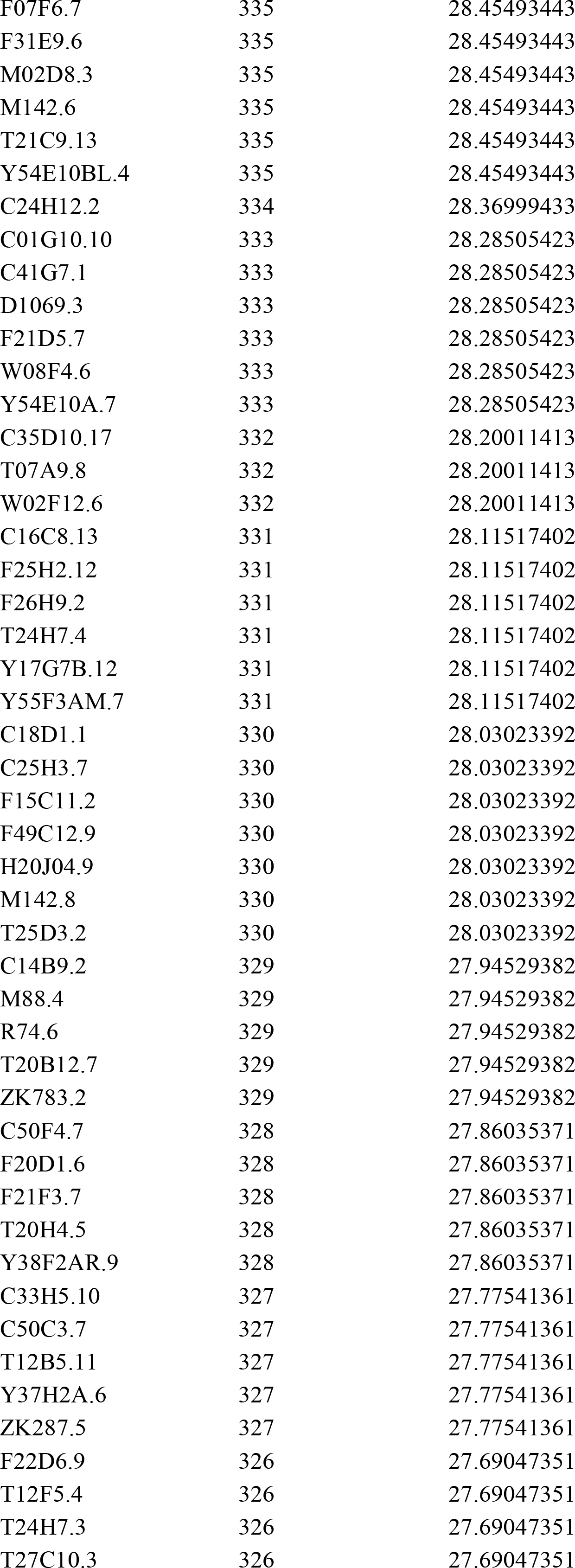
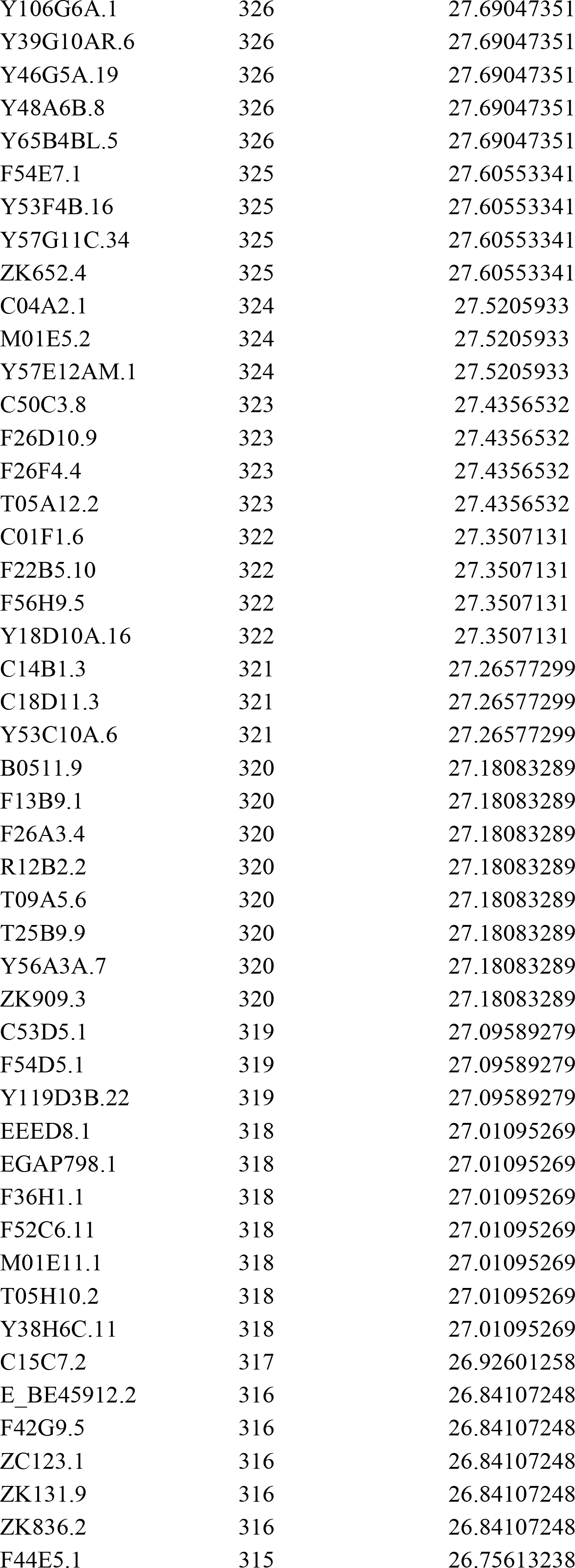
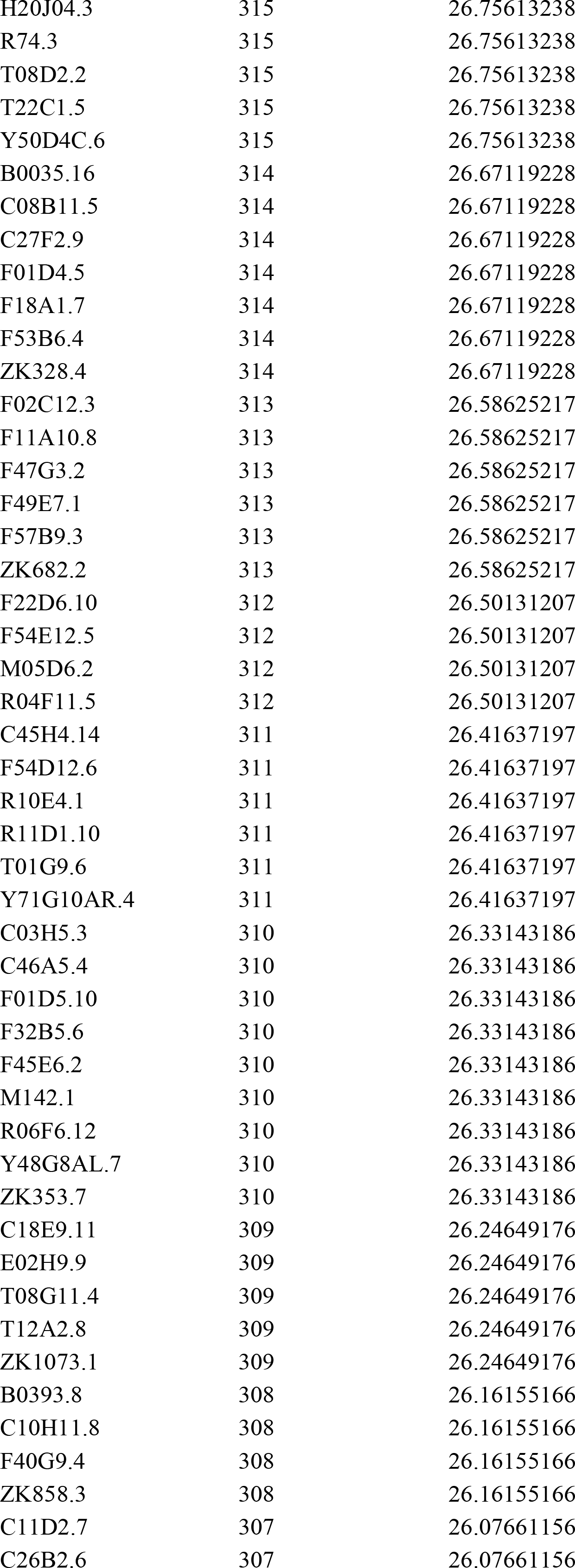
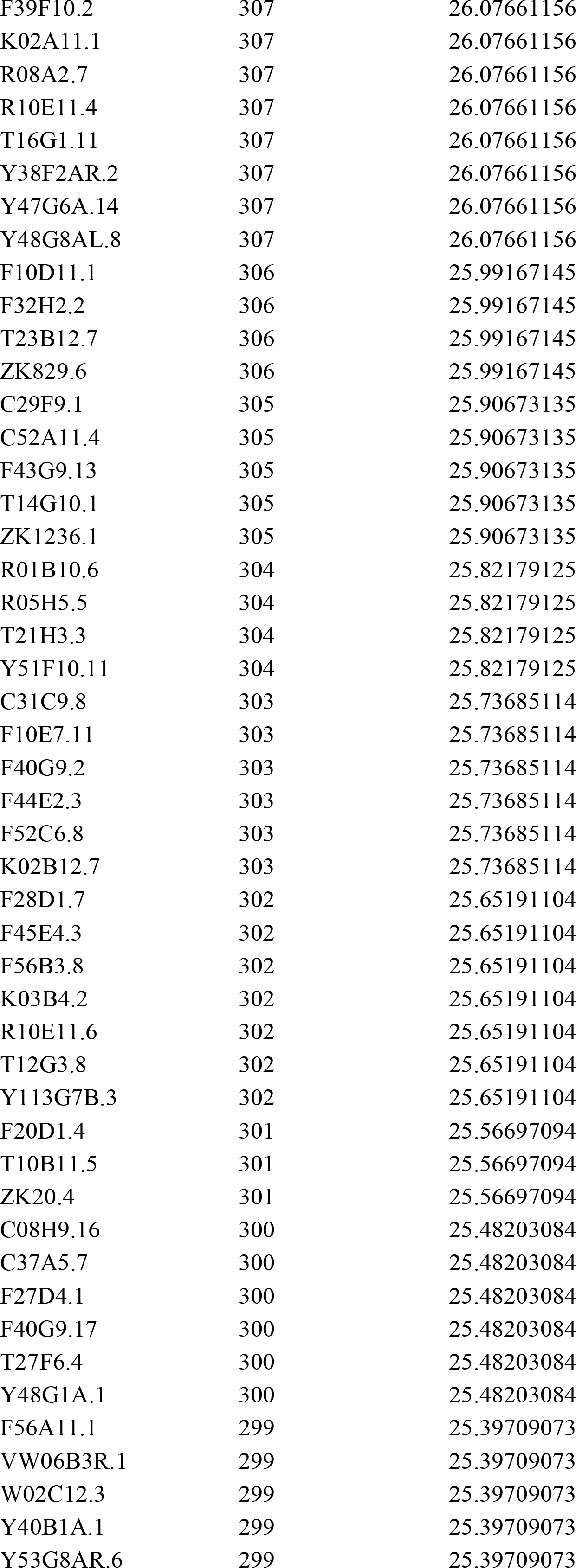
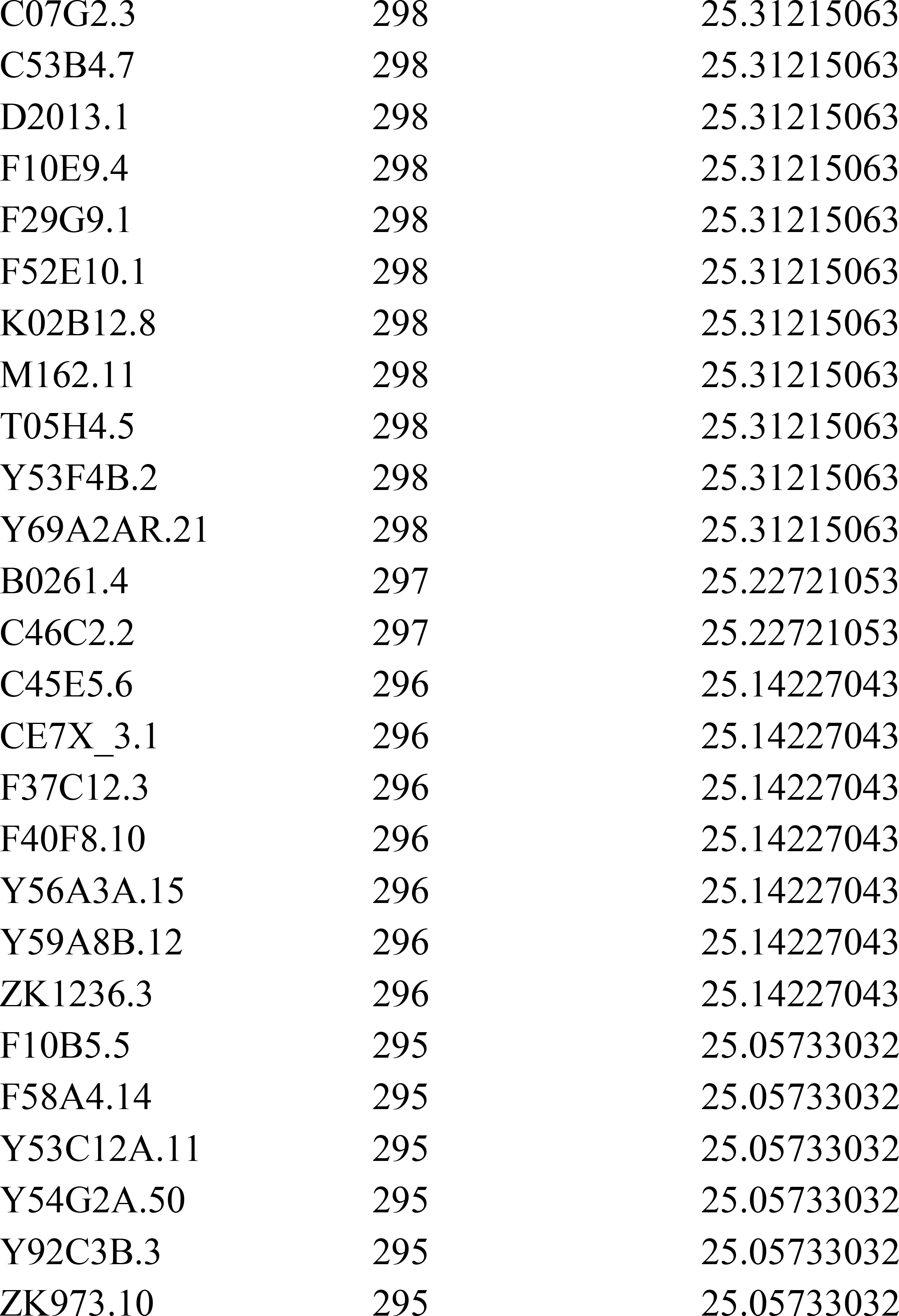
WAGO-4-associated siRNAs targeted genes, Related to Figure 5 The targets were selected using a threshold of 25 reads per million total reads.

